# Nance-Horan Syndrome-like 1 protein negatively regulates Scar/WAVE-Arp2/3 activity and inhibits lamellipodia stability and cell migration

**DOI:** 10.1101/2020.05.11.083030

**Authors:** Ah-Lai Law, Shamsinar Jalal, Tommy Pallett, Fuad Mosis, Ahmad Guni, Simon Brayford, Lawrence Yolland, Stefania Marcotti, James A. Levitt, Simon P. Poland, Maia Rowe-Sampson, Anett Jandke, Robert Köchl, Giordano Pula, Simon M. Ameer-Beg, Brian Marc Stramer, Matthias Krause

**Affiliations:** King’s College London, Krause group, Randall Centre for Cell and Molecular Biophysics, New Hunt’s House, Guy’s Campus, London, SE1 1UL, UK; King’s College London, Stramer group, Randall Centre for Cell and Molecular Biophysics, New Hunt’s House, Guy’s Campus, London, SE1 1UL, UK; King’s College London, Ameer-Beg group, Richard Dimbleby Cancer Research Laboratories, Comprehensive Cancer Centre, School of Cancer and Pharmaceutical Sciences, New Hunt’s House, Guy’s Campus, London, SE1 1UL, UK; King’s College London, School of Immunology and Microbial Sciences, Guy’s Campus, London, SE1 1UL, UK; The Francis Crick Institute, Immunosurveillance Laboratory, 1 Midland Road, London, NW1 1AT, UK; University Medical Center Hamburg (UKE), Institute for Clinical Chemistry and Laboratory Medicine, Martinistrasse 52, O26, D-20246 Hamburg, Germany; University of Bedfordshire. School of Life Sciences. Luton, LU1 3JU

**Keywords:** NHSL1, Nance Horan Syndrome-like 1 protein, Scar/WAVE complex, cell migration

## Abstract

Cell migration is important for development and its aberrant regulation contributes to many diseases. The Scar/WAVE complex is essential for Arp2/3 mediated lamellipodia formation during mesenchymal cell migration and several coinciding signals activate it. However, so far, no direct negative regulators are known. We have identified Nance-Horan Syndrome-like 1 protein (NHSL1) as a novel, direct binding partner of the Scar/WAVE complex, which co-localise at protruding lamellipodia. This interaction is mediated by the Abi SH3 domain and two binding sites in NHSL1. Furthermore, active Rac binds to NHSL1 at two regions that mediate leading edge targeting of NHSL1. Surprisingly, NHSL1 inhibits cell migration through its interaction with the Scar/WAVE complex. Mechanistically, NHSL1 may reduce cell migration efficiency by impeding Arp2/3 activity, as measured in cells using a novel Arp2/3 FRET-FLIM biosensor, resulting in reduced F-actin density of lamellipodia, and consequently impairing the stability of lamellipodia protrusions.

## Introduction

Cell migration is essential for embryonic development, homeostasis, and wound healing and its deregulation causes developmental defects, impaired wound healing, and cancer metastasis^1^. Mesenchymal cell migration depends on the polymerisation of actin filaments, which push the plasma membrane forward to form membrane extensions such as lamellipodia. Establishment of adhesions underneath lamellipodia stabilise protrusions allowing cells to advance^2^. Lamellipodium formation is controlled by Rac, a small GTPase of the Rho family. Active Rac, tyrosine phosphorylation and binding to phosphoinositides activates the Scar/WAVE complex at the very edge of lamellipodia and thereby recruits the Arp2/3 complex to nucleate branched actin networks. The Scar/WAVE complex is a heteropentameric, autoinhibited protein complex, consisting of Scar/WAVE1-3, Nap1, Sra1/Pir121, Abi1-3, and HSPC300^2–4^. We have previously shown that Lamellipodin (Lpd) localises to the very edge of lamellipodia^5^ and directly binds active Rac and the Scar/WAVE complex^6^. Rac increases the interaction between Lpd and the Scar/WAVE complex^6^. Lpd functions to promote cell migration via the Scar/WAVE complex^6,7^, which is consistent with a positive role for the Scar/WAVE complex in enhancing migration^8–11^. We postulate that the Scar/WAVE complex needs also to be negatively regulated at the leading edge to tightly control lamellipodial protrusion dynamics and cell steering. However, so far, it is not known how the Scar/WAVE complex is directly inhibited at the leading edge.

Here, we identify NHSL1 (Nance-Horan Syndrome-like 1) protein as a negative regulator of cell migration and we found that this is mediated by its interaction with the Scar/WAVE complex. NHSL1 belongs to the poorly investigated Nance-Horan Syndrome protein family along with Nance-Horan Syndrome (NHS) and NHSL2 proteins. Mutations in the NHS gene cause Nance-Horan syndrome, which is characterised by dental abnormalities, developmental delay, and congenital cataracts^12–14^. We show that NHSL1 directly binds to the Scar/WAVE complex and co-localises with it at the very edge of protruding lamellipodia. We found that active Rac binds to NHSL1 at two regions of the protein, which mediate leading edge targeting of NHSL1. The negative regulatory function of NHSL1 in cell migration may be due to its role in lamellipodia since we found that it reduces lamellipodia stability. NHSL1 acts to reduce Arp2/3 activity, which is consistent with our finding that NHSL1 reduces F-actin content of lamellipodia via its interaction with the Scar/WAVE complex. In NHSL1 CRISPR knockout cells we observed a reduction in total cellular Scar/WAVE2, Arp2/3 complex and F-actin levels indicating the permanent overactivation of Scar/WAVE and Arp2/3 complexes caused by loss of NHSL1 may lead to their degradation. Taken together, our data suggest that NHSL1 negatively regulates the Scar/WAVE complex, and hence reduces Arp2/3 activity, to control lamellipodia stability and consequently cell migration efficiency.

## Results

### NHSL1 localises to the very edge of lamellipodia

The Nance-Horan Syndrome (NHS) protein family consists of the Nance-Horan Syndrome (NHS) protein, Nance-Horan Syndrome-like 1 (NHSL1) protein and Nance-Horan Syndrome-like 2 (NHSL2) protein^12–16^. Expression analysis of NHSL1 by northern blot revealed a 7.5 kb ubiquitously expressed isoform and several shorter isoforms with a tissue specific distribution (Fig. 1A). In contrast, NHS expression was detected in brain, kidney, lung, and thymus^13^ suggesting that the NHS protein has evolved a more specialised function. We generated a rabbit polyclonal anti-NHSL1 antiserum (4457) (Fig. 1E) and a monoclonal antibody (clone C286F5E1). Both NHSL1 antibodies detect endogenous NHSL1 as a 260 kDa protein by immunoblotting in the melanoma cell line, B16-F1, and the breast epithelial cell line, MCF10A (Fig. 1B-D; S1A-C). B16-F1 cells display large lamellipodia when plated on 2D substrates and have been used extensively in other studies to characterise lamellipodia dynamics and cell migration^17^ and are thus used throughout this study.

**Figure 1.**
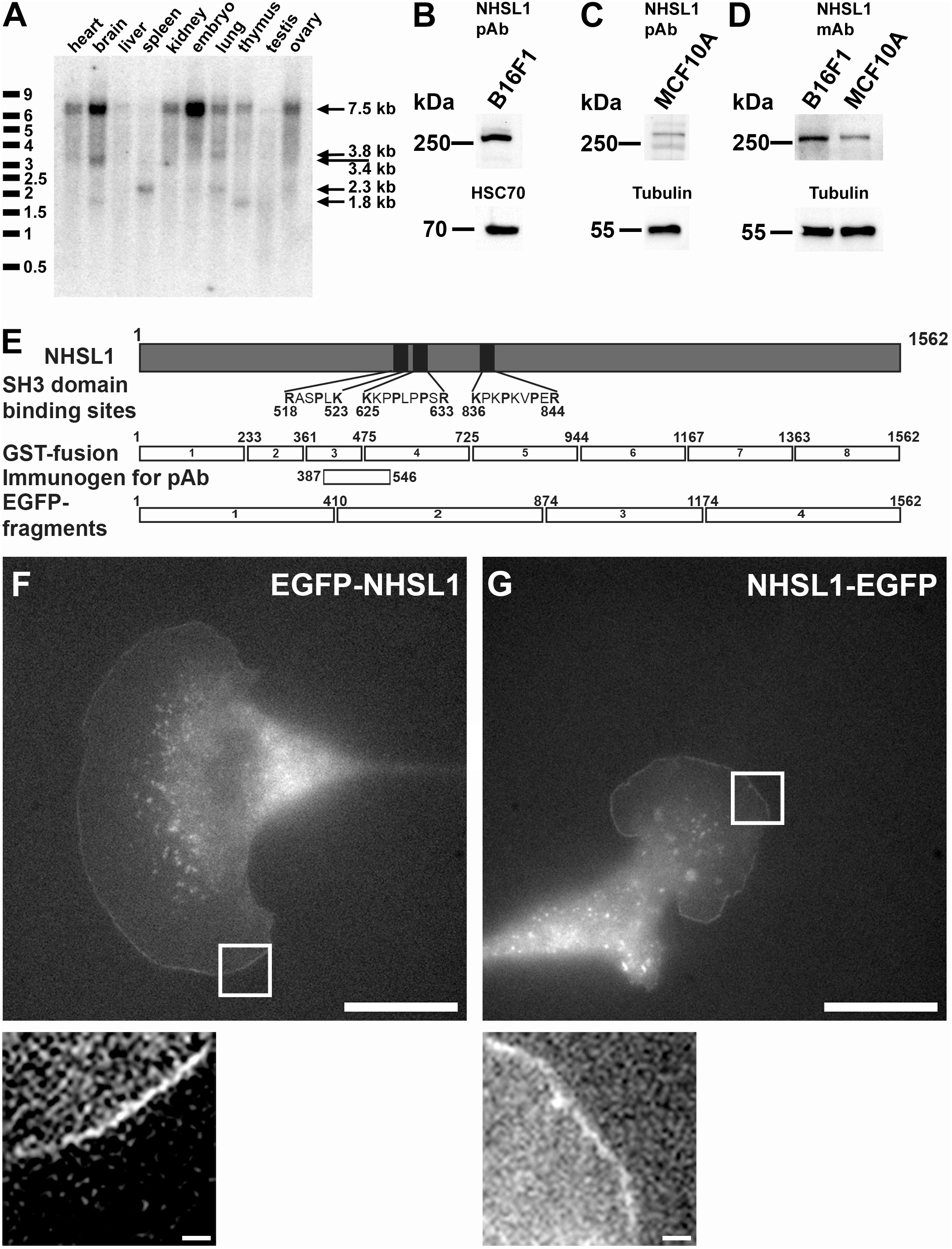
NHSL1 localises to the very edge of lamellipodia. **(A)** Northern blot analysis of NHSL1 expression in different tissues. Arrows show a large 7.5kb isoform that is ubiquitously expressed and various shorter isoforms with varying tissue specific distribution. **(B-D**) Western blots detecting NHSL1 protein in indicated cell lines using the (B,C) polyclonal (4457) or (D) monoclonal antibody (C286F5E1). For full blots see Fig. S1A-C. **(E)** Scheme of NHSL1 protein (grey rectangle) containing three putative SH3 binding sites (indicated black boxes). GST-fusion proteins 1 to 8 and EGFP-fusion proteins 1 to 4 of NHSL1 are indicated (white rectangles). Immunogen for the NHSL1 polyclonal antibody is indicated. **(F** and **G)** N-terminal EGFP-tagged NHSL1 (F) and C-terminal EGFP-tagged NHSL1 (G) were expressed in B16-F1 cells and, after plating on laminin, localisation of both EGFP-tagged NHSL1 constructs were imaged live. Representative images shown from three independent experiments. Scale bar: 20 μm. Inset represents a magnified view of the white box. Scale bar in inset: 1.25 μm. See also related video S1.

NHS localises to lamellipodia but the function of NHS and NHS-like proteins in cell migration are not known^14^. We found that NHSL1 also localises to the very edge of protruding lamellipodia and, in addition, to vesicular structures when tagged with EGFP either N- or C-terminally (Fig. 1F,G; video S1). This localisation is not due to space filling as B16-F1 cells display very thin lamellipodia and subtraction of the intensity of co-expressed cytoplasmic mScarlet does not remove the leading-edge signal of EGFP-NHSL1 (Fig. S1D; video S2). We previously showed that Lpd also localises at the very edge of lamellipodia^5^. When we co-expressed NHSL1-EGFP with mScarlet-Lpd in B16-F1 cells we found that both co-localise at the edge of lamellipodia (Fig. S1E; video S3). This prompted us to test whether both proteins can form a complex in cells. We tested this by co-expressing EGFP-Lpd with Myc-NHSL1 in HEK cells but could not detect an interaction after pulling down EGFP-Lpd with GFP-trap beads (Fig. S1F).

### NHSL1 negatively regulates cell migration speed and persistence

To explore the role of NHSL1 in cell migration we created stable NHSL1 CRISPR knockout B16-F1 cell lines by knocking in a stop codon into exon 2 (Fig. S2A), which is common to all isoforms. Clonal cell lines were tested by western blot and qPCR analysis, which revealed that NHSL1 expression was greatly reduced in clone CRISPR 2 but less reduced in clone CRISPR 21 (Fig. 2A,B S3A,E). The partially reduced mRNA and protein levels (Fig. 2A,B; S3A,E) in the CRISPR 21 cell line are consistent with Cas9 induced indels only in some alleles (Fig S2A), whereas the absence of NHSL1 wild type alleles (Fig S2B) and mRNA in the CRISPR 2 cell line (Fig. S3E) indicates that the CRISPR 2 line represents a full NHSL1 knockout.

**Figure 2.**
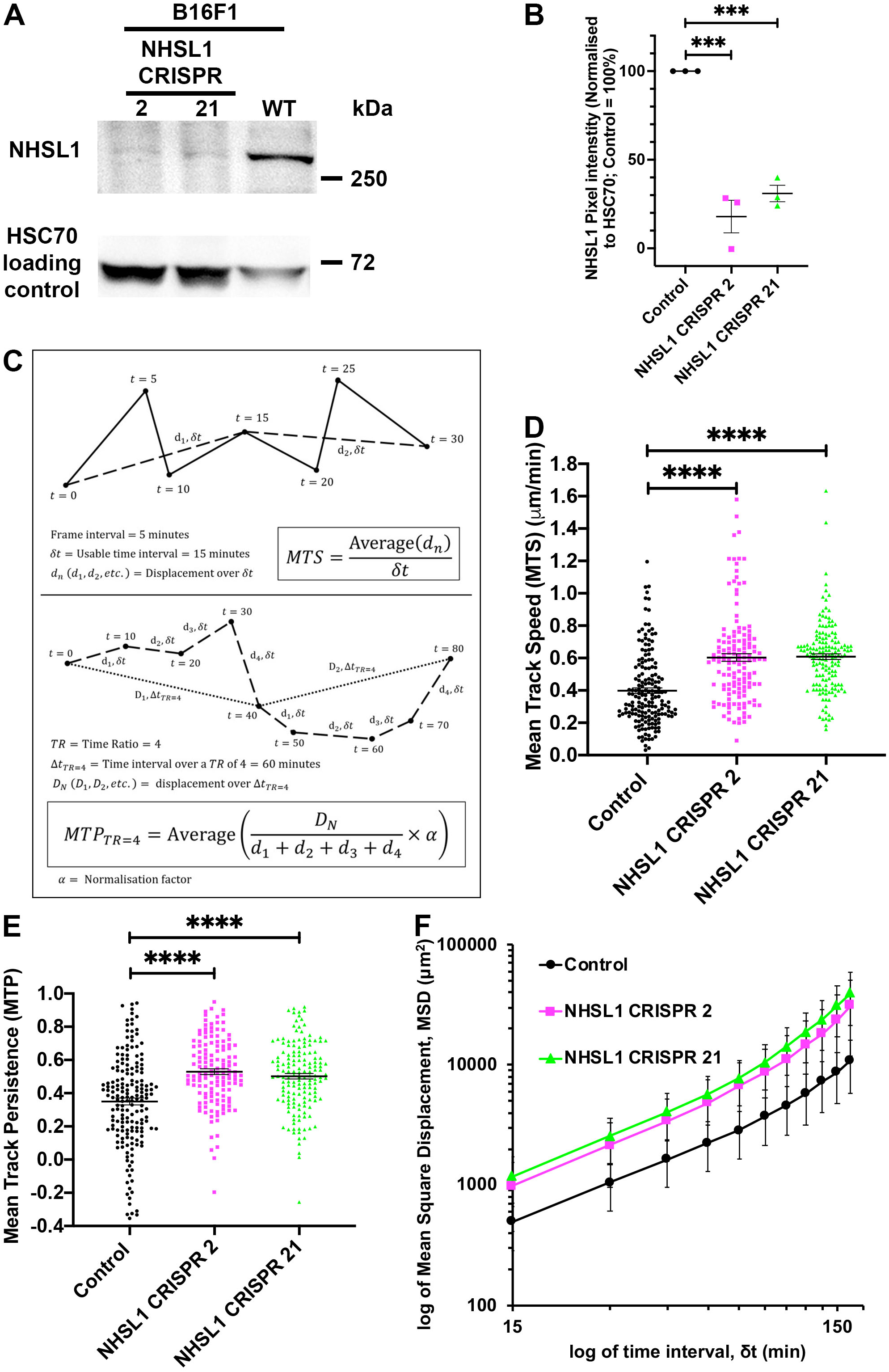
NHSL1 negatively regulates cell migration and persistence. **(A)** Western blot showing extent of reduction of NHSL1 expression in the clonal NHSL1 CRISPR B16-F1 cell lines 2 and 21 was probed with polyclonal NHSL1 antibodies and HSC70 as a loading control. Representative blot from three independent biological repeats. **(B)** Quantification of western blots in (A) normalised to the HSC70 loading control; One-way ANOVA: P<0.0001; F(2,6)=54.85; Dunnett’s multiple comparisons test: control vs. CRISPR 2: *** p=0.0001; control vs. CRISPR 21: *** p=0.0003. **(C)** Schematic diagram of the definition of Mean Track Speed (MTS) and Mean Track Persistence (MTP). **(D)** Cell migration speed of both NHSL1 CRISPR clones 2 and 21 in random cell migration assays was significantly increased by an average of approximately 52% compared to wild type control B16-F1 cells; Mean Track Speed (MTS) (dt = 3, TR = 4; see materials and methods for calculation). One-way ANOVA: P<0.0001; F(2,469)=41.87; Dunnett’s multiple comparisons test: Results are mean (μm/min) +/− SEM (error bars): control: 0.398 ± 0.017 μm/min; CRISPR 2: 0.603 ± 0.024 μm/min; control vs. CRISPR 2: **** p<0.0001; CRISPR 21: 0.609 ± 0.017 μm/min; control vs. CRISPR 21: **** p<0.0001. **(E)** Cell migration persistence was significantly increased by 53% or 51% for the NHSL1 CRISPR 2 or 21 cell line, respectively. Mean Track Persistence (MTP) (dt = 3, TR = 4; see materials and methods for calculation). One-way ANOVA: p<0.0001; F(2,469)=28.01; Dunnett’s multiple comparisons test: control: 0.350±0.021; CRISPR 2: 0.529±0.017; control vs. CRISPR 2: **** p<0.0001; CRISPR 21: 0.502±0.016; control vs. CRISPR 21: **** p<0.0001; Results are mean (this is dimensionless see Fig. 2C for explanation of MTP) +/− SEM (error bars); (D,E) data points are from four independent biological repeats with a total cell number of 177, 140 and 156 for wild type, NHSL1 clones 2 and 21, respectively. **(F)** log-log plot of Mean Square Displacement (MSD) of control B16-F1 cells, NHSL1 CRISPR clones 2 and 21 in random cell migration assays from (D,E). Source data are provided as a Source Data file.

In random cell migration assays on fibronectin, cell speeds significantly increased by 52% for both NHSL1 CRISPR clones compared to wild-type B16-F1 cells (Fig. 2C,D; S3B-D). Traditionally, cell migration persistence is measured as the directionality ratio: the ratio of the straight-line distance and the total trajectory path length of each track. This leads to inaccurate measurements as this ratio is affected by different track lengths as well as cell speeds^19^. Instead we quantified cell persistence using the Dunn persistence method^6^ by calculating the directionality ratio over a short interval of the movie, defined by a time ratio, *TR*, and then averaging over all these intervals comprising the whole track to obtain the “Mean Track Persistence” (MTP). This analysis revealed that cell migration persistence was also significantly increased by 53% and 51% for the CRISPR 2 and CRISPR 21 cell lines, respectively (Fig. 2C,E). This increase in persistence could also be seen when we calculated the directionality ratio over time (Fig. S4A) and the direction autocorrelation, a measure of how the angle of displacement vectors correlate with themselves which is independent of speed^19,20^ (Fig. S4B). Analysing mean square displacement also indicated that NHSL1 CRISPR 2 and CRISPR 21 cells explore a larger area (Fig. 2F). In agreement, we found that directional migration into a scratch wound was significantly increased in NHSL1 knockdown MCF10A (Fig. S5A-C, video S4). Together these results suggest that NHSL1 negatively regulates cell migration speed and persistence.

### NHSL1 is a novel binding partner of active Rac

To characterise the region of NHSL1 required for leading edge recruitment (Fig. 1F,G), four EGFP-tagged fragments covering the entire length of NHSL1 (Fig. 3A) were expressed in B16-F1 cells. Live cell imaging revealed that only fragments 2 and 3 localised to lamellipodia (Fig. 3D,E; video S5) In addition, fragments 1 and 3 were detected at vesicular structures (Fig. 3C,E; video S5).

**Figure 3.**
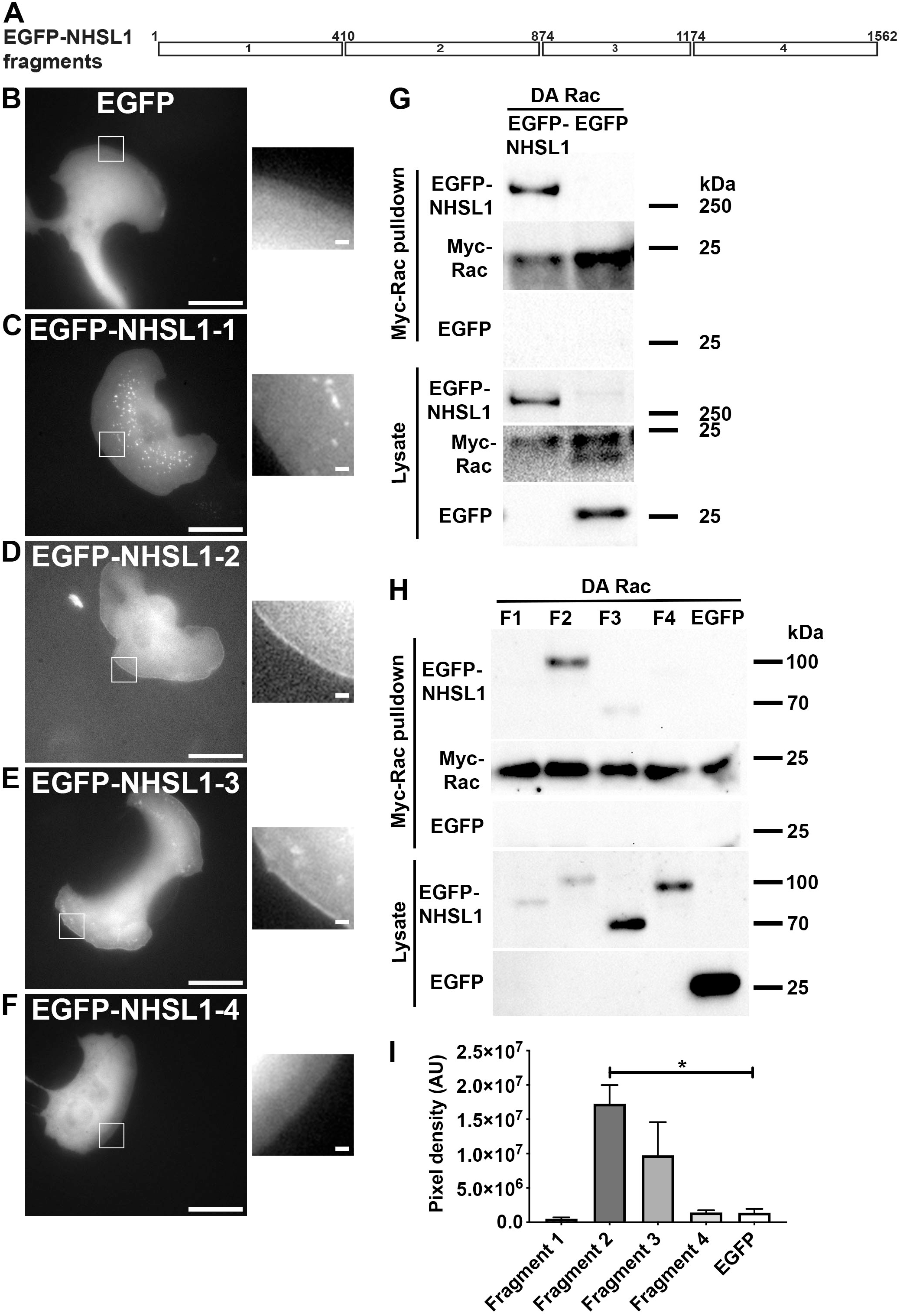
NHSL1 is a novel binding partner of active Rac. **(A)** Scheme of NHSL1 protein: EGFP-fusion proteins 1 to 4 of NHSL1 are indicated (white rectangles). **(B-F)** Four EGFP-tagged fragments covering the entire length of NHSL1 (see Fig. 1E and 3A) were generated and expressed in B16-F1 cells **(C-F)** along with EGFP as a negative control **(B).** Insets show enlarged pictures of the boxed areas from stills taken from live cell imaging. Fragments 2 and 3 localise to the very edge of lamellipodia similar to full length NHSL1. Representative images from three independent experiments. Scale bar: 20 μm. Inset represents a magnified view of the white box. Scale bar in inset: 1.25 μm. See also related video S5. **(G)** Western blot showing that dominant active (DA) Rac pulls down NHSL1 using Myc-trap beads from HEK cell lysates expressing Myc-tagged DA Rac co-expressed with EGFP-tagged NHSL1 or EGFP only as control. Representative blots from three independent biological repeats (see Fig. S6A for full western blots). **(H)** Western blot showing Myc-tagged DA Rac pulldowns of the four NHSL1 fragments (see Fig. 3A and Fig. 3B-E) from HEK cells expressing the EGFP-tagged fragments covering the entire length of NHSL1 or EGFP only as control co-expressed with Myc-tagged DA Rac. Representative blots from three independent experiments. **(I)** Quantification of band intensity of chemiluminescence from (H) imaged with a charge-coupled device (CCD) camera shows that Myc-tagged DA Rac binds significantly to fragment 2 compared to GFP. The amount of EGFP-NHSL1 fragments were normalised to the amount of Myc-Rac that was pulled down. Bars indicate mean ± SEM (error bars), n = 3. One-way ANOVA: p=0.00108, F(5,12)= 4.963; and Dunnett’s multiple comparisons test: *, p=0.0159. Source data are provided as a Source Data file.

Since Rac is an important regulator of lamellipodia formation, we hypothesised that Rac may bind NHSL1 for its recruitment to lamellipodia. We therefore tested for interaction between dominant active (DA) Myc-tagged Rac1-Q61L and EGFP-tagged NHSL1 or EGFP only as control in HEK cells. Myc-tagged DA Rac1 was pulled down with Myc-trap beads and interaction with EGFP-NHSL1 was evaluated by western blot against EGFP. This analysis revealed that NHSL1 is in complex with active Rac (Fig. 3G; S6A). This was verified in a reciprocal experiment in which EGFP-Rac1-Q61L or EGFP was pulled down with GFP-trap beads and the interaction with Myc-NHSL1 evaluated by western blot against Myc (Fig. S6B). Again, we observed an interaction between dominant active Rac and NHSL1 suggesting that NHSL1 is a novel binding partner of active Rac.

We again expressed the four EGFP-tagged NHSL1 fragments to determine which fragment may mediate the interaction with Myc-tagged dominant active Rac1 and pulled down Myc-DA-Rac1 with Myc-trap beads. A western blot against EGFP showed robust pulldown of EGFP-NHSL1 fragment 2 and weak pulldown of fragment 3 by active Rac (Fig. 3H,I). To further delineate the Rac binding sites in NHSL1, we generated nine overlapping sub-fragments covering fragments 2 and 3 (Fig. S7A). Again, Myc-trap pulldown of Myc-DA-Rac1 from HEK lysates co-expressing EGFP-tagged NHSL1 sub-fragments 1 to 9 revealed that sub-fragments 1 and 6, which overlap with each other at the N-terminus of fragment 2, and sub-fragment 9, which is centrally located in fragment 3, are in complex with active Rac (Fig. S7B).

Taken together, our data suggest that NHSL1 may be recruited to the leading edge by membrane associated active Rac binding directly or indirectly to two regions in NHSL1 located at amino acid 410-563 in fragment 2 and at amino acid 946-1099 in fragment 3.

### NHSL1 interacts with the Scar/WAVE complex via Abi

As the Scar/WAVE complex localises to the leading edge of cells^2^, we tested whether NHSL1 co-localises with it at this site. Using live cell imaging, we observed that NHSL1-EGFP co-localises with mScarlet-tagged Abi1 and Nap1, two components of the Scar/WAVE complex at the very edge of protruding lamellipodia (Fig. 4A,B; video S6). In agreement, we also found that endogenous NHSL1 co-localises with Abi1, Scar/WAVE1 and Scar/WAVE2 at this site (Fig. 4C-E; see line scans in Fig. S8A-H).

**Figure 4.**
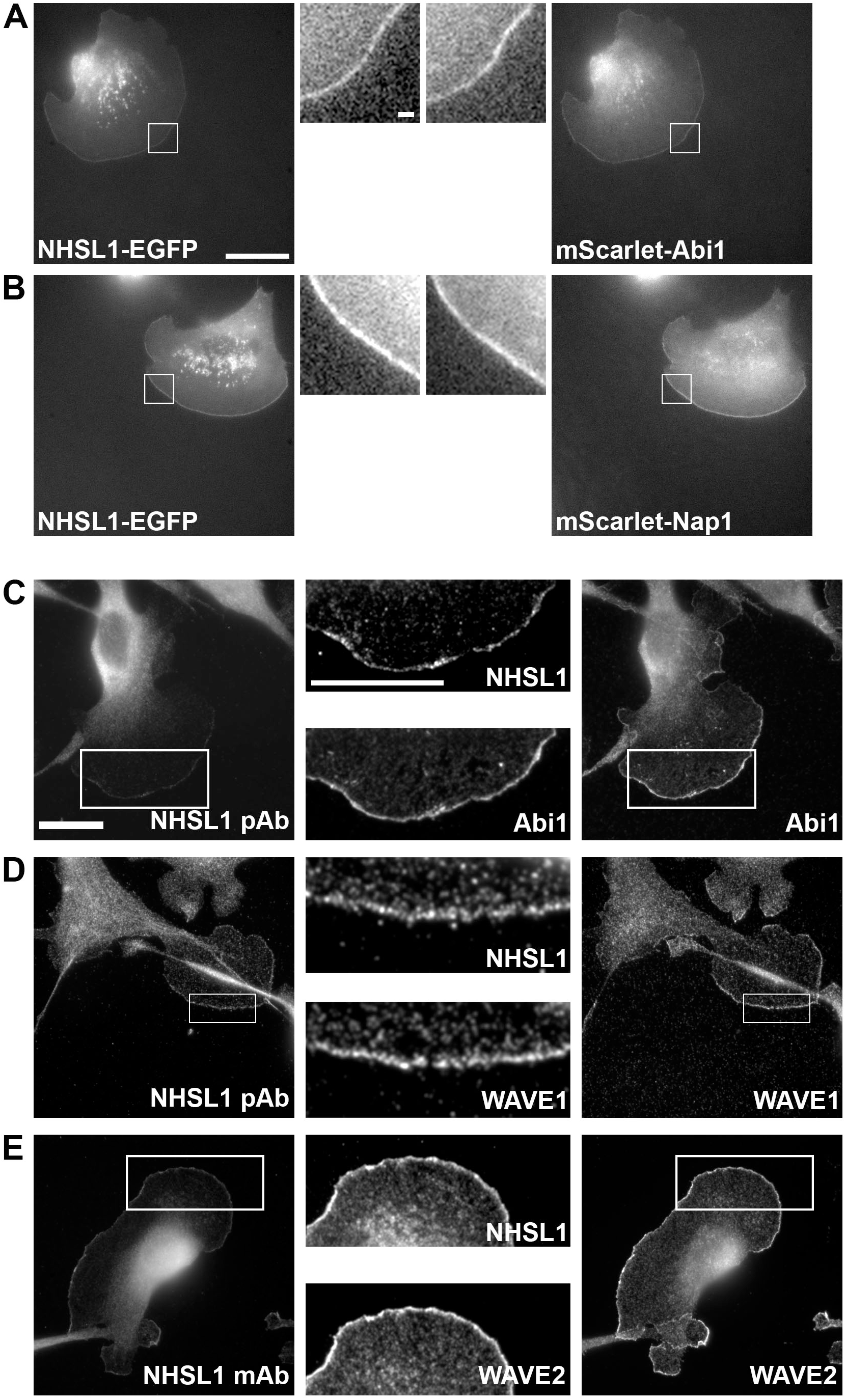
NHSL1 co-localises with Abi and the Scar/WAVE complex. **(A)** Still images from live cell imaging showing NHSL1-EGFP co-expressed with mScarlet-I-tagged Abi1 and **(B)** Nap1 in B16-F1 cells. Representative images shown from three independent experiments. Scale bar represent 20 µm. Inset is a magnified view of the white box. Scale bar in inset (A) applies to both insets (A,B): 1.25 μm. See also related video S6. **(C-E)** Endogenous NHSL1 co-localises with Abi1 (C, NHSL1 pAb) Scar/WAVE1 (D, NHSL1 pAb), and Scar/WAVE2 (E, NHSL1 mAb) at the very edge of lamellipodia in B16-F1 mouse melanoma cells. Scale bar in (C) applies also to (D,E): 20 µm. Inset represents a magnified view of the white box. Scale bar in inset in (C) applies also to inset for (D,E): 20 μm. Representative images shown from three independent experiments. See Fig. S8 for line scans of co-localisations in (C) and (E).

To explore a potential interaction with the Scar/WAVE complex, Myc-tagged components of the Scar/WAVE complex and EGFP-NHSL1 were expressed in HEK cells. Immunoprecipitation of EGFP-NHSL1 revealed that NHSL1 interacts with the Scar/WAVE complex (Fig. 5A; S9A,B). To confirm whether the respective endogenous proteins can be found in a complex in cells we immunoprecipitated endogenous NHSL1 and observed co-immunoprecipitation with Abi1 and Scar/WAVE2 from B16-F1 and MCF10A cell lysates (Fig. 5B,C).

**Figure 5.**
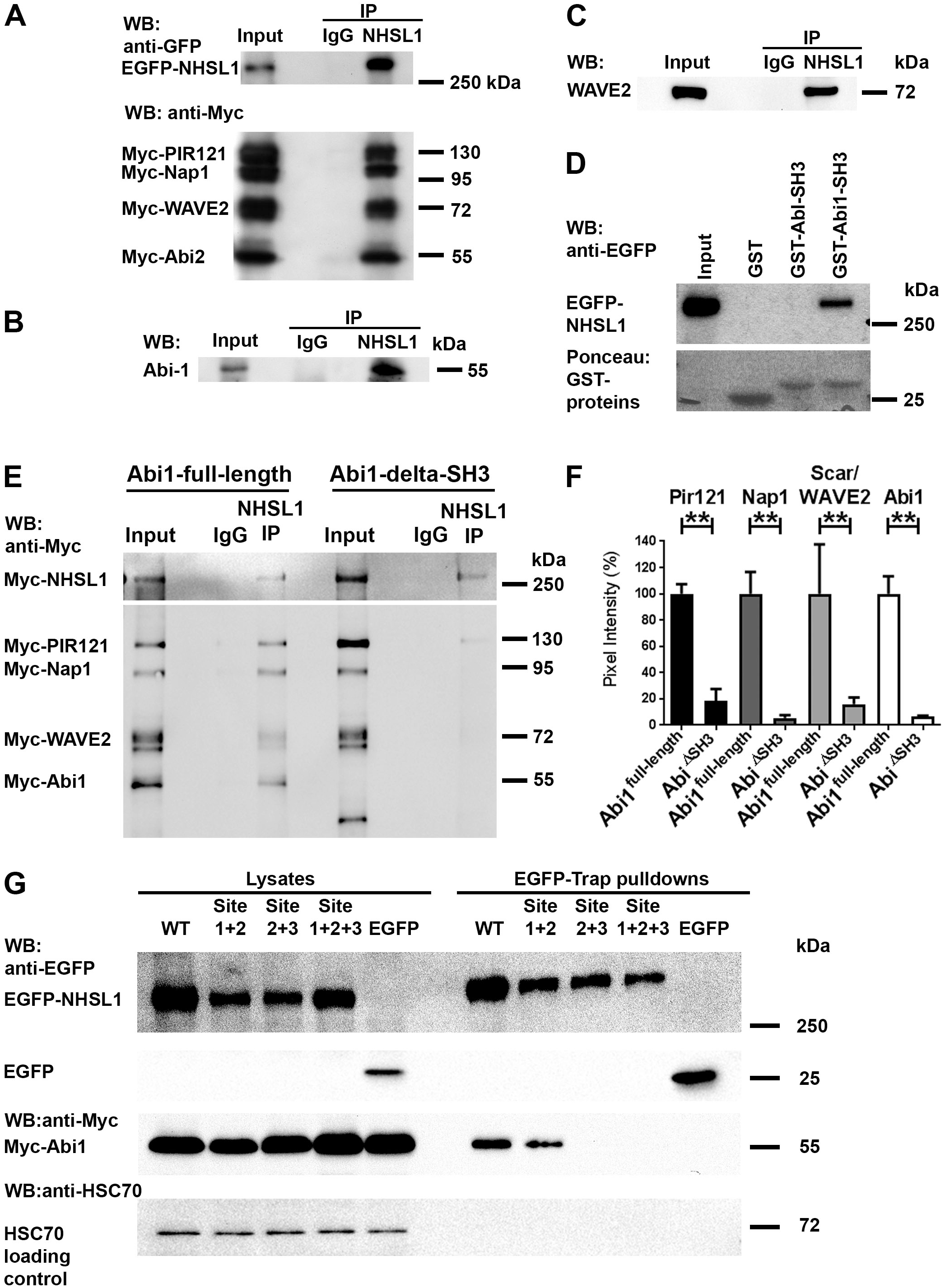
NHSL1 interacts with Abi and the Scar/WAVE complex. **(A)** The Scar/WAVE complex co-immunoprecipitates with NHSL1. HEK cells were transfected with EGFP-NHSL1, and Myc-Pir121, -Nap-1, -WAVE2, -Abi2. NHSL-1 was immunoprecipitated (pAb 4457) from lysates and co-immunoprecipitation tested on a western blot with Myc and EGFP antibodies. Representative blots from three independent experiments (see Fig. S9A,B for full western blots). **(B)** and **(C)** Western blot showing endogenous NHSL1 pulled down with polyclonal NHSL1 (4457) antibody followed by western blotting with (B) Abi1 and (C) Scar/WAVE2 antibodies detecting endogenous co-immunoprecipitation of these proteins in MCF10A (Abi1) and B16-F1 (Scar/WAVE2) cell lysates. Immunoprecipitation using non-immune rabbit IgG served as a negative control. Representative blots from three independent experiments. **(D)** GST-pull downs using purified Glutathione-sepharose coupled GST-fusion proteins of Abi1 and c-Abl SH3 domains or GST alone from HEK cell lysates that were transfected with EGFP-NHSL1. Following GST-pulldown EGFP-NHSL1 was detected in a western blot with anti-EGFP antibodies. Ponceau staining of membrane reveals GST or GST-tagged Abi- or Abl-SH3 domains used. Representative blots from three independent experiments. **(E)** Western blot showing immunoprecipitation using NHSL1 polyclonal (4457) antibody or non-immune rabbit IgG control from HEK cell lysates expressing Myc-NHSL1 and all Myc-tagged components of the Scar/WAVE complex, including Myc-Abi1-full-length (E, left), Myc-Abi1-delta-SH3 (E, right). The western blot shows co-immunoprecipitation between NHSL1 and all components of the Scar/WAVE complex only when the Abi SH3 domain is present (Myc-HSPC300 is not shown). Representative blots from three independent experiments. **(F)** Quantification of band intensity from chemiluminescence from (E) imaged with a CCD camera. Co-immunoprecipitation is reduced by >80%. Bars indicate mean ± SEM (error bars); n = 3; One-way ANOVA: p=0.0002; F(7,16)=8.855; Sidak’s multiple comparisons test: PIR121 **, p=0.0093; Nap1 **, p=0.0028; WAVE2 **, p=0.0074; Abi1 **, p=0.0030. **(G)** HEK cells were transfected with EGFP-tagged NHSL1 or NHSL1-SH3 binding mutants or EGFP only as negative control and Myc-Abi1. After GFP-trap pulldown of wild type (WT) NHSL1 or different NHSL1-SH3 binding mutants co-precipitation was detected in a western blot with Myc antibody. Representative blots from five independent experiments. Source data are provided as a Source Data file.

Since NHSL1 contains several putative SH3 binding sites we hypothesised that the SH3 domain of Abi, which is part of the Scar/WAVE complex, may mediate the interaction with NHSL1. The interaction between NHSL1 and Abi was tested by GST-pulldown experiments with purified Abi1-SH3 and c-Abl-SH3 from HEK lysates containing EGFP-NHSL1 or EGFP only as control. c-Abl SH3 was used as another control since binding to Abi may be mediated indirectly by binding with the Abi ligand c-Abl which also contains a SH3 domain^21,22^. We found that while the EGFP negative control did not bind (Fig. S9C), EGFP-NHSL1 specifically bound to the SH3 domain of Abi and not to the SH3 domain of c-Abl (Fig. 5D).

We next explored whether the SH3 domain of Abi is necessary for the interaction between NHSL1 and the Scar/WAVE complex. NHSL1 was immunoprecipitated from HEK cell lysates containing Myc-NHSL1 and all five Myc-tagged components of the Scar/WAVE complex including either full-length Abi1 (Abi1-full-length) or an Abi1 construct that lacked the SH3 domain (Abi1-delta-SH3). Co-immunoprecipitation was detected with an antibody against the Myc-tag (Fig. 5E). Quantification of co-immunoprecipitation of NHSL1 with the Scar/WAVE complex revealed that expression of Abi1-delta-SH3 disrupted the interaction between NHSL1 and the Scar/WAVE complex (Fig. 5E,F). This suggests that the SH3 domain of Abi mediates the interaction of NHSL1 with the Scar/WAVE complex.

To verify that the Scar/WAVE complex properly forms in cells when all components of the Scar/WAVE complex are expressed as tagged proteins, we expressed Myc-tagged PIR121, Nap1, HSPC300, and Scar/WAVE2 together with either EGFP-Abi1-full-length, EGFP-Abi1-delta-SH3, or EGFP in HEK cells. After GFP-trap pulldown from cell lysates, we detected bound Myc-tagged proteins with an antibody against the Myc-tag. This analysis revealed that indeed all tagged components of the Scar/WAVE complex faithfully associate with each other as a complex (Fig. S9D).

To test whether the interaction between Abi and NHSL1 is direct and to map the SH3 binding site in NHSL1, we performed a far western blot experiment. We purified from *E.coli* eight GST-fusion proteins covering the entire length of NHSL1 (Fig. 1E, S10A,B), which were separated on SDS-PAGE, followed by blotting onto membrane. We overlaid this membrane with purified MBP-tagged full-length Abi1 (MBP-Abi1 full-length) or an MBP fusion protein with Abi1 in which the SH3 domain had been deleted (MBP-Abi1-delta-SH3) or MBP as control. The far western overlay showed that only fragments 4 and 5 of NHSL1 directly interacted with wild type Abi but neither with Abi missing the SH3 domain nor MBP on its own (Fig. S10A). In agreement, fragments 4 and 5 contain three putative SH3 binding sites suggesting that Abi binds directly via its SH3 domain to NHSL1.

Next, we explored whether these putative SH3 binding sites were sufficient for the interaction with Abi. We mutated SH3 binding sites 1 and 2 (site 1+2), or sites 2 and 3 (site 2+3) or all three sites together (site 1+2+3) in full-length NHSL1 and expressed the EGFP-tagged mutant and wild type cDNA’s together with Myc-tagged Abi1 in HEK cells. After GFP-trap pulldown from lysates, western blot against the Myc-tag revealed that only EGFP-NHSL1 (site 2+3) and NHSL1 (sites 1+2+3) showed loss of interaction with Abi1 (Fig.5G). Taken together, these data indicate that Abi binds via its SH3 domain to two sites in NHSL1.

### NHSL1 negatively regulates cell migration via the Scar/Wave complex

We observed that loss of NHSL1 resulted in increased cell migration speed and persistence (Fig. 2, S3, S4). To examine the consequences of increasing NHSL1 expression, we overexpressed EGFP-tagged wild type NHSL1 (EGFP-NHSL1 WT) or the NHSL1 cDNA which cannot interact with the Abi SH3 domain and hence cannot interact with the Scar/WAVE complex (Fig. 5G) (EGFP-NHSL1 SW Mut) in B16-F1 cells (Fig. S11A). We quantified random cell migration behaviour after plating the cells on fibronectin and observed a moderate but significant reduction in cell migration speed (Fig. 6A) and a moderately reduced mean square displacement (Fig. S11B) for cells overexpressing wild type EGFP-NHSL1 compared to EGFP control. This is consistent with the result from the NHSL1 CRISPR cells, which displayed the opposite effect (Fig. 2C-F). Cell migration persistence was increased upon overexpression of NHSL1 (Fig. S11C-E). Since CRISPR knockout of NHSL1 also increases persistence, this suggests that optimal expression levels of NHSL1 may finetune cell migration persistence.

**Figure 6.**
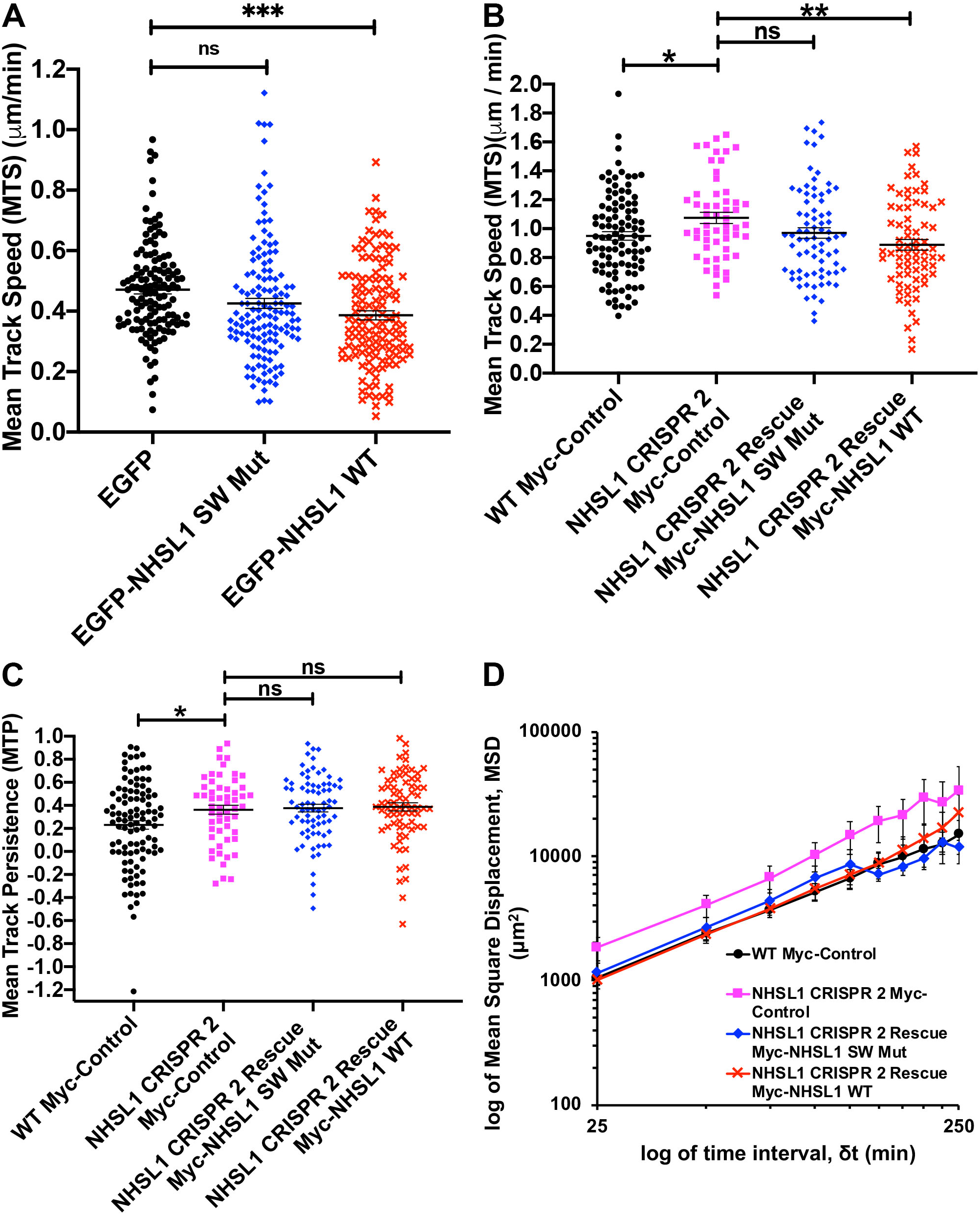
NHSL1 negatively regulates cell migration via the Scar/WAVE complex. **(A)** Quantification of speed of randomly migrating B16-F1 cells overexpressing either wild type NHSL1 (NHSL1 WT, black circles) or the NHSL1 mutant in the Scar/WAVE complex binding sites (NHSL1 SW Mut, blue diamonds) or EGFP alone (red crosses) as control plated on fibronectin after selection using a bicistronic expression plasmid also conferring resistance to puromycin to ensure that all cells analysed overexpressed NHSL1. Mean track speed (dt = 3, TR = 4; see materials and methods for calculation). Results are mean ± SEM (error bars), of four independent biological experiments. Each data point represents the mean speed of a cell from a total of number of 106, 104, 108 cells for control, NHSL1 WT, and NHSL1 SW Mut, respectively. One way ANOVA: p=0.0007; F(2,395)=7.385; and Dunnett’s multiple comparisons test: ***, p=0.0003; ns, p=0.0655; **(B, C)** Quantification of speed and persistence of randomly migrating wild type B16-F1 cells expressing Myc alone as control (black circles) or CRISPR 2 cells expressing either the NHSL1 mutant in the Scar/WAVE complex binding sites (NHSL1 SW Mut, blue diamonds) or NHSL1 (NHSL1 WT, red crosses) or Myc alone as control (pink squares) plated on laminin after selection using a bicistronic expression plasmid also conferring resistance to blasticidin to ensure that all cells analysed expressed NHSL1. Mean track speed (B) and persistence (C) (dt = 5, TR = 2; see materials and methods for calculation). Each data point represents the mean speed of a cell from a total of number of 102, 55, 77, 72 for wild-type B16-F1 expressing Myc only as control, CRISPR 2 cells expressing Myc only, CRISPR 2 Rescue Myc-NHSL1 SW Mut and CRISPR 2 Rescue Myc-NHSL1 WT, respectively. Results are mean ± SEM (error bars) from five independent biological experiments. (B) One-way ANOVA: p=0.0077; F(3,302)=4.044; and Dunnett’s multiple comparisons test: **, p=0.0018; *, p=0.0364; ns, p=0.1215; (C) One-way ANOVA: p=0.0034; F(3,302)=4.648; and Dunnett’s multiple comparisons test: CRISPR2 vs. WT control: *, p=0.0397; NHSL1 CRISPR2 vs. Rescue Myc-NHSL1 SW Mut: ns, p=0.9877; NHSL1 CRISPR2 vs. Rescue Myc-NHSL1 WT: ns, p=0.9442. **(D)** Mean Square Displacement analysis (log-log plot) of data shown in (B,C). Source data are provided as a Source Data file.

The NHSL1 Scar/WAVE binding mutant localises to the very edge of lamellipodia (Fig. S11F, video S7) like wild type EGFP-NHSL1 (Fig. 1F,G, video S1). In contrast, the NHSL1 Scar/WAVE binding mutant did not reduce migration speed (Fig. 6A) suggesting that NHSL1 negatively regulates cell migration speed via an interaction with the Scar/WAVE complex. To verify this and to test whether the observed phenotypes in the NHSL1 CRISPR knockout clones were not due to off-target effects, we re-expressed Myc-tagged wild type NHSL1 (Myc-NHSL1 WT) or the NHSL1 Scar/WAVE complex binding mutant (Myc-NHSL1 SW Mut) in B16-F1 cells. After plating the cells on laminin we again observed an increase in random cell migration speed and persistence between wild type control and NHSL1 CRISPR2 knockout cells (Fig. 6B,C) confirming our previous results (Fig. 2C-E). Re-expression of neither wild type Myc-NHSL1 nor Myc-NHSL1 SW Mut in CRISPR 2 cells rescued cell migration persistence (Fig. 6C,D; S12A,B). This result and the increase in cell migration persistence upon NHSL1 overexpression (Fig. S11C-E) and NHSL1 knockout (Fig. 2E) together suggest that optimal expression levels of NHSL1 may be required for the lower cell migration persistence observed in the wild type cells. However, re-expression of wild type Myc-NHSL1 but not Myc-NHSL1 SW Mut in CRISPR 2 cells resulted in a significant reduction in cell migration speed (Fig. 6B). This indicates that wild type NHSL1 but not NHSL1 SW Mut (that cannot bind to the Scar/WAVE complex) can rescue the NHSL1 knockout phenotype and suggests that NHSL1 negatively regulates cell migration speed via an interaction with the Scar/WAVE complex.

### NHSL1 CRISPR knockout increases Arp2/3 activity in cells

As the Scar/WAVE complex is the major activator of the Arp2/3 complex, which induces branched F-actin nucleation at the leading edge of cells, we sought a direct method to investigate whether NHSL1 controls Arp2/3 activity. In order to directly quantify Arp2/3 activity in cells, we developed a novel FRET-FLIM Arp2/3 biosensor. We designed this biosensor based on *in vitro* FRET probes for the Arp2/3 complex^23,24^ which were successfully used to measure the activity of the purified complex and relies on the conformational change of the Arp2/3 complex upon binding to nucleation promotion factors such as the Scar/WAVE complex^24–30^. Here we tagged the Arp2/3 subunits ARPC3 with mTurq2 as the FRET donor and ARPC1B with mVenus as the FRET acceptor and expressed them from a bicistronic construct (ARPC1B-mVenus-P2A-ARPC3-mTurq2). When the Arp2/3 complex is activated by Scar/WAVE, ARP2 and ARP3 and also ARPC3 and ARPC1B move closer to each other^23–30^ allowing FRET between mTurq2 and mVenus to occur. To evaluate whether this biosensor faithfully reports changes in Arp2/3 activity in cells, we expressed our Arp2/3 biosensor together with Myc-DA-Rac1 or empty Myc vector as control and measured FRET efficiency by Fluorescence Lifetime Imaging (FLIM). This analysis revealed that FRET efficiency and hence Arp2/3 activity was significantly increased upon expression of dominant active Rac compared to control (Fig. 7A-C; S13A-E). We then expressed our biosensor in control B16-F1 cells or in the NHSL1 CRISPR 2 and 21 cell lines and again measured FRET by FLIM. We found that FRET efficiency and hence Arp2/3 activity was significantly increased in both NHSL1 CRISPR 2 and 21 cell lines compared to control B16-F1 cells (Fig. 7D-F; S14A-F).

**Figure 7.**
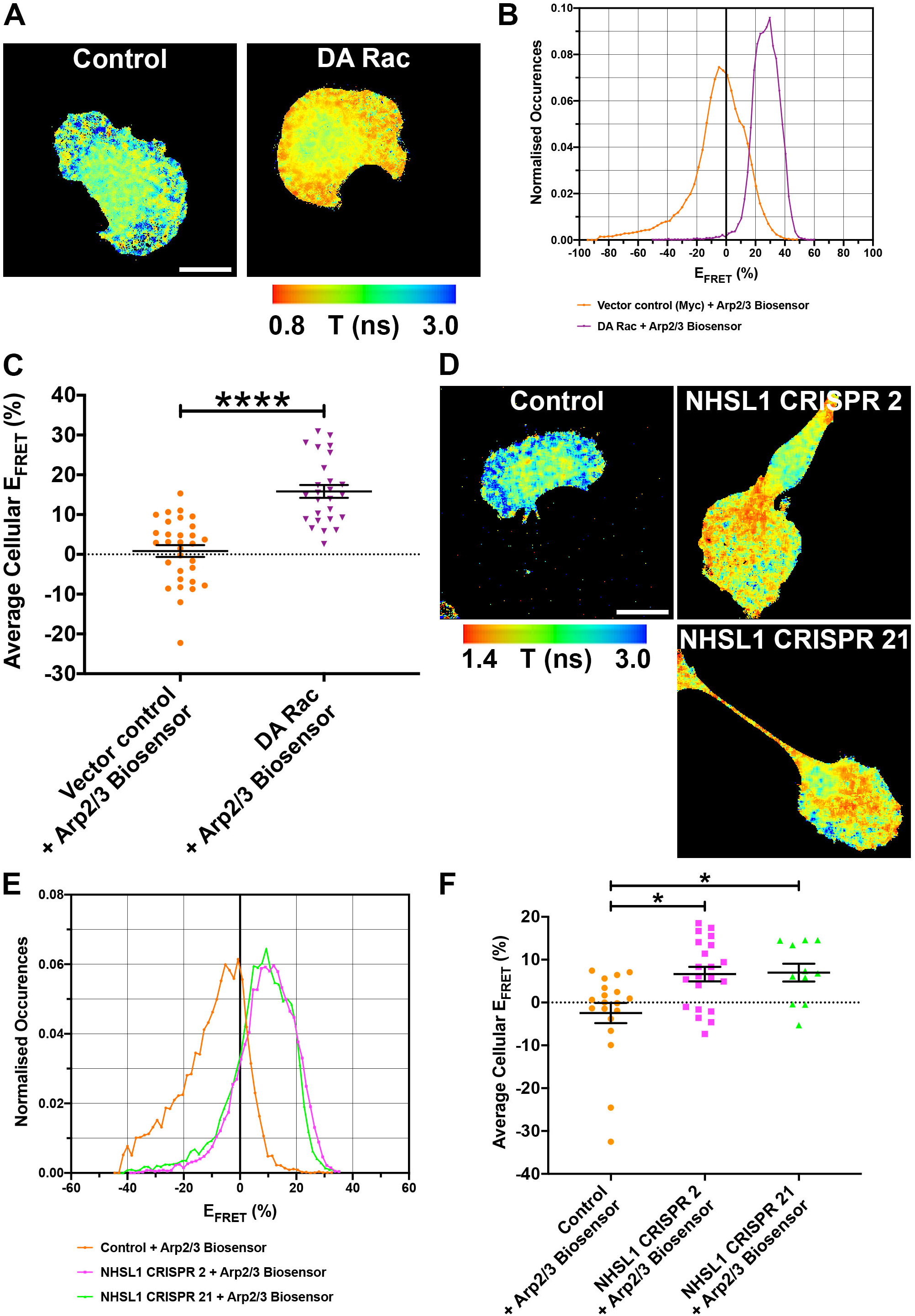
NHSL1 negatively regulates Arp2/3 activity. **(A)** Lifetime images of B16-F1 expressing the Arp2/3 biosensor and either empty Myc-control plasmid or Myc-tagged dominant active Rac (DA Rac). Warm colours indicate short lifetimes of the donor mTurq2 which represent high FRET efficiency and Arp2/3 activity. Representative images from 4 independent experiments are shown. Scale bar: 20 μm. **(B)** FRET efficiency histograms from the same representative wild type B16-F1 (orange circles) and DA Rac (lilac upside-down triangles) cells expressing the Arp2/3 biosensor as shown in (A). **(C)** Quantification of average cellular FRET efficiency which represents Arp2/3 activity from 4 independent biological repeats from control: 31 cells (orange circles); DA Rac: 26 cells (lilac upside-down triangles). Data points are the weighted average means for each cell, calculated from the FRET efficiency histograms and used instead of the normal (unweighted) mean in order to better represent the true FRET efficiency. Results are the average weighted average mean ± SEM (error bars), ****, p<0.0001; t=7.379; df=54. Unpaired, two-tailed t-test. **(D)** Lifetime images of wild-type B16-F1, NHSL1 CRISPR clone 2 and NHSL1 CRISPR clone 21 cells expressing the Arp2/3 biosensor. Warm colours indicate short lifetimes of the donor mTurq2 which represent high FRET efficiency and Arp2/3 activity. Representative images from 4 independent experiments are shown. Scale bar: 20 μm **(E)** FRET efficiency histograms from the same representative wild-type B16-F1 (orange circles), NHSL1 CRISPR clone 2 (magenta squares) and NHSL1 CRISPR clone 21 (green triangles) cells expressing the Arp2/3 biosensor as shown in (D). **(F)** Quantification of average cellular FRET efficiency which represents Arp2/3 activity from 4 independent biological repeats from control: 19 cells (orange circles); NHSL1 CRISPR 2: 21 cells (magenta squares); NHSL1 CRISPR 21: 11 cells (green triangles). Data points are the weighted average means for each cell, calculated from the FRET efficiency histograms. Data points represent the average weighted mean ± SEM (error bars), Kruskal-Wallis test: p=0.0091; Kruskal Wallis statistic: 9.407 and Dunn’s multiple comparisons test: control versus CRISPR 2 *, p=0.0136; control versus CRISPR 21 *, p=0.0260. Source data are provided as a Source Data file.

To test whether the observed phenotypes in the NHSL1 CRISPR knockout clones were not due to off-target effects, we re-expressed Myc-tagged wild type NHSL1 (Myc-NHSL1 WT) or the NHSL1 Scar/WAVE complex binding mutant (Myc-NHSL1 SW Mut) in the NHSL1 CRISPR 2 cell line and also expressed our Arp2/3 biosensor and quantified FRET by FLIM. Again, we observed a significant increase in Arp2/3 activity in the NHSL1 CRISPR 2 cell line compared to control B16-F1 cells, which was rescued with the wild type NHSL1. This confirmed that indeed the observed increase in Arp2/3 activity in the NHSL1 CRISPR 2 cells was due to loss of NHSL1 and not due to off target effects (Fig. S14G). We also found a significant rescue with the NHSL1 Scar/WAVE complex binding mutant (Fig. S14G) suggesting that in whole cells NHSL1 affects Arp2/3 activity through other mechanisms in addition to its interaction with the Scar/WAVE complex.

As an additional control we performed immunofluorescence analysis on wild type B16-F1 cells or the NHSL1 CRISPR 2 and 21 cells and quantified whole cell Arp2/3 intensity. We found that total Arp2/3 levels were not increased but rather reduced in NHSL1 CRISPR cells (Fig. S15A,C) and this excludes the possibility that the observed increase in Arp2/3 activity is due to increased Arp2/3 levels when NHSL1 is knocked out. Similarly, western blot analysis revealed that both total cellular Arp2/3 complex (subunit ARPC2) and Scar/WAVE2 levels were reduced in the NHSL1 CRISPR 2 line (Fig. S15D-F), suggesting that persistent overactivation of the Scar/WAVE and Arp2/3 complexes in the absence of NHSL1 may lead to their proteasomal degradation^31,32^. Consistent with a reduction in total Arp2/3 levels, we observed that the total cellular F-actin content was significantly reduced in NHSL1 CRISPR 2 cells compared to wild type cells. This phenotype was rescued by re-expressing wild type Myc-NHSL1 but not Myc-NHSL1 SW Mut in the NHSL1 CRISPR 2 cells (Fig. 8C; S16A) suggesting that NHSL1 controls cellular F-actin levels by binding to the Scar/WAVE complex. Similarly, we observed a significant reduction in total cellular F-actin content upon knockdown of NHSL1 and, conversely, an increased total cellular F-actin content upon overexpression of EGFP-NHSL1 in HEK293 cells (Fig. S16 B,C).

**Figure 8.**
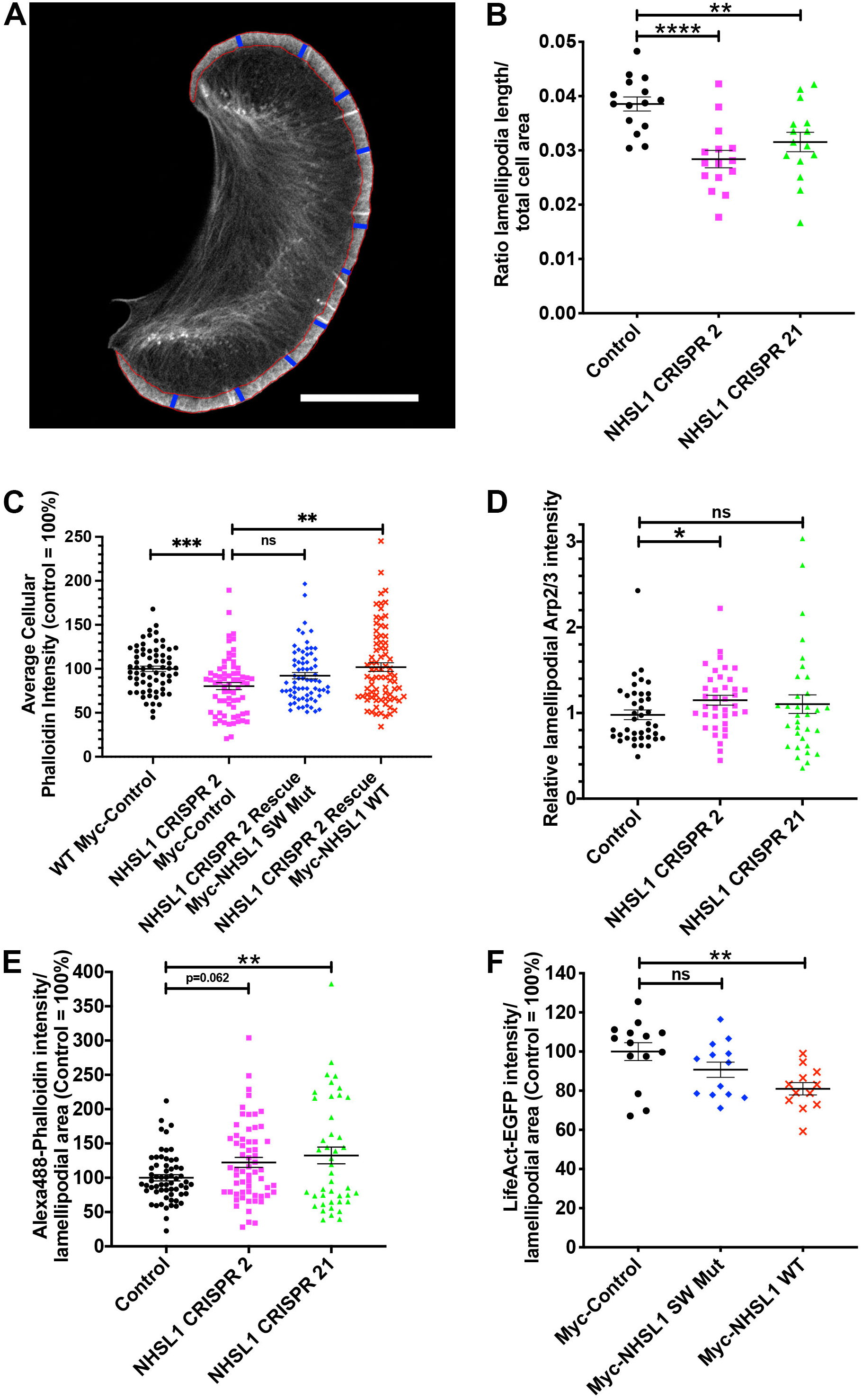
NHSL1 negatively regulates Arp2/3 and F-actin content in the lamellipodium. **(A)** Example B16-F1 cell shown indicating definition of length and width used in (B; S17A-D): First frame of a movie of LifeAct-EGFP expressing B16-F1 cells to show how length, width and area was quantified: red line indicates area of the lamellipodium. The length was quantified using the Fiji plugin “measure_ROI” (http://www.optinav.info/Measure-Roi.htm:Measure_Roi_Curve.java) which measures the length of curved objects (see materials and methods for details). For the width, the line tool in Fiji was used to draw lines (blue lines in (A)) at 90 degree angle to the leading edge to measure width of lamellipodium at 10 roughly equal distance points along the leading edge and the mean of these measurements was used. Scale bar: 20 μm. **(B)** Quantification of the ratio lamellipodia length/total cell area, One-way ANOVA, Dunnett’s multiple comparisons test; ****, p< 0.0001; **, p=0.0059; F(2,42)=10.84 **(C)** Quantification of total cellular F-actin (Alexa568 phalloidin) intensity/area ratio in wild type B16-F1 cells expressing Myc alone as control (black circles) or CRISPR 2 cells expressing either the NHSL1 mutant in the Scar/WAVE complex binding sites (NHSL1 SW Mut, blue diamonds) or NHSL1 (NHSL1 WT, red crosses) or Myc alone as control (pink squares) plated on laminin. Results are mean ± SEM (error bars) from four independent biological experiments. One-way ANOVA: p=0.0009; Kruskal Wallis statistics 16.44; ***, p=0.0003; **, p=0.0083; ns, p=0.1518; Outliers were removed using the ROUT method (Q=1%) (n analysed/ n outliers = n_o_ removed); WT control: n=67 cells/ n_o_=0; NHSL1 CRISPR2: n=71 cells/ n_o_=3; NHSL1 CRISPR2+Myc-NHSL1 SW Mut Rescue: n=76 cells/ n_o_=5; NHSL1 CRISPR2+Rescue Myc-NHSL1 WT: n=82 cells/ n_o_=4. **(D)** Quantification of relative lamellipodial Arp2/3 intensity: Arp2/3 intensity in the lamellipodium was normalized cell-by-cell against Arp2/3 intensity of the whole cell. Kruskal-Wallis test (p=0.0514) and Dunn’s multiple comparisons test: * p=0.0353; ns p>0.9999; in wild-type (n=40 cells, control), NHSL1 CRISPR 2 (n=38 cells) and NHSL1 CRISPR 21 (n=33 cells) B16-F1 cells plated on laminin and stained with anti-ARPC2 (subunit of Arp2/3 complex) antibodies; Results are mean ± SEM (error bars), three independent biological repeats. **(E)** Quantification of lamellipodial F-actin (Alexa488 phalloidin) intensity/area ratio in B16-F1 wild-type control cells or NHSL1 CRISPR 2 and NHSL1 CRISPR 21 B16-F1 cells plated on laminin. Results are mean ± SEM (error bars), five independent biological repeats, n= control (61), CRISPR 2 (61), CRISPR 21 (43) cells; One-way ANOVA: p=0.0127; F(2,162)=4.486; and Dunnett’s multiple comparisons test: **, p=0.0099. **(F)** Quantification of lamellipodial F-actin (LifeAct-EGFP) intensity/area ratio in B16-F1 cells transfected with a tri-cistronic plasmid to express Myc-tagged NHSL1 (wild type or Scar/WAVE binding mutant), puromycin resistance and LifeAct-EGFP or empty plasmid (Myc-IRES-Puro-T2A-LifeAct-EGFP) after selection with puromycin plated on laminin. Results are mean ± SEM (error bars), four independent experiments, n= control (14), WT (12), NHSL1-SW mut (13) cells. One-way ANOVA: p=0.0074; F(2,36)=5.636; and Dunnett’s multiple comparisons test: **, p=0.0036; ns p=0.1810. Source data are provided as a Source Data file.

### Loss of NHSL1 increases lamellipodial Arp2/3 and F-actin content

To gain a deeper insight on how NHSL1 affects cell motility we quantified lamellipodium parameters. We transfected our stable NHSL1 CRISPR B16-F1 cell lines or control wild type B16-F1 cells with LifeAct-EGFP. We found that NHSL1 CRISPR 2 and 21 cells had a larger cell area (Fig. S17A) and thus we normalised the lamellipodia length to total cell area. This analysis revealed that the NHSL1 CRISPR 2 and 21 cells have a significantly reduced ratio of lamellipodia length to cell area (Fig. 8A,B; S17A,B). However, both lamellipodia width and number of microspikes per length of lamellipodium were unchanged (Fig. 8A; S17C,D).

To quantify Arp2/3 activity in lamellipodia we sought to estimate Arp2/3 biosensor FRET efficiency at the leading edge by manually outlining lamellipodia. This approach is limited by the resolution of the FRET-FLIM microscope. Nevertheless, we observed a change in FRET efficiency from 0.01 to 3.97% for Control vs. CRISPR 2 (p=0.6914) and 0.01 to 8.87% for Control vs. CRISPR 21 (p=0.2502) for Arp2/3 activity in the lamellipodium. Neither changes are significant (Fig. S14H).

As an alternative method, endogenous Arp2/3 staining intensity in the lamellipodium can be used as an approximation of Arp2/3 activity since Arp2/3, after activation at the leading edge, is incorporated into the lamellipodial F-actin meshwork. We found that relative Arp2/3 complex (ARPC2) intensity was increased in the lamellipodium in the NHSL1 CRISPR 2 cells (Fig. 8D; S15A). This suggests that NHSL1 functions to reduce Arp2/3 activity in the lamellipodium. When we quantified Alexa488-phalloidin labelled F-actin intensity in the lamellipodium, we observed an increase in intensity per area in the NHSL1 CRISPR cells (Fig. 8E; S15B). Conversely, we found a significant reduction of lamellipodia LifeAct-EGFP intensity per area upon wild type NHSL1 overexpression but not with the NHSL1 Scar/WAVE binding mutant (Fig. 8F; S17E) suggesting that NHSL1 may inhibit F-actin nucleation in the lamellipodium by binding to Scar/WAVE.

### NHSL1 CRISPR knockout decreases F-actin retrograde flow speed in lamellipodia

To explore effects of NHSL1 on F-actin retrograde flow in the lamellipodium we again used our NHSL1 CRISPR 2 cell line and rescued it with Myc-NHSL1 WT or the Myc-NHSL1 SW Mut and transfected them with LifeAct-EGFP (Fig. 9A; video S8). The protrusion speed of lamellipodia was not significantly changed by NHSL1 knockout (Fig. S18A). However, particle image velocimetry analysis revealed a significant reduction of F-actin retrograde flow in the NHSL1 CRISPR 2 lamellipodia compared to wild type control lamellipodia (Fig. 9A,B). Re-expression of EGFP-NHSL1 WT in the NHSL1 CRISPR 2 cells changed the mean values of the retrograde flow speed in lamellipodia from 1.34 to 1.43 μm/min and re-expression of EGFP-NHSL1 SW Mut changed the mean values of the flow speed from 1.34 to 1.51 μm/min but these changes were not significant (Fig. 9A,B).

**Figure 9.**
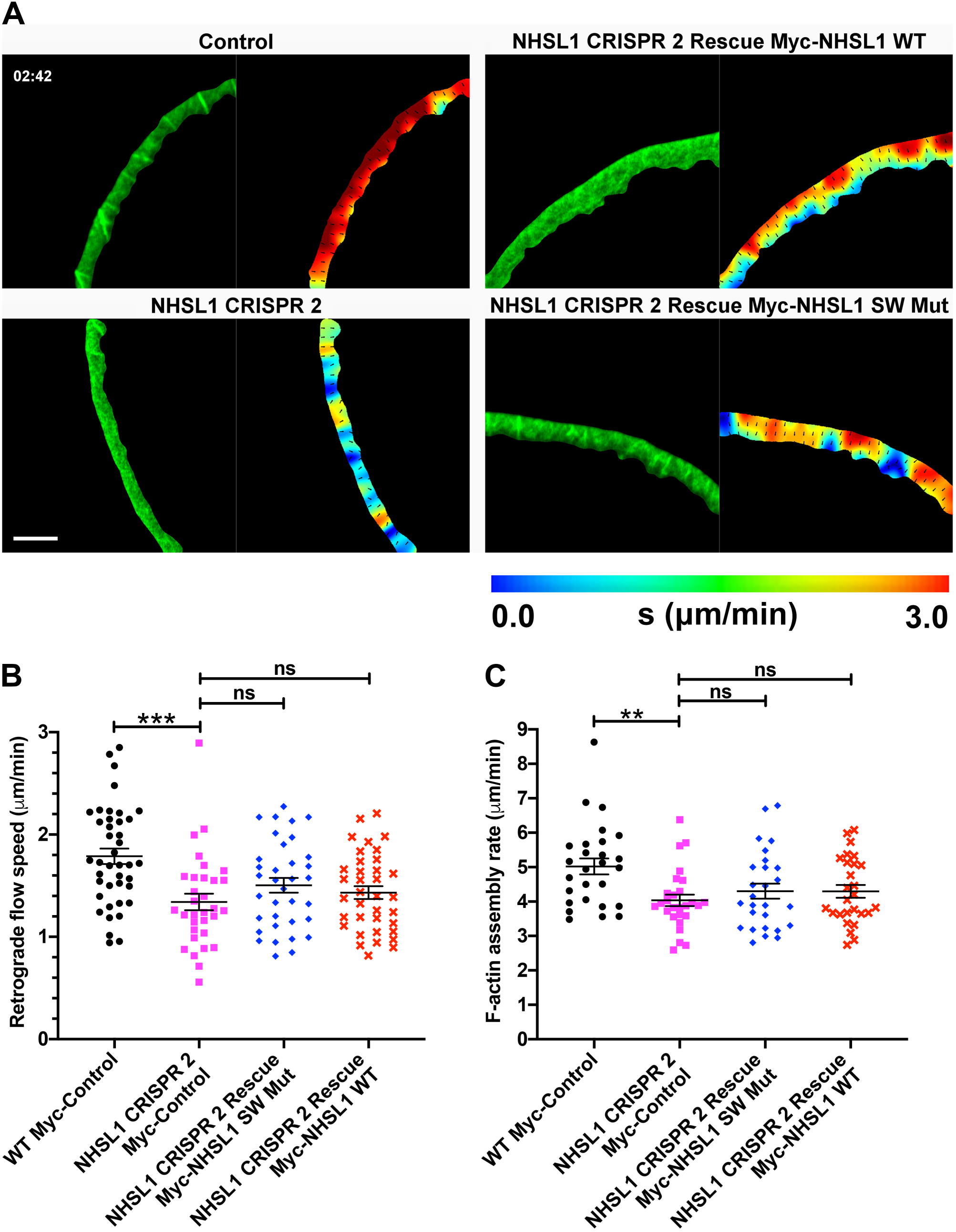
NHSL1 increases actin retrograde flow speed and F-actin assembly rate. (**A**) A representative snapshot in time (02:42 minutes:seconds) of a segmented lamellipodium from a wild type cell (WT), NHSL1 knockout cell (NHSL1 CRISPR 2), and NHSL1 knockout cells rescued with a wild type NHSL1 construct (NHSL1 CRISPR 2 Rescue Myc-NHSL1 WT) and with a mutant NHSL1 construct (NHSL1 CRISPR 2 Rescue Myc-NHSL1 SW Mut) using a bicistronic expression plasmid also conferring resistance to blasticidin to ensure that all cells analysed expressed NHSL1 and also expressing lifeAct-EGFP. Left panels show a frame from segmented Airyscan movies; right panels show the corresponding frame from the PIV interpolated movie. Each colourmap has the same range (0-3 μm/min) and distance between vector arrows (1 μm); warmer colours represent faster flow, vector arrows are all of unit length. Scale bar = 10 μm. (**B**) Quantification of the average speed of actin retrograde flow for each condition over all frames for each cell. Flow is significantly reduced in NHSL1 knockout cells but cannot be fully rescued by either the WT or mutant NHSL1 constructs. Results are mean ± SEM; n = 41, 32, 35, and 36 for the respective conditions (left to right on x-axis), from 6 independent biological repeats. One-way ANOVA: p=0.0001; F(3,140)=7.312; Tukey’s multiple comparisons test: *** p=0.0002, ns = not significant: CRISPR2 versus rescue Myc-NHSL1 SW Mut: ns p=0.4285; CRISPR2 versus rescue Myc-NHSL1 WT: ns p=0.8275. (**C**) Quantification of the F-actin assembly rate for each condition over all frames for each cell. The F-actin assembly rate is significantly reduced in NHSL1 knockout cells but cannot be rescued by either the WT or mutant NHSL1 constructs. Results are mean ± SEM; n = 41, 32, 35, and 36 for the respective conditions (left to right on x-axis), from 6 independent biological repeats. One-way ANOVA: p=0.0059; F(3,104)=4.400; Tukey’s multiple comparisons test: ** p=0.0044, ns = not significant: CRISPR2 versus rescue Myc-NHSL1 SW Mut: ns p=0.7874; CRISPR2 versus rescue Myc-NHSL1 WT: ns p=0.7967. Source data are provided as a Source Data file.

A more intuitive measure of effective actin polymerisation at the lamellipodium is the F-actin assembly rate which takes the forward protrusion of the plasma membrane and the retrograde actin flow into account and can be calculated in this case as the sum of the magnitude of the vectors of the flow speed and the protrusion speed^33^. Even though we measured increased Arp2/3 activity in the NHSL1 CRISPR 2 cells (Fig. 7D-F), we surprisingly observed a significant reduction of F-actin assembly rate in NHSL1 CRISPR 2 lamellipodia compared to wild type control lamellipodia (Fig. 9A,C). Re-expression of EGFP-NHSL1 WT or the EGFP-NHSL1 SW Mut changed the mean values of the F-actin assembly rate for both from 4.04 to 4.30 μm/min compared to the NHSL1 CRISPR knockout lamellipodia but this change was not significant (Fig. 9A,C). This seemingly counterintuitive observation that NHSL1 CRISPR knockout causes increased Arp2/3 activity yet decreases F-actin retrograde flow and assembly rate in the lamellipodium may be explained by the effects of higher filament densities on membrane tension and the tension gradient across the lamellipodium^33,34^. Our observations are consistent with results from modelling of actin filament density in the lamellipodium and calculating resulting F-actin retrograde flow, protrusion speed, and assembly rate^33,34^ (see discussion).

### NHSL1 reduces stability of lamellipodia protrusion

Since NHSL1 reduced Arp2/3 activity (Fig. 7) we further explored whether NHSL1 affects lamellipodia dynamics and/or lamellipodia stability. We quantified leading edge morphodynamics (video S9, video S10) and this revealed that protrusion speed did not change significantly in the NHSL1 CRISPR cells compared to control (Fig. 10E) in agreement with lamellipodia protrusion speed measurements during flow analysis (Fig. S18A). In addition, we did not observe a significant change in lamellipodia protrusion speed when we re-expressed wild type or the Scar/WAVE binding mutant of NHSL1 in the NHSL1 CRISPR 2 knockout cell line (Fig. 10E).

**Figure 10.**
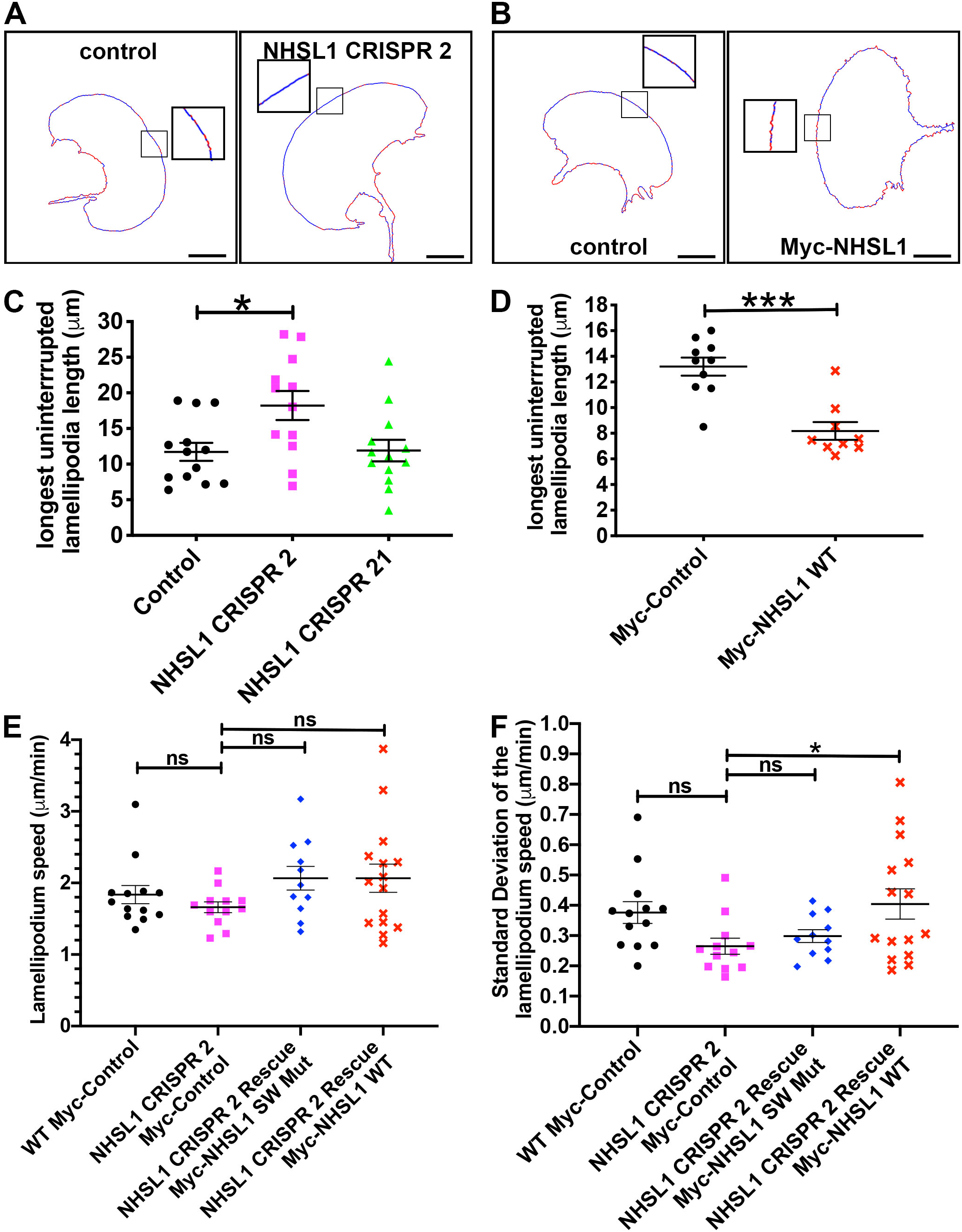
NHSL1 reduces stability of lamellipodia protrusion. **(A,B)** Still images from movies showing protrusion dynamics (blue = protrusion, red = retraction) of wild-type B16-F1, NHSL1 CRISPR clone 2 and B16-F1 cells overexpressing Myc-tagged NHSL1 by automatically segmenting the cell outline in each frame of the movies and calculating protrusion vectors at each pixel along the cell edge. Scale bar: 20 μm. **(C,D)** NHSL1 reduces length of longest uninterrupted lamellipodium. Each data point, square or triangle represents the mean length of longest uninterrupted lamellipodium automatically quantified from all frames of a movie of one LifeAct-EGFP expressing B16-F1 cell (5 sec intervals, 10 minutes duration) for control (black circles) and NHSL1 CRISPR cell lines 2 (magenta squares) and 21 (green triangles) (C) and control (black circles) and Myc-NHSL1 overexpression (red crosses) (D). Results are mean ± SEM (error bars), (C) control: n=13; NHSL1 CRISPR 2: n=12; NHSL1 CRISPR 21: n=13 cells, four independent biological repeats. One-way ANOVA: p=0.0111; F(2,35)=5.133; and Dunnett’s multiple comparisons test *, p=0.0144, control versus CRISPR21: not significant: ns p=0.9949. (D) control n = 10; Myc-NHSL1 n=9 cells, four independent biological repeats. ***, p=0.006, Mann-Whitney test. **(E,F)** Quantification of lamellipodia protrusion speed (E) and lamellipodia stability (F) of randomly migrating wild type B16-F1 cells or CRISPR 2 cells expressing either NHSL1 (NHSL1 WT) or the NHSL1 mutant in the Scar/WAVE complex binding sites (NHSL1 SW Mut) or Myc alone as control plated on laminin by automatically segmenting the cell outline in each frame of the movies and calculating protrusion vectors at each pixel along the cell edge. Lamellipodia protrusion speed was quantified in a cone of 60 degree centered about the direction of migration as quantified by tracking the centroid of the cell. Results are mean ± SEM (error bars), WT Control: n=13, NHSL1 CRISPR 2: n=12, Rescue Myc-NHSL1 SW Mut: n=11, Rescue Myc-NHSL1 SW WT: n=15 cells, four independent biological repeats. One-way ANOVA: p=0.2097, F(3,47)=1.568; and Dunnett’s multiple comparisons test: NHSL1 CRISPR2 vs Rescue SW mut: ns: p=0.2043; NHSL1 CRISPR2 vs Rescue SW WT: ns: p = 0.1573; NHSL1 CRISPR2 vs WT control: ns p=0.7639. **(F)** The standard deviation of the cone speed shows the fluctuation of speeds along the edge and thus can serve as a measure for the stability of lamellipodial protrusions. Results are mean ± SEM (error bars), WT Control: n=13, NHSL1 CRISPR 2: n=12, Rescue Myc-NHSL1 SW Mut: n=11, Rescue Myc-NHSL1 SW WT: n=15 cells, four independent biological repeats. One-way ANOVA: p=0.0412, F(3,47)=2.971: and Dunnett’s multiple comparisons test: NHSL1 CRISPR2 vs Rescue SW mut: ns: p=0.8865; NHSL1 CRISPR2 vs Rescue SW WT: *, p = 0.0299; NHSL1 CRISPR2 vs WT control: ns p=0.1173. Source data are provided as a Source Data file.

Lamellipodia stability can be defined both temporally, as the duration of lamellipodium protrusion until the next retraction, as well as spatially: lamellipodia can be characterised by protruding segments interrupted by small retractions. Stable lamellipodia with longer sections of protrusions may more likely increase cell migration speed and persistence than lamellipodia with many retractions. We thus analysed lamellipodia stability and observed that the NHSL1 CRISPR cells displayed a significantly increased uninterrupted lamellipodium (Fig. 10A,C, video S9) whereas this parameter was significantly reduced in cells overexpressing EGFP-NHSL1 (Fig. 10B,D, video S10) compared to controls. When we compared the standard deviation (S.D.) of the lamellipodia protrusion speed from all frames of a movie as a readout for the temporal stability of protrusions over time, we observed a change from S.D. = 0.376 μm/min for WT Myc Control to S.D. = 0.265 μm/min for NHSL1 CRISPR 2 knockout cells. This standard deviation of the lamellipodia protrusion speed of the NHSL1 CRISPR 2 knockout cells was significantly increased in the NHSL1 CRISPR 2 cells re-expressing wild type but not the NHSL1 Scar/WAVE complex binding mutant (Fig. 10F). Taken together, this indicates that NHSL1 reduces the stability of lamellipodia protrusions potentially via its interaction with the Scar/WAVE complex.

## Discussion

The Scar/WAVE complex, which is activated by several signals including Rac, is essential for lamellipodia formation and promotes mesenchymal cell migration^2^. We postulate that the Scar/WAVE complex needs also to be negatively regulated at the leading edge to tightly control lamellipodial protrusion dynamics and cell steering. Yet, so far, no direct inhibitor of the Scar/WAVE complex has been described. Here we reveal that NHSL1 is a novel, direct, negative regulator of the Scar/WAVE complex, which consequently reduces Arp2/3 activity. This reduction in Scar/WAVE-Arp2/3 activity may cause a decrease in lamellipodia stability, and thus also cell migration efficiency.

NHSL1 knockout cells also displayed reduced retrograde F-actin flow speed which coincided with a reduced F-actin assembly rate (Fig. 9). Actin polymerisation at the very edge of lamellipodia creates the force for plasma membrane protrusion. How the assembly rate is divided between protrusion and retrograde flow controls the speed of protrusions. Dolati et al. modelled the relationship between actin filament density at the leading edge of cells and the resulting retrograde flow, lamellipodia protrusion and F-actin assembly rates^33^. The modelling scenario, which agrees best with experimental results, revealed that protrusion rate and assembly rate increase with actin filament density at the leading edge up to a certain threshold. This threshold corresponds to the density measured for wild type B16-F1 lamellipodia. When actin filament density is further increased the protrusion rate then plateaus and the actin assembly rate again decreases^33^. This is most likely due to membrane tension which counteracts the protrusive force^33,34^. In NHSL1 knockout cells Arp2/3 activity is increased (Fig. 7) and in the lamellipodium Arp2/3 and actin filament densities (Fig. 8) are higher compared to wild type B16-F1 cells. Thus, our observed experimental values of reduced retrograde flow coinciding with reduced F-actin assembly rates are consistent with the predicted rates from the model^33,35^. We did not observe an increase in protrusion speed at higher Arp2/3 activity. This may seem counterintuitive but agrees with the predictions of the model. Consequently, the faster cell migration speed observed in the NHSL1 knockout cells cannot be explained by a faster lamellipodial protrusion speed. The increase in cell migration speed may therefore be due to an increase in lamellipodial stability. Taken together, the observed reduction in retrograde flow and assembly rates fit very well with the increased Arp2/3 activity in the NHSL1 CRISPR cells. The resulting increased actin filament branching density in the NHSL1 CRISPR cells may exert more force to counteract the membrane tension and thereby increase lamellipodial stability and consequently increase cell migration efficiency.

We found that NHSL1 knockout cells display a reduction in relative overall lamellipodia length (Fig. 8) while the longest uninterrupted lamellipodia length increases (Fig. 10). How the overall length of lamellipodia is controlled and why some cell types such as fibroblasts have smaller lamellipodia than others, for example B16-F1 cells or fish keratocytes is unknown. The Scar/WAVE complex appears to form higher order linear oligomers^36^ but how a lateral extension of such oligomers is controlled is not known. Our finding that NHSL1 interacts with the Scar/WAVE complex and affects lamellipodia length and continuity opens the possibility that NHSL1 may be involved in this control of lateral lamellipodia propagation.

Several isoforms of NHSL1 have been predicted from the human genome sequence, which results from alternative splicing of several different exon 1 sequences (Ensembl,^13,14,16^). Only one out of several alternative exon 1 sequences includes a Scar homology domain (SHD) also found in Scar/WAVE family proteins which in Scar/WAVE proteins mediates the interaction with Abi, HSPC300, Nap1, and Sra1^2^. The NHSL1 isoform that we cloned harbours an exon 1 which does not include the SHD. A similar exon-intron structure has been described for the related Nance-Horan Syndrome (NHS) and NHSL2 proteins^13,14,37^ but overall conservation between family members is low at the amino acid level (NHSL1 versus NHS: 30.3% and NHSL1 versus NHSL2: 25.1% see Fig. S18B). The function of NHS in cell migration is not known but NHS has been reported to localise to lamellipodia in MTLn3 breast cancer cells^14^ which is consistent with our observation that NHSL1 localises to the edge of protruding lamellipodia (Fig. 1).

The SHD domain of NHS has been shown to co-immunoprecipitate with the Scar/WAVE complex components Abi, HSPC300, Nap1, and Sra1^14^. Here we show that an NHSL1 isoform without the SHD domain interacts with the entire Scar/WAVE complex via the SH3 domain of Abi. Interestingly, the two Abi binding sites which mediate binding to NHSL1 (site 2+3; Fig. 5G) are highly conserved in the NHS family (PFAM PF15273 HHM logo, (Fig. S18C,D)) even though overall conservation is low (Fig. S18B)^13,37^ highlighting the importance of Scar/WAVE complex interaction at these particular sites. Furthermore, this suggests that all isoforms of NHS may also directly interact with the Scar/WAVE complex via the Abi SH3 domain mediated by the two novel Abi binding sites identified here. We thus postulate that at least some functions such as regulation of cell migration may be conserved between the family members. Therefore, deregulated cell migration may play a role in the pathogenesis of Nance-Horan syndrome.

Overexpression of an isoform of NHS harbouring the SHD domain has been reported to inhibit the ability of MTLn3 breast cancer cells to respond to EGF by forming lamellipodia, whilst overexpression of an isoform of NHS without the SHD domain did not affect lamellipodia extension^14^. In agreement, we show that overexpression of the NHSL1 isoform without the SHD domain does not inhibit lamellipodia extension (video S1). Furthermore, we found that knockout of all isoforms of NHSL1 increases lamellipodia stability, cell migration speed and persistence (Fig. 2, 6, 10). However, only lamellipodia stability and cell migration speed but not persistence was rescued in the NHSL1 CRISPR knockout cells when we re-expressed wild type NHSL1. Since overexpression of NHSL1 also increased cell migration persistence, optimal expression levels of NHSL1 may be required for low cell migration persistence.

The zebrafish orthologue of NHSL1 had been identified in a genetic screen for defects in a specific developmental migration of facial branchiomotor neurons suggesting a critical role of NHSL1 in neuronal migration^38^. Here, we found that NHSL1 negatively regulates random 2D cell migration by reducing lamellipodia stability (Fig. 2, 6, 10). We hypothesise that this reflects a role in guidance of NHSL1 during developmental migration as cells have to be able to change their path. In addition, the neurons need to integrate additional inputs in this mode of migration since zebrafish NHSL1 functions together with the planar cell polarity components Scrib and Vangl cell autonomously in facial branchiomotor neurons, which involves the interactions between the neurons and their planar-polarised environment of the surrounding tissue^38^.

Active Rac binds to NHSL1 via two sites and thus Rac may recruit NHSL1 to the leading edge of cells (Fig. 3, S6, S7). Whether this interaction is direct or not needs to be investigated in future studies. We show here that NHSL1 is a direct interactor of the Scar/WAVE complex and co-localises with it at the very edge of lamellipodia. In addition, an EGFP-NHSL1 Scar/WAVE binding mutant localises at lamellipodia like wild type NHSL1 (Fig. S11F). Finally, NHSL1 knockout cells display increased Arp2/3 activity (Fig. 7). Taken together this suggests that NHSL1 may act in a Rac-NHSL1-Scar/WAVE-Arp2/3 pathway negatively regulating Scar/WAVE and thus also Arp2/3 activity.

Interestingly, the two Rac binding sites in NHSL1 do not resemble any known Rac binding sites such as the CRIB or DUF1394 domains. The latter domain is found in the Scar/WAVE complex component Sra1/PIR121 but also in CYRI. CYRI binds to active Rac via a DUF1394 domain thereby competing with Sra1/PIR121 in the Scar/WAVE complex for Rac binding and indirectly counteracting Scar/WAVE recruitment to the leading edge^39^.

Surprisingly we found that NHSL1 reduces cell migration via the Scar/WAVE complex. This is interesting since the Scar/WAVE complex also directly binds to Rac and this increases Arp2/3 activation via the Scar/WAVE complex. A similar paradoxical regulation has been observed at the level of the Arp2/3 complex: Arpin negatively regulates Arp2/3 activity whereas the Scar/WAVE complex positively regulates Arp2/3 activity, and both of these activities occur downstream of active Rac at the same time in a protruding lamellipodium. Through this paradoxical circuitry, called an incoherent feedforward loop^40^, Arpin controls cell migration speed and persistence^41^.

Similar to NHSL1, Lpd functions downstream of active Rac but in contrast, Lpd promotes cell migration speed and persistence^6,7^. Both Lpd and NHSL1 appear to co-localise at the very edge of lamellipodia during protrusion (Fig. S1E, video S3). We therefore propose that NHSL1 together with Lpd forms a novel incoherent feedforward loop at the level of the Scar/WAVE complex downstream of active Rac at the leading edge of cells in which Lpd promotes whereas NHSL1 inhibits cell migration via Scar/WAVE to enable cell steering.

## Materials and methods

### Molecular biology, plasmids, and reagents

Full length human NHSL1d cDNA including exon 1d and excluding exon 5 was generated by cloning overlapping EST clones Flj35425 and DKFZp686P1949 into pENTR3C (Invitrogen). Fragments of NHSL1 (Fig. 1E, 3A, S7A) were cloned into pENTR3C (Invitrogen). NHSL1 cDNA in pENTR was mutated using Quikchange (Agilent) to create double (sites 1+2 or sites 2+3) and triple (sites 1+2+3) SH3 binding sites: Site 1: aa518-524(R>A)AS(P>A)LK(P>A); Site 2: aa625-633KKPPLP(P>A)S(R>A); Site 3: aa836-844 KPK(P>A)KV(P>A)ER.

ARPC1B (HsCD00370547), ARPC3 (HsCD00288065), Sra1 (CYFIP1; HsCD00042136), PIR121 (CYFIP2; HsCD00045545), Nap1 (NCKAP1; HsCD00045562), Abi2 (HsCD00042752), and HSPC300 (C3orf10; HsCD00045008) cloned into pENTR233 or pDONR221 (Harvard Institute of Proteomics). pDONR221-WAVE-2 (German Resource Centre for Genome Research). hsAbi1d (BC024254; Geneservice) full-length and Abi1d-Δ-SH3 (aa 1–417) were cloned into pENTR11. Human Lpd (AY494951) was amplified and cloned into pENTR3C (Invitrogen)^6^. ARPC1B-mVenus-P2A-ARPC3-mTurq2 was generated by Gibson assembly (NEB Hifi, New England Biolabs, Inc.) in pENTR3C (Invitrogen). pENTR3C-ARPC1B-mVenus-P2A-ARPC3-mTurq2 was transferred to pRK5-Myc-DEST for CMV driven expression in mammalian cells.

cDNAs in pENTR or pDONR were transferred into tagged mammalian expression vectors using Gateway^®^ recombination (Invitrogen): pCAG-DEST-EGFP, pCAG-DEST-EGFP-T2A-Puro, pCAG-EGFP-DEST-IRES-Puro, pRK5-Myc-DEST, pCAG-Myc-DEST-IRES-Puro, pCAG-Myc-DEST-IRES-BLAST, pCAG-Myc-DEST-IRES-Puro-T2A-LifeAct-EGFP, pCDNA3.1-mScarlet-I-DEST, pV16-gateway (MBP) which were generated by traditional restriction-based cloning or Gibson assembly (NEB Hifi, New England Biolabs, Inc.) cloning.

The SH3 domains of Abi1d, c-Abl, and eight fragments covering the entire length of NHSL1 were cloned by PCR and restriction cloning into pGEX-6P1 and purified from *E.coli* on Glutathione-sepharose (Amersham). Dominant active Myc-tagged Rac1 Q61L in pRK5-Myc was kindly provided by Laura Machesky (Beatson Institute, Glasgow, UK).

The NHSL1 shRNA constructs (shRNA A: TGCCCATTCTGTGATGATTA shRNA B: TGGATAAATCCCTATCAAGA), Control-shRNA: GCCGATAACCGAGAATACC were cloned into pLL3.7Puro which is derived from pLL3.7 with the addition of a puromycin selection. All constructs were verified by sequencing.

### Northern blot and qPCR

The murine multiple tissue northern blot (Ambion) was probed with a 650 bp NHSL1 fragment covering GST-NHSL1-6 (Fig. 1A,E) according to manufacturer’s instructions.

For Fig. 2B RNA was extracted from cells using the RNeasy Mini Qiagen kit. cDNA was prepared using Superscript IV (Invitrogen) and expression levels of NHSL1 quantified using standard curves relative to GusB using gene specific primers (PrimeTime qPCR primer assays, IDT):

GusB: Mm.PT.39a.22214848 (primer 1: ACCACACCCAGCCAATAAAG; primer 2: AGCAATGGTACCGGCAG)

NHSL1 (detecting all isoforms): Mm.PT.58.41480072 (primer 1: TCTTCATCTTGATAATCGTCGCA; primer 2: CAAGCTCAACCTCAAATCAGTG) in combination with Bioline Sensimix on a Roche lightcycler 480II qPCR machine.

For Fig. S5B mRNA levels were measured using the Cells to Ct Kit (Applied Biosystems) and an NHSL1-specific Taqman assay (Hs00325291_m1, Applied Biosystems). The amount of remaining mRNA was calculated with the deltaCt method.

### Generation of CRISPR cell lines

Stable CRISPR NHSL1 knockout B16-F1 cell lines were made by knocking in a stop codon into exon 2, which is common to all isoforms. We utilised a CRISPR-Cas9 nickase approach with two sgRNA’s, which minimises potential off-target effects: To generate the donor plasmid for homology directed repair, genomic DNA from B16-F1 cells (QiaAmp genomic DNA kit, Qiagen) was amplified with primers adding a stop codon at the beginning of exon 2 of NHSL1 common to all isoforms and cloned into the targeting plasmid pHR110PA-1(5’MCS-EF1-RFP+Puro-3’MCS) (SBI). Two closely spaced NHSL1 specific sgRNA (sgRNA1: caccgTCGACTCTCCTCGTCCAAGT; sgRNA2: caccgCTGTCCACTACACGGCACCA) were designed using http://crispr.mit.edu and cloned into pX330S-2 (Addgene 58778) and pX330A_D10A_x2 (Addgene 58772) harboring the Cas9 nickase and combined into one plasmid using a golden gate reaction^33^. B16-F1 cells were transiently transfected with both targeting plasmid and nickase plasmid including 2 sgRNA’s and after selection with puromycin, clonal cell lines were generated by limited dilution and reduction in expression of NHSL1 tested in a western blot. Cells were transiently transfected with a plasmid driving Cre recombinase expression from a PGK promoter and FACS sorted for absence of RFP. Genomic DNA was extracted using Qiagen QIAamp DNA mini kit and knockout cell lines genotyped using primers amplifying insertion of the selection cassette into the correct locus: left arm specific (primer 1: aggtctacactctgccgttctggg; primer 2: CGGCCGCtgTCTAGATTTTTGAa) or to detect WT or KO alleles after Cre mediated excision of the selection cassette: Primer pair 1: (gccccagcctcgcaatg; ccgtccagcaagcccagaga); primer pair 2: aatcctcgtgccccagcctc; gcggacgtgcaagccagtta).

### Antibodies

Immobilised GST-NHSL1-6, 7, 8 (mAb) or GST-aa387-546NHSL1 (pAb) (Fig. 1F) were digested with Prescission protease (Amersham Pharmacia Biotech) and used to produce monoclonal antibodies as described^42^ and to raise polyclonal rabbit antiserum #4457 (Eurogentec). The NHSL1 monoclonal antibody was subcloned twice (clone C286F5E1; IgG1). Commercial primary antibodies: EGFP (Roche), Myc (9E10, Sigma), MBP (New England Biolabs), Abi1 (MBL, clone 1B9, D147-3), Abi2 (Epitomics, 3189-1), Scar/WAVE1 (BD 612276), Scar/WAVE2 rabbit mAb (D2C8, CST 3659), ARPC2 (07-227-I-100UG, Millipore). Secondary antibodies: HRP-goat anti-rabbit, -goat anti-mouse (Dako).

### Cell culture and transient transfections

HEK 293FT cells (Thermo Fisher Scientific R70007) and B16-F1 mouse melanoma cells (ATCC CRL-6323) were cultured in Dulbeccos modified Eagle’s medium containing penicillin/streptomycin, L-glutamine and 10% fetal bovine serum. MCF10A cells (ATCC CRL-10317) were cultured as previously described^43^. Transient transfections in HEK 293FT cells were carried out using Lipofectamine 2000 (Invitrogen) according to manufacturer’s instructions. Transient transfections with B16-F1 cells were carried out with X-tremeGene 9 (Roche) and replaced with normal growth media 4-6 hours after transfection. All cells were maintained at 37°C in 5% (MCF10A) or 10% (B16-F1, HEK 293FT) CO_2_.

### Immunoprecipitations, Pulldowns and Western Blotting

Cells were lysed in glutathione S-transferase (GST) buffer (50mM Tris-HCL, pH 7.4, 200 mM NaCl, 1% NP-40, 2 mM MgCl_2_, 10% glycerol, NaF +Na_3_VO_4_, complete mini tablets without EDTA, Roche). Lysates were incubated on ice for 15 minutes and centrifuged at 17,000x*g* at 4°C for 10 minutes. Protein concentration was then determined (Pierce BCA protein assay kit; Thermo Fisher Scientific). For pulldowns, the lysate was incubated with either glutathione beads (Amersham), GFP-trap or Myc-trap beads (Chromotek). For immunoprecipitations, Protein A bead precleared lysates were incubated with primary antibody or non-immune control IgG followed by 1% BSA blocked protein A beads (Pierce) or protein A/G beads (Alpha Diagnostics). For GFP-trap or Myc-trap pulldowns, beads were blocked with 1% BSA before incubating with lysates for 1-2 hours or 30 mins, respectively. Following bead incubation, all beads were washed with lysis buffer, separated on SDS-PAGE gels and transferred onto Immobilon-P membranes (EMD Millipore). Western blotting was performed by transferring at 100 V, 350 mA, 50 W for 1.5 hours before blocking in 5% BSA, 5% or 10% milk overnight followed by 1 hour incubation with the indicated primary antibodies followed by HRP conjugated secondary antibodies (Dako) for 1 hour at room temperature. Blots were developed with the Immun-Star WesternC ECL kit (Bio-Rad Laboratories) using the Bio-Rad Imager and ImageLab software.

### Far-Western blot

GST and MBP fusion proteins were purified from BL21-CodonPlus (DE3)-RP *E. coli* (Stratagene) using glutathione (GE Healthcare) or amylose (New England Biolabs, Inc.) beads. Purified GST-NHSL1 fragments were separated on SDS-PAGE and transferred to PVDF membranes and overlayed as described previously^44^ with purified MBP-Abi full-length or MBP-Abi-delta-SH3, and MBP was detected with MBP antibodies (New England Biolabs).

### Immunofluorescence and live cell imaging

For immunofluorescence analysis cells were plated on nitric acid washed coverslips (Hecht-Assistant), coated with 25 ug/ml laminin (L2020, Sigma) and fixed 10 minutes with 4% paraformaldehyde-PHEM (60mM PIPES, 25mM HEPES, 10mM EGTA, 2mM MgCl_2_, 0.12M sucrose). For NHSL1 mAb and ARPC2 pAb (07-227-I-100UG, Millipore): cells were permeabilised 2 min with 0.1% Triton-X-100/TBS and quenched for 10 min with 1 mg/ml Sodium Borohydride; For NHSL1 pAb: cells were permeabilised with 0.05% saponin, 2 hours at RT. For all antibodies: cells were blocked with 10% normal goat serum and 10% BSA, TBS. Secondary antibodies: goat anti-rabbit, or anti-mouse Alexa488 or 568 (Invitrogen) and mounted in Prolong Diamond (Invitrogen). For Alexafluor 488 or 568-conjugated Phalloidin staining: cells were fixed 20 minutes with 4% paraformaldehyde-PHEM at room temperature. Cells were permeabilised for 2 minutes with 0.1% Triton-X-100/TBS and washed in a TBS solution. Cells were stained with Alexafluor 488 or 568-conjugated Phalloidin (Invitrogen) diluted 1:250 in 10% normal goat serum/TBS for 1 hour at room temperature, then washed three times in a TBS solution, and mounted in a Prolong Diamond (Invitrogen). Cells were imaged on an IX81 Olympus microscope (see below). Equal exposure times were used for all cells and all conditions for a given experiment. Intensity values were then normalised for each experiment to the mean of the control condition for that experiment.

Cell profiler analysis of F-actin content in HEK293 cells was analysed using Cellprofiler software (http://www.cellprofiler.org, Broad Institute). Briefly, cells were identified based on the nuclear DAPI and their phalloidin staining. F-Actin content was measured using the MeasureObjectIntensity module. Data was exported into a text file and analysed with Graphpad Prism. Values obtained from control samples (GFP or scrambled shRNA) where set to 100%.

For low magnification phase contrast and high magnification imaging, cells were plated on 12-well tissue culture dishes or glass bottom dishes (Ibidi; 81218-200) coated with 10 µg/ml fibronectin (F1141, Sigma) or coated with 25 µg/ml laminin (L2020, Sigma). For immunofluorescence and live imaging an IX 81 microscope (Olympus), with a Solent Scientific incubation chamber, filter wheels (Sutter), an ASI X-Y stage, Cascade II 512B camera (Photometrics), and 4× UPlanFL, 10× UPlanFL, 60× Plan-Apochromat NA 1.45, or 100× UPlan-Apochromat S NA 1.4 objective lenses) controlled by MetaMorph software or an LSM 880 Airyscan confocal microscope (Zeiss) with environmental chamber (37°C; 10% CO_2_) driven by Zen Black software with a 63× oil (NA 1.4) Plan-Apochromat objective (Zeiss) was used. Supplementary movies were prepared in Metamorph and Fiji.

### Scratch assays and quantification of random cell migration speed and persistence

Confluent control or shRNA MCF10A cells were scratched with a P200 pipette tip and treated with mitomycin C to inhibit proliferation. Movies were acquired for 12 hours with a frame every 5 minutes. The scratch area was measured at 0 and 12 hours with Fiji.

For random migration, B16-F1 cells were plated on to fibronectin or laminin coated 12-well dishes for 3 hours before imaging for 24 hours every 5 minutes. Cells were manually tracked by their nuclear position using the Manual Tracking plugin (FIJI), and the cell track coordinates imported into Mathematica for analysis using the Chemotaxis Analysis Notebook v1.6β (G. Dunn, King’s College London, UK^6^).

Speed and persistence measurements from tracking data are susceptible to positional error (due to manual tracking) as well as biological noise (from cell morphology changes or nuclear repositioning). To address the former, we estimated the positional error by tracking the same cell multiple times (each time blinded to the previous track), and then calculated the time interval (as an integer multiple of the frame interval) over which this error fell to at least below 10% of the average displacement measurement. We selected the smallest multiple for which this condition was true so as to avoid error from sampling too infrequently to measure the true path length. This time interval is termed the ‘usable time interval’ and denoted *δt* (see Fig. 2C, upper panel). In order to be consistent over data sets we a adopted a usable time interval that was common to all data within a dataset. This was *δt* = 15 (equivalent to a multiple of 3 frame intervals or taking a displacement measurement every 3 frames) for the CRISPR knockout and overexpression experiments on fibronectin and *δt* = 25 minutes (equivalent to a multiple of 5 frame intervals or taking a displacement measurement every 5 frames) for the CRISPR 2 knockout and rescue experiments on laminin. This ‘usable time interval’ approach also helps to reduce the impact of biological noise on the data.

Cell speed is the displacement (*d_n_*) over *δt*, where *n* denotes which interval of the track we are measuring (for example *n* = 1 would indicate the first 15 minutes, *n* = 2 the second 15 minutes etc.). The Mean Track Speed (*MTS*) is then the average cell speed over the whole track length (Fig. 2C, upper panel, boxed equation).

Cell persistence has traditionally been defined by the directionality ratio: the net (straight line) displacement divided by the total track length. This measurement comes with additional issues to noise, namely that it is dependent on interval size, track length, and even cell speed. The interval size is determined by the usable time interval, *δt* for the same reasons as previously described. To address the dependence on track length, we measure persistence over another time interval, denoted Δ*t*, which is an integer multiple of *δt* (see Fig. 2C, lower panel). This multiple is called the Time Ratio (*TR*, equal to Δ*t/δt*). This approach allows tracks of different total length to be compared equally, without skewing average results.

Persistence is defined by the Mean Track Persistence (*MTP*), equal to the average directionality ratio over the intervals Δ*t* (Δ*t* = 60 mins for *TR* = 4), multiplied by a normalisation factor, α, which sets the maximum persistence to 1 (a perfectly straight migration path) and the minimum persistence to that of a purely random walk. The directionality ratio is the net displacement, *D_N_*, where *N* is the interval number for each Δ*t* (i.e. *N* = 1 indicates the first 60 minutes, *N* = 2 indicates the second 60 minutes etc.) divided by the track length over Δ*t* (the sum of the displacements *d_n_*) (Fig. 2C, lower panel, boxed equation).

Finally, the issue of cell speed dependence can be addressed by selecting an appropriate combination of *δt* and *TR*. Plotting *MTS* as a function of *δt* reveals that speed falls off exponentially with increasing time interval, for all intervals which yield displacement values above the level of track noise. The mean persistence profile is a plot of the MTP obtained for all possible choices of *δt* for a given *TR*. In the mean persistence profile, the persistence normally starts off low at small speed intervals and rapidly increases to a peak before falling off again. To avoid noise, we chose a suitable *δt* and *TR* combination allowing the MTP to be measured close to this peak. The same *TR* must be used across all data sets so they are directly comparable; thus, another compromise must be that the *TR* chosen is appropriate for all populations. After some investigation, a *δt* of 15 and *TR* of 4 was chosen, for the CRISPR knockout and overexpression experiments on fibronectin and a *δt* of 25 and *TR* of 2 was chosen for the CRISPR 2 knockout and rescue experiments on laminin. These combinations ensured in both cases that tracks were long enough to contain multiple intervals.

The direction autocorrelation, directionality ratio over time and mean square displacement was calculated and plotted using excel macros provided in Gorelik and Gautreau, Nature Methods 2014^19^ according to the instruction provided.

### Quantification of lamellipodia dynamics

B16-F1 cells expressing LifeAct-EGFP plated on laminin were imaged for 10 minutes at 5 seconds per frame using an LSM 880 Airyscan confocal microscope or IX81 Olympus microsope and cells were automatically segmented and protrusion vectors at each pixel along the cell edge calculated using a MATLAB script kindly provided by Andrew Jamieson and Gaudenz Danuser (UT Southwestern, USA). Tracking the direction of migration of these cells allowed us to automatically quantify membrane protrusion speed only of the lamellipodium in the direction of migration. Custom written MATLAB scripts and Metamorph journals were used to calculate protrusion speed, the longest uninterrupted lamellipodium, and the length distribution of protrusions from all frames of the movie from the leading edge automatic profiling data. Fiji and Excel were used to quantify the LifeAct-EGFP intensity, total cell area, lamellipodia width and length, and microspike number per length of lamellipodium in the first frame of each movie. Lamellipodia length was quantified using the Fiji plugin “measure_ROI” (http://www.optinav.info/Measure-Roi.htm:Measure_Roi_Curve.java). This plugin measures the length of curved objects: For each particle, it first finds the points with the largest separation. These are designated as the end points. Next, it computes two curves that connect the end points along the left and right sides of the object. These curves are parameterized by arc length. The left and right side curves are averaged for each arc length to create a centerline curve. The arc length of the centerline curve is reported as the length of the object. Lamellipodia width was measured using the line tool in Fiji to draw lines at 90 degree angle relative to the leading edge at ten roughly equal distance points along the lamellipodium and these were averaged. Supplementary movies were prepared in Fiji.

### FRET-FLIM analysis of Arp2/3 activity

B16-F1 cells plated on nitric acid cleaned No 1.5 coverslips (Hecht Assistent), coated with 25 ug/ml laminin (Sigma) were fixed for 20 minutes with 4% paraformaldehyde-PHEM (60mM PIPES, 25mM HEPES, 10mM EGTA, 2mM MgCl_2_, 0.12M sucrose), permeabilised for 2 minutes with 0.1% TX-100/PBS, background fluorescence quenched with 1 mg/ml sodium borohydride in PBS and mounted in ProLong Diamond (Thermo Fisher).

Time-domain FLIM data were acquired via a time-correlated single photon counting (TCSPC) custom-built, automated, 2-photon microscope. Briefly, this consisted of a Modelocked femtosecond Ti:Sapphire laser (Coherent Vision II; Coherent (UK) Ltd, Scotland) for fluorescence excitation, a dual-axis scanner, a photomultiplier detector (HPM-100-06; Becker & Hickl GmbH, Germany) and TCSPC electronics (SPC830; Becker & Hickl GmbH, Germany). Images were acquired using a 1.3 NA 40× Plan Fluor Oil Immersion objective (Nikon Instruments Ltd, UK). Fluorescence lifetimes were determined for every pixel using a modified Levenberg-Marquardt fitting technique as described previously^45^. The FLIM images were batch analysed by running an in-house exponential fitting algorithm (TRI2 software) written in LabWindows/CVI (National Instruments, Austin, TX)^45^. The fitting parameters for each time-resolved intensity image were recorded in individual output files and used to generate a distribution of lifetime and an average fluorescence lifetime. FLIM/FRET analysis was performed to investigate Arp2/3 activity using the Arp2/3 biosensor (ARPC1B-mVenus-P2A-ARPC3-mTurq2). FRET efficiencies were calculated based on the equation *E* = 1 − *τ_DA_*/*τ_D_*, where *τ_D_* and *τ_DA_* are the measured fluorescence lifetimes of the donor in the absence and presence of the acceptor, respectively.

### Analysis of F-actin retrograde flow

Cells transfected with LifeAct-GFP were harvested and re-plated on to 35 mm glass-bottomed dishes (Ibidi), pre-coated with laminin (L2020, Sigma). The dishes were centrifuged for 2 minutes at 200 x g and incubated for ≥ 3 hours before imaging. The cells were then counterstained with 0.5 μM SiR-DNA (Spirochrome) for approximately 30 minutes prior to imaging. This stain was not used in actin flow experiments but was used for cell tracking in quantification of lamellipodial dynamics experiments. Videos were captured on a Zeiss LSM880 Airyscan confocal microscope at 5x zoom, 0.4-1.0% laser power, and frame rate of 3.22 seconds with at least 20 frames per movie.

#### Pre-processing

Zen Black software was used to Airyscan-process the raw data with automatic parameter fitting. Lamellipodia were segmented in Fiji by using a semi-automated method: frame-by-frame normalisation of contrast and Gaussian blur, and then correcting for photobleaching over the entire movie by histogram matching and using the result to construct a binary mask determined by semi-automatic thresholding with manual editing to remove certain features such as vesicles. This mask was then multiplied by the corresponding contrast-enhanced original Airyscan movie (not blurred or bleach-corrected) to obtain a segmented movie which formed the input for PIV. Movies were truncated when loss of focus, lamellipodial collapse, or significant translocation of the cell out of the field of view occurred.

#### PIV analysis

PIV analysis was carried out using a custom MATLAB script. The script utilises a two-dimensional cross-correlation algorithm adapted from classical PIV and is published here: ^46,47^. The method relies on searching for a region of interest (source area) in a larger region of a subsequent frame (search area) and finding the best match by cross-correlation. For our analysis we optimised the region size, overlap, and correlation threshold using a source box size of 0.3 μm, search box size of 0.5 μm, grid size (defining the overlap) of 0.2 μm, and correlation coefficient threshold of 0.5. Subsequently, spatial and temporal convolution are used to interpolate the displacement vectors over all the pixels in the image. Here, the spatial kernel size was 3 μm (σ = 0.5 μm) and the temporal kernel size was 15 s (σ = 6 s). All colourmaps presented were generated by setting a maximum flow velocity of 3 μm/min, and the number of vector arrows displayed is arbitrarily defined by the distance between them to represent the interpolation.

### Statistical analyses

Data were tested for normal distribution by D’Agostino & Pearson and Shapiro-Wilk normality tests. Statistical analysis was performed in Prism 8 (GraphPAD Software) using a Student’s *t*-test, One-way ANOVA or non-parametric Kruskal-Wallis test with appropriate post-hoc tests (see figure legends in each case). P values <0.05 were considered significant.

## Acknowledgments

We thank Laura Machesky (Beatson Institute, Glasgow, UK) for reagents. Andrew Jamieson and Gaudenz Danuser (UT Southwestern, USA) for providing the Windowing MATLAB script. William Barrell (King’s College London, UK) for qPCR training. F.M. was supported by a Medical Research Council (MRC) studentship. T.P. was supported by an Engineering and Physical Sciences Research Council (EPSRC) studentship. S.J. was supported by a Malaysian Public Service Department (PSD) studentship. This work was supported by grants from the European Research Council (ERC) under the European Union’s Horizon 2020 research and innovation programme (grant agreement No 681808) (B.M.S.), CRUK Comprehensive Cancer Centre block grant (S.A.-B), Medical Research Council (MRC) (S.A.-B.), Richard Dimbleby Cancer Trust (S.A.-B.), Wellcome Trust (107859/Z/15/Z) (B.M.S.) (082907/Z/07/Z) (M.K.), the Biotechnology and Biological Science Research Council, UK (BB/F011431/1; BB/J000590/1; BB/N000226/1; BB/R015953/1) (M.K.), and Cancer Research UK (S.A.-B.) (C22104/A7155) (M.K.).

## Data availability statement

The datasets generated or analysed during the current study are available from the corresponding author on reasonable request. All source data are provided as a Source Data file.

## Computer code availability statement

Custom written MATLAB code which has not been published before has been provided to the referees and is available as supplemental material.

## Availability of unique biological material generated in this study

All unique biological materials such as cell lines or plasmids generated in this study are available upon reasonable request from the corresponding author.

---

**Figure S1. NHSL1 does not interact with Lpd but co-localises with it at the leading edge of migrating cells**

**(A-C**) Full Western blots detecting NHSL1 protein in indicated cell lines using the (A,B) polyclonal (4457) or (C) monoclonal antibody (C286F5E1) from Figure 1B-D. Please note that both antibodies should recognise all isoforms, lower molecular weight bands may also represent degradation products). **(D**) Still images from live cell imaging showing EGFP-NHSL1 co-expressed with mScarlet-I only in B16-F1 cells to control for detection of leading edge due to space filling of the fluorescent protein. The subtracted image is generated from the subtraction of the signal of the mScarlet-I from the NHSL1-EGFP signal. Representative images from three independent biological repeats. Scale bar: 20 μm. Scale bar in inset: 5 μm. See also related video S2. **(E**) Still images from live cell imaging showing NHSL1-EGFP co-expressed with mScarlet-I-tagged Lpd in B16-F1 cells plated on laminin. Representative images from three independent biological repeats. Scale bar: 20 μm. Inset represents a magnified view of the white box. See also related video S3. **(F)** Western blot showing pulldown using GFP-trap beads from HEK cell lysates co-expressing either EGFP-Lpd or EGFP as control with Myc-NHSL1. Co-immunoprecipitation of Myc-NHSL1 with EGFP-Lpd was examined by western blot with antibody against GFP or Myc. Representative blots from three independent biological repeats. Source data are provided as a Source Data file.

**Figure S2. Genomic analysis of NHSL1 CRISPR knockout clones**

**(A,B)** Genomic characterization of the NHSL1 CRISPR clones revealed that the targeting cassette was inserted in the correct genomic NHSL1 location for CRISPR 2 but not inserted in CRISPR 21. Upper panel: schematic diagram of the NHSL1 targeting construct to knock in a stop codon into exon 2 and also a LoxP site flanked selection cassette. This diagram also indicates the locations of the genotyping primers for testing proper insertion at the genomic locus and the resulting PCR fragment size of 1300 bp. Please note that the 5’ primer sits outside the left arm and the 3’ primer is located in the selection cassette and thus this pair only amplifies a fragment if the targeting cassette was inserted in the correct genomic NHSL1 location. Lower panel: Agarose gel showing correct PCR fragment size of 1300 bp only for NHSL1 CRISPR 2 cell line. H_2_O instead of genomic DNA served as the negative control for specificity of the amplification. **(B)** After transient Cre recombinase expression in CRISPR 2, all wild type alleles are absent suggesting that the CRISPR 2 line represents a full NHSL1 knockout. Upper panel: schematic diagram of the NHSL1 wild type exon 2 or NHSL1 knockout exon 2 after Cre mediated excision of the selection cassette containing the knocked in stop codon. This diagram also indicates the locations of the genotyping primers for testing wild type or knockout alleles and the resulting PCR fragment size of Primer pair 1: WT 312 bp and KO 376 bp or Primer pair 2: WT 474 bp and KO 538 bp. Lower panel: Agarose gel showing PCR fragment sizes corresponding to WT alleles for WT and CRISPR 21 cells and only KO alleles for CRISPR 2 cells H_z_O instead of genomic DNA served as the negative control for specificity of the amplification.

**Figure S3. Characterisation of NHSL1 CRISPR knockout cell lines**

**(A)** Full western blot (same as shown in Fig. 2A) showing extent of reduction of NHSL1 expression in the clonal NHSL1 CRISPR B16-F1 cell lines 2 and 21 was probed with polyclonal NHSL1 antibodies and HSC70 antibodies as a loading control. Please note that lower molecular weight bands which may represent degradation products are also reduced in the CRISPR lines. Representative blot from three independent biological repeats. **(B-D)** Migration tracks of both NHSL1 CRISPR clones 1 and 21 in comparison to wild-type B16-F1 cells. **(E)** Relative expression of NHSL1 in wild type B16-F1 (control) or NHSL1 CRISPR 2 and 21 cells by qPCR using isoform independent gene specific primer sets compared to expression of housekeeping gene GusB. One-way ANOVA: p=0.2066; F(2,6)=46.6; Dunnett’s multiple comparisons test: *** p=0.0002; ns: not significant.

**Figure S4. NHSL1 negatively regulates cell migration persistence**

**(A, B)** Cell migration persistence was significantly increased for the NHSL1 CRISPR 2 or 21 cell line, respectively. **(A)** The directionality ratio is plotted to explore how it changes over the time of the movie. **(B)** Direction autocorrelation is shown, a measure of how the angle of displacement vectors correlate with themselves^19^ which is independent of speed. Source data are provided as a Source Data file.

**Figure S5. NHSL1 negatively regulates cell migration in MCF10A cells**

**(A)** MCF10A normal breast epithelial cells were infected with lentiviruses harbouring control or NHSL1 specific shRNAs (shRNA A or B) and also conferring puromycin resistance and selected by puromycin. Stable MCF10A NHSL1 knockdown cell pools were grown until confluent before being scratched and imaged for 12 hours. Still images from live cell imaging showing one still image at the beginning (0 hours, left panel) and at the end of imaging (12 hours, right panel.) Scale bar: 300 μm. **(B)** The area of the scratch was measured at 0 and 12 h. Area closure is shown as percentage increase over control cells. Results are mean ± SEM (error bars), from four independent biological repeats; full circles: control shRNA; empty circles: NHSL1 shRNA A; stars: NHSL1 shRNA B; One-way ANOVA: p=0.0031, F(2,46)=6.573; and Dunnett’s multiple comparisons test: control vs. shRNA A: **, p=0.0033. control vs. shRNA B: **, p=0.0085. See video S4. **(C)** NHSL1 mRNA levels in stable MCF10A cell lines from (A) were measured by quantitative PCR. NHSL1 knockdown reduced mRNA levels to (41.6±8.7)% SEM (shRNA A), or (35.2±6.0)% SEM (shRNA B) compared to scrambled control shRNA. Results are mean ± SEM (error bars); full circles: control shRNA; empty circles: NHSL1 shRNA A; stars: NHSL1 shRNA B; One-way ANOVA: p=0.0024, F(2,6)=19.37; and Dunnett’s multiple comparisons test: control versus shRNA A: **, p=0.0040; control versus shRNA B: **, p=0.0024. Source data are provided as a Source Data file.

**Figure S6. NHSL1 is a novel binding partner of active Rac**

**(A)** Full western blots are shown for Fig. 3G: Western blot showing that dominant active (DA) Rac pulls down NHSL1 using Myc-trap beads from HEK cell lysates expressing Myc-tagged DA Rac1 co-expressed with EGFP-tagged NHSL1 or EGFP only as control. Representative blots from three independent biological repeats. **(B)** Western blot showing that dominant active (DA) Rac pulls down NHSL1 using GFP-trap beads from HEK cell lysates expressing Myc-tagged NHSL1 co-expressed with EGFP-tagged dominant active (DA) Rac1 or EGFP only as control.

**Figure S7. Active Rac binds to two sites in NHSL1**

**(A-B)** Nine overlapping EGFP-tagged subfragments covering fragments 2 and 3 of NHSL1 were generated (A) and expressed in HEK cells along with Myc-tagged DA Rac. After Myc-trap pulldown of Myc-DA-Rac, co-precipitation of EGFP-NHSL1 subfragments 1-9 was detected in a western blot with Myc antibody (B). Representative blots from three independent biological repeats. Source data are provided as a Source Data file.

**Figure S8. NHSL1 co-localises with Abi and the Scar/WAVE complex**

**(A-D)** Endogenous NHSL1 (NHSL1 pAb, magenta) co-localises with Abi1 (cyan) at the very edge of lamellipodia in B16-F1 mouse melanoma cells. **(E-H)** Endogenous NHSL1 (NHSL1 mAb, magenta) co-localises with Scar/WAVE2 (cyan) and at the very edge of lamellipodia in B16-F1 mouse melanoma cells. (A, E) Dual color merge of the insets shown in Fig. 4 (C,E): Scale bar in (A,E): 20 μm. (B-D;F-H) Three line scans (green, magenta, orange) were placed in arbitrary locations perpendicular to the leading edge and the intensity profile plotted; interrupted line: Abi1/WAVE2; continuous line: NHSL1; The Pearson’s correlation coefficient (r^2^) indicating degree of correlation of co-localisation was calculated and is displayed next to each pair of lines.

**Figure S9. Exogenous expression of all tagged components of the Scar/WAVE complex allows Scar/WAVE complex formation**

**(A,B)** Full western blots of experiment shown in Fig. 5A: The Scar/WAVE complex co-immunoprecipitates with NHSL1. HEK cells were transfected with EGFP-NHSL1, and Myc-Pir121, -Nap-1, -WAVE2, -Abi2. NHSL-1 was immunoprecipitated (pAb 4457) from lysates and co-immunoprecipitation tested on a western blot with (A) Myc and reprobed with (B) EGFP antibodies. Representative blots from three independent experiments. **(C)** Negative control for experiment in Fig. 5D: GST-pull downs using purified Glutathione-sepharose coupled GST-fusion proteins of Abi1 and c-Abl SH3 domains or GST alone from HEK cell lysates that were transfected with EGFP. Following GST-pulldown EGFP was detected in a western blot with anti-EGFP antibodies. Ponceau staining of membrane reveals GST or GST-tagged Abi- or Abl-SH3 domains used. Representative blots from three independent experiments. **(D)** HEK cells were transfected with EGFP-tagged Abi1 or Abi1-delta-SH3 or EGFP only as negative control and the remaining Myc-tagged Scar/WAVE components, PIR121, Nap1, Scar/WAVE2, HSPC300. After GFP-trap pulldown, co-precipitation was detected in a western blot with Myc antibody. Representative blots from three independent biological repeats. Source data are provided as a Source Data file.

**Figure S10. Abi SH3 domain binds to two fragments of NHSL1**

**(A)** Far western overlay with purified MBP-tagged full-length Abi1 (MBP-Abi1 full length) or an MBP fusion protein with Abi1 in which the SH3 domain had been deleted (MBP-Abi1-delta-SH3) and MBP as control on a blot of different purified GST-NHSL1 fusion proteins covering the entire length of NHSL1. Representative blots from three independent experiments. Fragments 4 and 5 contain three putative SH3 binding sites. **(B)** Coomassie gel showing GST fragments covering the entire length of the NHSL1 amino acid sequence (see Fig. 1E for fragment sizes and location within NHSL1) and GST only as control which are used in the Far Western Blot in (A).

**Figure S11: Effect of NHSL1 overexpression on cell migration persistence**

**(A)** Western blot showing B16-F1 cells expressing EGFP-NHSL1 wild type (GFP-NHSL1-WT) or the NHSL1 mutant in the Scar/WAVE binding sites (GFP-NHSL1-SW-Mut) or EGFP only as control after selection of the cells with puromycin. The blot was probed with polyclonal antibodies specific for NHSL1 to display the ratio between overexpressed EGFP-NHSL1 (upper band) and endogenous NHSL1 (lower band). Tubulin served as a loading control. **(B-E)** Quantification of cell migration persistence of randomly migrating B16-F1 cells expressing either wild type NHSL1 (NHSL1 WT) or the NHSL1 mutant in the Scar/WAVE complex binding sites (NHSL1 SW Mut) or EGFP alone as control plated on fibronectin. **(B)** Mean Square Displacement (MSD) analysis (log-log plot) of data shown in Fig. 6A. **(C)** Mean track persistence (dt = 3, TR = 4; see materials and methods for calculation). Results are mean ± SEM (error bars), from four independent biological experiments. Each data point represents the mean speed of a cell from a total of number of 106, 104, 108 cells for control, NHSL1 WT, and NHSL1 SW mut, respectively. One-way ANOVA: p<0.0001, F(2,395)=9.573; and Dunnett’s multiple comparisons test: *** p=0.0001; ** p=0.0011. **(D)** The directionality ratio is plotted to explore how it changes over the time of the movie. **(E)** Direction autocorrelation is shown, a measure of how the angle of displacement vectors correlate with themselves^19^ which is independent of speed. **(F)** Still image from live cell imaging showing EGFP-NHSL1 Scar/WAVE binding mutant (EGFP-NHSL1 SW Mut) in B16-F1 cells plated on laminin localises to the leading edge. Representative image from three independent biological repeats. Scale bar: 20 μm. Inset represents a magnified view of the white box. Scale bar in inset: 5 μm. See also related video S7. Source data are provided as a Source Data file.

**Figure S12 NHSL1 negatively regulates cell migration**

**(A,B)** Quantification of persistence of randomly migrating wild type B16-F1 cells or CRISPR 2 cells expressing either NHSL1 (NHSL1 WT) or the NHSL1 mutant in the Scar/WAVE complex binding sites (NHSL1 SW Mut) or Myc alone as control plated on laminin. (A) The directionality ratio is plotted to explore how it changes over the time of the movie. (B) Direction autocorrelation is shown, a measure of how the angle of displacement vectors correlate with themselves^19^ which is independent of speed.

**Figure S13. Active Rac increases Arp2/3 activity - FRET-FLIM controls**

**(A)** Lifetime images of B16-F1 cells expressing the donor ARPC3-mTurq2 and either empty Myc-control plasmid (vector control) or Myc-tagged dominant active Rac (DA Rac). Warm colours indicate short lifetimes of the donor ARPC3-mTurq2 only as control. Representative images from 4 independent biological repeats are shown. Scale bar: 20 μm. **(B)** FRET efficiency histograms from the same representative B16-F1 cells expressing control Myc plasmid (black circles) and Myc-tagged dominant active Rac (DA Rac) (brown upside-down triangles) cells expressing the donor ARPC3-mTurq2. **(C-E)** Quantification of FRET efficiency controls used to verify that any change in efficiency was a result of real changes in Arp2/3 activity and not skewed by, or an artefact of, the environment, vectors, or the FRET pair itself. The weighted mean average for each cell was calculated from the FRET efficiency histograms and were used instead of the normal mean in order to better represent the true FRET efficiency. Data is shown as weighted average mean ± SEM (error bars). Quantification from 4 independent biological repeats for B16-F1 cells expressing (C) control Myc plasmid and the donor ARPC3-mTurq2 (black circles; n=30 cells) or Myc DA Rac and the donor ARPC3-mTurq2 (brown upside-down triangles; n=30 cells); ns=0.9886; t=0.01436, df=58, Unpaired, two-tailed t-test. (D) control Myc plasmid and the donor ARPC3-mTurq2 (black circles; n=30 cells) or control Myc plasmid and the Arp2/3 biosensor (orange circles; n=31 cells), ns=0.6790, t=0.4158, df=59, Unpaired, two-tailed t-test and (E) Myc DA Rac and the donor ARPC3-mTurq2 (brown upside-down triangles; n=30 cells) or Myc DA Rac and the Arp2/3 biosensor (lilac upside-down triangles; n=26 cells), ****, p<0.0001, t=7.379, df=54, Unpaired, two-tailed t-test. Source data are provided as a Source Data file.

**Figure S14. NHSL1 negatively regulates Arp2/3 activity - FRET-FLIM controls**

**(A)** Lifetime images of wild type B16-F1, NHSL1 CRISPR clone 2 and NHSL1 CRISPR clone 21 cells expressing the donor ARPC3-mTurq2. Warm colours indicate short lifetimes of the donor mTurq2 only as control. Representative images from 4 independent biological repeats are shown. Scale bar: 20 μm. **(B)** FRET efficiency histograms from the same representative wild type B16-F1 (orange circles), NHSL1 CRISPR clone 2 (magenta squares) and NHSL1 CRISPR clone 21 (green triangles) cells expressing the donor ARPC3-mTurq2. **(C-F)** Quantification of FRET efficiency controls used to verify that any change in efficiency was a result of real changes in Arp2/3 activity and not skewed by, or an artefact of, the environment, vectors, or the FRET pair itself. Quantification from 4 independent biological repeats is for B16-F1 cells expressing (C) donor fluorophore only controls for each biological condition and control Myc plasmid (black circles; n=22 cells), NHSL1 CRISPR2 (black squares; n=22 cells), and NHSL1 CRISPR21 (black triangles; n=16 cells), One-way ANOVA and Tukey’s multiple comparisons test: p=0.9859, F(2,57)=0.01420, control vs. CRISPR2: ns p=0.9991; control vs. CRISPR21: ns p=0.9913; CRISPR2vs. CRISPR21: ns p=0.9853. (D) control Myc plasmid with donor-only fluorophore (black circles; n=22 cells) and donor-acceptor fluorophore (Arp2/3 biosensor; orange circles; n=19 cells), Mann-Whitney test, two tailed: ns: p=0.5957. (E) NHSL1 CRISPR2 with donor-only fluorophore (black circles) and donor-acceptor fluorophore (Arp2/3 biosensor; magenta squares), (F) NHSL1 CRISPR21 with donor-only fluorophore (black circles; n=22 cells) and donor-acceptor fluorophore (Arp2/3 biosensor; green triangles; n=21 cells), Mann-Whitney test, two tailed: * p=0.0126. The weighted mean for each cell was calculated from the FRET efficiency histograms and were used instead of the normal mean in order to better represent the true FRET efficiency. Results are an average of the weighted mean ± SEM (error bars). **(G)** Quantification of average cellular FRET efficiency which represents Arp2/3 activity from five independent biological repeats from wild type B16-F1 cells expressing Myc alone as control (black circles; 37 cells) or CRISPR 2 cells expressing either the NHSL1 mutant in the Scar/WAVE complex binding sites (NHSL1 SW Mut, blue diamonds; 33 cells) or NHSL1 (NHSL1 WT, red crosses; 34 cells) or Myc alone as control (pink squares; 36 cells) plated on laminin after selection using a bicistronic expression plasmid also conferring resistance to blasticidin to ensure that all cells analysed expressed NHSL1. Data points are the weighted average means for each cell ± SEM (error bars), calculated from the FRET efficiency histograms. One-way ANOVA: p=0.0001; F(3,136)=7.421; and Dunnett’s multiple comparisons test: CRISPR2 vs. WT control: *, p=0.0324; NHSL1 CRISPR2 vs. Rescue Myc-NHSL1 SW Mut: ****, p<0.0001; NHSL1 CRISPR2 vs. Rescue Myc-NHSL1 WT: **, p=0.0052. **(H)** Quantification of FRET efficiency which represents Arp2/3 activity in the approximate area of the lamellipodium from 4 independent biological repeats from control: (orange circles; n=15 cells); NHSL1 CRISPR 2: (magenta squares; n=18 cells); NHSL1 CRISPR 21: (green triangles; n=9 cells). Again, data points are the weighted mean for each cell calculated from the FRET efficiency histograms. Results are the average weighted mean ± SEM (error bars), Kruskal-Wallis: p=0.2966 and Dunn’s multiple comparisons test: Control vs. CRISPR 2: ns, p=0.6914 and Control vs. CRISPR 21: ns, p=0.2502. Source data are provided as a Source Data file.

**Figure S15. NHSL1 knockout reduces whole cell Scar/WAVE2, ArpC2, and F-actin levels**

**(A,B)** Example images for immunofluorescence analysis of (A) ARPC2 for whole cell Arp2/3 complex intensity quantification in (C) and Fig. 8D and (B) F-actin (Alexa488-Phalloidin) in Fig. 8E. **(C)** Quantification of whole cell Arp2/3 intensity after background subtraction. One-way ANOVA: p=0.0010; F(2,192)=7.202, Dunnett’s multiple comparisons test: *** p=0.0004; ns p=0.0758; in wild-type (n=75 cells, control), NHSL1 CRISPR 2 (n=60 cells) and NHSL1 CRISPR 21 (n=60 cells) B16-F1 cells plated on laminin and stained with anti-ARPC2 (subunit of Arp2//3 complex) antibodies; Results are mean ± SEM (error bars), three independent biological repeats. **(D)** Western blot comparing Scar/WAVE2 and ARPC2 expression levels in wild type control B16-F1 cell lysates with NHSL1 CRISPR 2 or 21 cell lysates. Beta-Tubulin serves as the loading control. **(E,F)** Quantification of the western blot in (D) for Scar/WAVE2 in (E) and for ARPC2 in (F) normalised to the Tubulin loading control; One-way ANOVA: (E) p=0.0008; F(2,9)=17.63; (F) p=0.0017; F(2,6)=22.19; Dunnett’s multiple comparisons test: control vs. CRISPR 2: (E) *** p=0.0004; (F) * p=0.0163; control vs. CRISPR 21(E) * p=0.0110; (F) ns p=0.0502.

**Figure S16. NHSL1 knockout reduces whole cell F-actin levels**

**(A)** Example images for whole cell F-actin (Alexa488-Phalloidin) intensity quantification in Fig. 8C. **(B and C)** Quantification of F-Actin content in HEK293 cells that were transfected with (B) EGFP or EGFP-NHSL1 and (C) control shRNA or two independent NHSL1-specific shRNA’s (see Fig. S5C), which were puromycin selected and replated for 1 hour on fibronectin coated coverslips. Three independent experiments were used for this quantification. (B) EGFP-NHSL1 overexpression increased F-Actin content by 28.8±5.7% SEM, EGFP (black circles): 134 cells total in all 3 experiments; EGFP-NHSL1 (red crosses): 128 cells total in all 3 experiments; Each dot or cross represents the mean of one independent biological repeats; t=5.054, df=4, **, p=0.0072, unpaired, two-tailed t-test. (C) Box and whiskers plot: Box 25 and 75 percentile; whiskers: 10 and 90 percentile; line: median; cross: mean; NHSL1 knockdown decreased F-Actin content by 10.9±0.7% SEM (shRNA A) or 12.2±0.7% SEM (shRNA B), *** = p≤0.001, Kruskal-Wallis test: Kruskal Wallis statistics: 164.3; p<0.0001 and Dunn’s multiple comparisons test: scrambled shRNA vs. shRNA A: ****, p<0.0001 and scrambled shRNA vs. shRNA B: ****, p<0.0001; scrambled shRNA: 1811 cells; shRNA A: 2196 cells; shRNA B: 2148 cells from three independent biological repeats.

**Figure S17 NHSL1 negatively regulates cell area and lamellipodial length**

**(A,B)** Quantification of total cell area and lamellipodia length from first frame of a movie of LifeAct-EGFP expressing wild type B16-F1 or NHSL1 CRISPR 2 or 21 cells plated on laminin. (A) Quantification of total cell area, ****, p<0.0001; *, p=0.0160; F(2,42)=12.86; (B) Quantification of lamellipodia length: The length was quantified using the Fiji plugin “measure_ROI” (http://www.optinav.info/Measure-Roi.htm: <u>Measure_Roi_Curve.java</u>) which measures the length of curved objects (see materials and methods for details). See Fig. 8 (A) for an example B16-F1 cell shown indicating the definition of length. ****, p<0.0001; ns, p=0.1193; F(2,42)=9.739. Results are mean ± SEM (error bars), four independent biological repeats, n = 15 cells. One-way ANOVA, Dunnett’s multiple comparisons test. **(C)** Quantification of number of microspikes per length of lamellipodium; One-way ANOVA: F(2,42)=0.8509; Dunnett’s multiple comparisons test: ns (CRISPR2), p=0.3400; ns(CRISPR21), p=0.6206; in wild type (control), NHSL1 CRISPR 2, and NHSL1 CRISPR 21 B16-F1 cells plated on laminin expressing LifeAct-EGFP; Results are mean ± SEM (error bars), four independent biological repeats, n = 15 cells. **(D)** Quantification of lamellipodia width, ns (CRISPR2), p=0.1173; ns(CRISPR21), p=0.4013; F(2,42)=1.89, One-way ANOVA, Dunnett’s multiple comparisons test. Source data are provided as a Source Data file. **(E)** Example images for lamellipodia LifeAct-EGFP intensity quantification in Fig. 8F.

**Figure S18. Analysis of segmented lamellipodium speed and two Abi binding sites are conserved in the NHS protein family**

**(A)** Quantification of the average speed of the centroid of the segmented lamellipodium from a wild type cell (WT), NHSL1 knockout cell (NHSL1 CRISPR2), and NHSL1 knockout cells rescued with a wild type NHSL1 construct (NHSL1 CRISPR 2 Rescue Myc-NHSL1 WT) and with a mutant NHSL1 construct (NHSL1 CRISPR 2 Rescue Myc-NHSL1 SW Mut). Results are mean ± SEM; n = 41, 32, 35, and 36 for the respective conditions (left to right on x-axis), from 6 independent biological repeats. One-way ANOVA: p=0.1948; F(3,140)=1.589; and Tukey’s multiple comparisons test: Control vs. CRISPR 2: p=0.2755; Control vs. NHSL1 SW Mut Rescue: p=0.3091; Control vs. NHSL1 WT Rescue: p=0.3399; CRISPR 2 vs. NHSL1 SW Mut Rescue: p=0.9996; CRISPR 2 vs. NHSL1 WT Rescue: p=0.9981; NHSL1 SW Mut Rescue vs. NHSL1 WT Rescue: p=0.9999. See video S8. **(B)** The conservation on the amino acid level in the NHS protein family is shown as percent amino acid identity as calculated from a Clustal Omega alignment (DNASTAR MegAlignPro) from the UniProt canonical amino acid sequences of each NHS family member. The UniProt identifier of each the sequence of each NHS family member used is displayed in the table. **(C,D)** HHM logo of conservation of Abi SH3 binding sites in the NHS protein family (from PFAM PF15273). Source data are provided as a Source Data file.

## Supplemental movie legends

**Video S1. NHSL1 localises to the very edge of lamellipodia**

This movie shows that either N- or C-terminally EGFP-tagged NHSL1 expressed in B16-F1 cells plated on laminin localises to the very edge of protruding lamellipodia. Representative movie shown from three independent biological repeats. Cells were imaged every 10 seconds for the indicated times in minutes and seconds by wide-field time-lapse video microscopy using an IX 81 microscope (Olympus). Scale bar: 20 μm.

**Video S2. EGFP-NHSL1 localisation to the leading edge is not due to space filling of the fluorescent protein**

This movie shows EGFP-NHSL1 co-expressed with mScarlet-I in B16-F1 cells to control for detection at the leading edge due to space filling of the fluorescent protein. The subtracted movie is generated from the subtraction of the signal of the mScarlet-I from the NHSL1-EGFP signal for each frame. Representative movie shown from three independent biological repeats. Cells were imaged every 10 seconds for the indicated times in minutes and seconds by wide-field time-lapse video microscopy using an IX 81 microscope (Olympus). Scale bar: 20 μm.

**Video S3. NHSL1 co-localise with Lpd at the edge of lamellipodia**

This movie shows that NHSL1-EGFP co-expressed with mScarlet-I-tagged Lpd co-localise together at the edge of lamellipodia in B16-F1 cells plated on laminin. Representative movie shown from three independent experiments. Cells were imaged every 10 seconds for the indicated times in minutes and seconds by wide-field time-lapse video microscopy using an IX 81 microscope (Olympus). Scale bar: 20 μm.

**Video S4. NHSL1 knockdown increases cell migration**

This movie shows that directional migration into a scratch wound was significantly increased in MCF10A normal breast epithelial cells in which NHSL1 expression had been knocked-down for both shRNAs tested. Confluent monolayers were scratch-wounded and imaged every 5 min for 12 hr by time-lapse phase contrast wide-field microscopy using an IX 81 microscope (Olympus). Scale bar: 300 μm.

**Video S5. Specific NHSL1 fragments localise to the leading edge and to vesicular structures**

These movies show the localisation of four EGFP-tagged fragments covering the entire length of NHSL1 expressed in B16-F1 cells. Fragment 2 and 3 localise to the very edge of lamellipodia similar to full length NHSL1 (see video S1). EGFP alone served as the negative control. In addition, fragment 1 and 3 were detected at vesicular structures. Representative movies shown from three independent experiments. Cells were imaged every 10 seconds for the indicated times in minutes and seconds by wide-field time-lapse video microscopy using an IX 81 microscope (Olympus). Scale bar: 20 μm.

**Video S6. NHSL1 co-localise with components of the Scar/WAVE complex**

These movies show NHSL1-EGFP co-expressed with mScarlet-I-tagged Abi1 or Nap1 co-localise at the leading edge of cells in B16-F1 cells plated on laminin. Representative movies shown from three independent experiments. Cells were imaged every 10 seconds for the indicated times in minutes and seconds by wide-field time-lapse video microscopy using an IX 81 microscope (Olympus). Scale bar: 20 μm.

**Video S7. The NHSL1 Scar/WAVE binding mutant localises to the very edge of lamellipodia**

This movie shows that the NHSL1 Scar/WAVE binding mutant localises to the very edge of lamellipodia like wild type EGFP-NHSL1. Representative movie shown from three independent experiments. Cells were imaged every 10 seconds for the indicated times in minutes and seconds by wide-field time-lapse video microscopy using an IX 81 microscope (Olympus). Scale bar: 20 μm.

**Video S8. NHSL1 reduces actin retrograde flow speed and F-actin assembly rate**

This movie shows that NHSL1 reduces actin retrograde flow speed and F-actin assembly rate. Shown is a segmented lamellipodium from a wild type cell (WT), NHSL1 knockout cell (NHSL1 CRISPR2), and NHSL1 knockout cells rescued with a wildtype NHSL1 construct (NHSL1 CRISPR 2 Rescue Myc-NHSL1 WT) and with a mutant NHSL1 construct (NHSL1 CRISPR 2 Rescue Myc-NHSL1 SW Mut). Left panels show segmented Airyscan movies; right panels show the corresponding movies from the PIV interpolation. Each colourmap has the same range (0-3 μm/min) and distance between vector arrows (1 μm); warmer colours represent faster flow, vector arrows are all of unit length. Representative movie shown from six independent biological repeats. Cells were imaged every 3.22 seconds for the indicated times in minutes and seconds by confocal time-lapse video microscopy using a Zeiss Airyscan microscope (Zeiss). Scale bar: 10 μm.

**Video S9. NHSL1 CRISPR cells display a lamellipodium which is less interrupted by retractions**

This movie shows that NHSL1 CRISPR cells displayed a lamellipodium which is less interrupted by retractions. B16-F1 cells were transfected with LifeAct-EGFP and plated onto laminin. Representative movie shown from four independent experiments. Cells were imaged every 5 seconds for the indicated times in minutes and seconds by confocal time-lapse video microscopy using a Zeiss Airyscan microscope (Zeiss). The cell outline was automatically segmented in each frame of the movies, protrusion vectors at each pixel along the cell edge calculated (red arrows shown in lower movie panels), the cell edge binarised and areas of protrusions along the cell edge indicated by a blue line and areas of retraction by a red line (shown in upper movie panels). Scale bar: 20 μm.

**Video S10. Cells overexpressing Myc-NHSL1 display a lamellipodium which is more interrupted by retractions**

This movie shows that the protruding lamellipodium is more interrupted by retractions in cells overexpressing Myc-NHSL1. B16-F1 cells were transfected with Myc-NHSL1-IRES-Puro-T2A-LifeAct-EGFP or empty Myc-IRES-Puro-T2A-LifeAct-EGFP, selected with puromycin and plated onto laminin. Representative movie shown from four independent experiments. Cells were imaged every 5 seconds for the indicated times in minutes and seconds by confocal time-lapse video microscopy using a Zeiss Airyscan microscope (Zeiss). The cell outline was automatically segmented in each frame of the movies, protrusion vectors at each pixel along the cell edge calculated (red arrows shown in lower movie panels), the cell edge binarised and areas of protrusions along the cell edge indicated by a blue line and areas of retraction by a red line (shown in upper movie panels). Scale bar: 20 μm.

