## Supplemental Figures for "Nance-Horan Syndrome-like 1 protein negatively regulates Scar/WAVE-Arp2/3 activity and inhibits lamellipodia stability and cell migration"

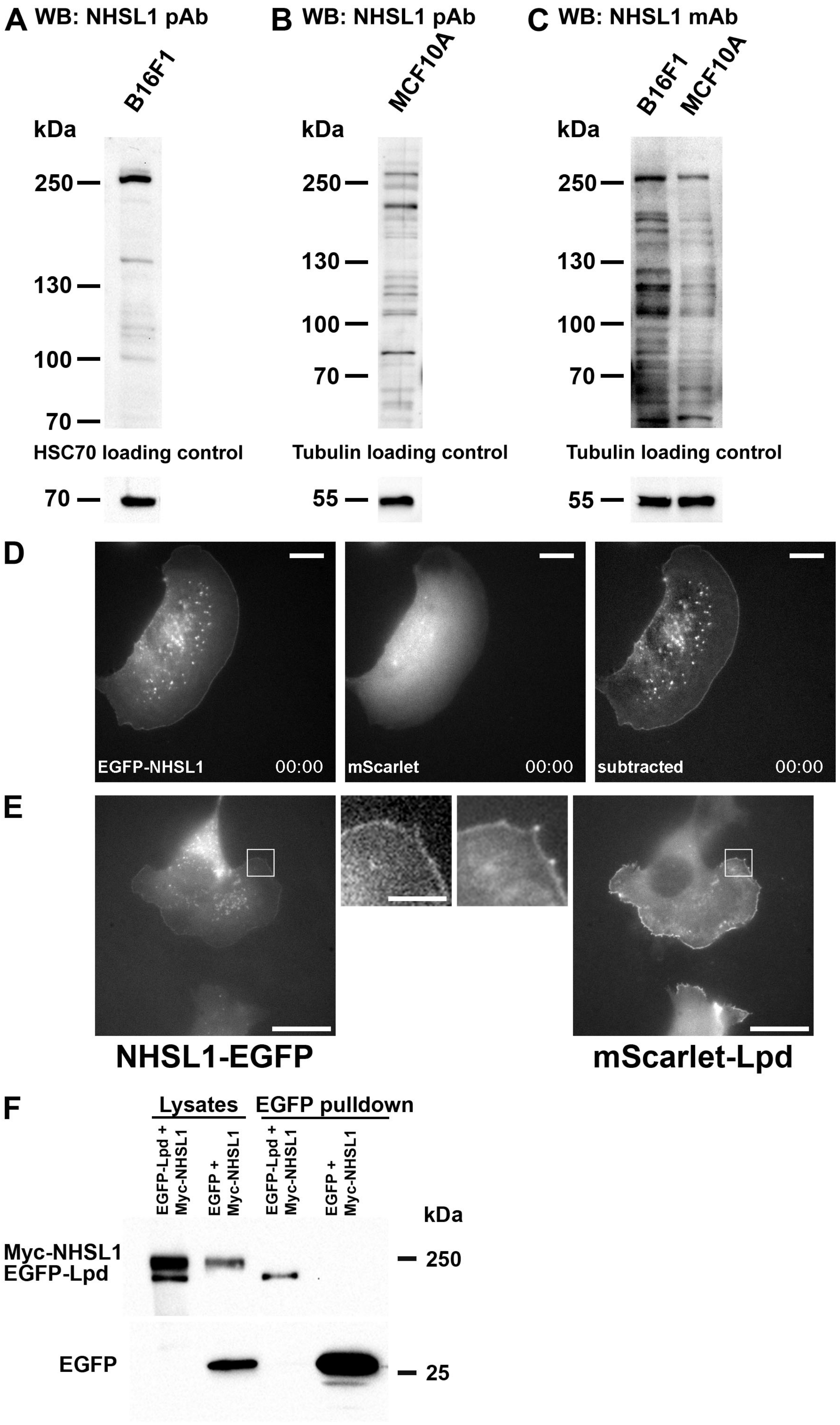
A WB: NHSL1 pAb
B16F1
kDa
250
130
100
70
HSC70 loading control
70
B WB: NHSL1 pAb
MCF10A
kDa
250
130
100
70
Tubulin loading control
55
C WB: NHSL1 mAb
B16F1
MCF10A
kDa
250
130
100
70
Tubulin loading control
55
D
EGFP-NHSL1
00:00
mScarlet
00:00
subtracted
00:00
E
NHSL1-EGFP
mScarlet-Lpd
F
Lysates
EGFP pulldown
EGFP-Lpd + Myc-NHSL1
EGFP + Myc-NHSL1
EGFP-Lpd + Myc-NHSL1
EGFP + Myc-NHSL1
kDa
Myc-NHSL1
EGFP-Lpd
250
EGFP
25

### Figure S1

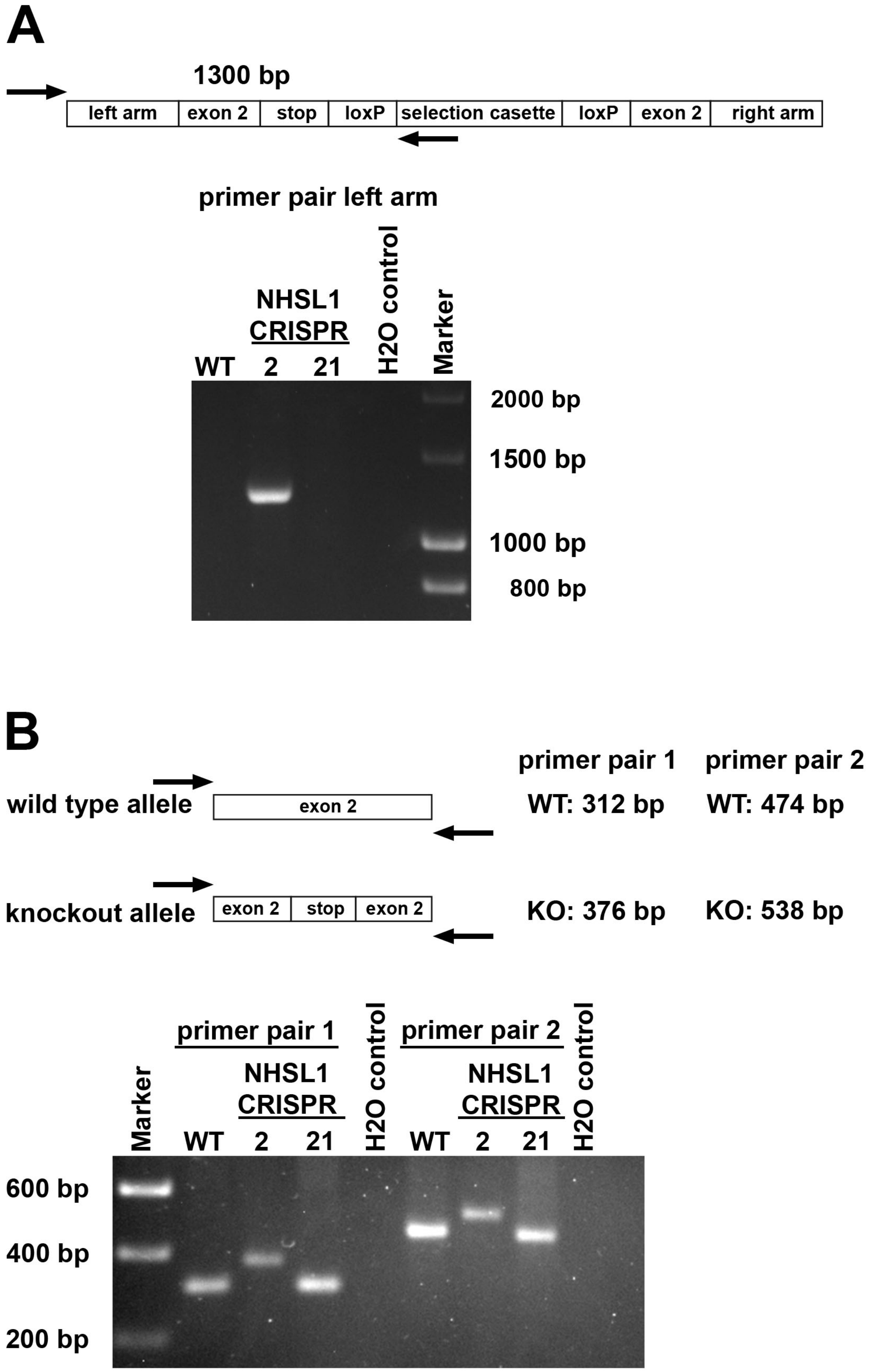
A
1300 bp
left arm
exon 2
stop
loxP
selection casette
loxP
exon 2
right arm
primer pair left arm
NHSL1
CRISPR
WT
2
21
H2O control
Marker
2000 bp
1500 bp
1000 bp
800 bp
B
wild type allele
exon 2
knockout allele
exon 2
stop
exon 2
primer pair 1
primer pair 2
WT: 312 bp
WT: 474 bp
KO: 376 bp
KO: 538 bp
Marker
primer pair 1
NHSL1
CRISPR
WT
2
21
H2O control
primer pair 2
NHSL1
CRISPR
WT
2
21
H2O control
600 bp
400 bp
200 bp

### Figure S2

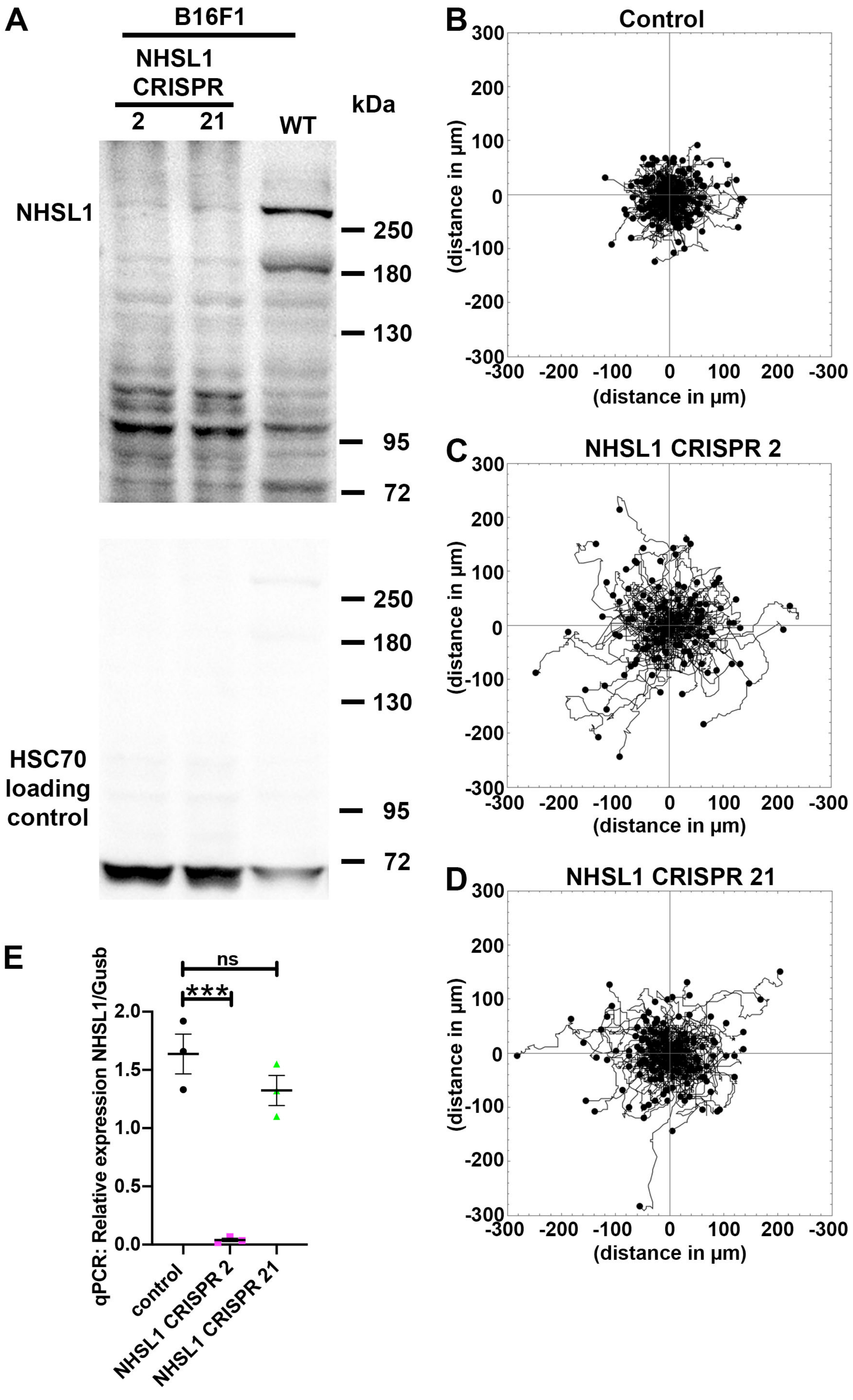
A
B16F1
NHSL1
CRISPR
2
21
WT
kDa
NHSL1
250
180
130
95
72
HSC70
loading
control
250
180
130
95
72
B
Control
C
NHSL1 CRISPR 2
D
NHSL1 CRISPR 21
300
200
100
0
-100
-200
-300
(distance in µm)
-300 -200 -100 0 100 200 -300
(distance in µm)
E
qPCR: Relative expression NHSL1/Gusb
2.0
1.5
1.0
0.5
0.0
ns
***
control
NHSL1 CRISPR 2
NHSL1 CRISPR 21

### Figure S3

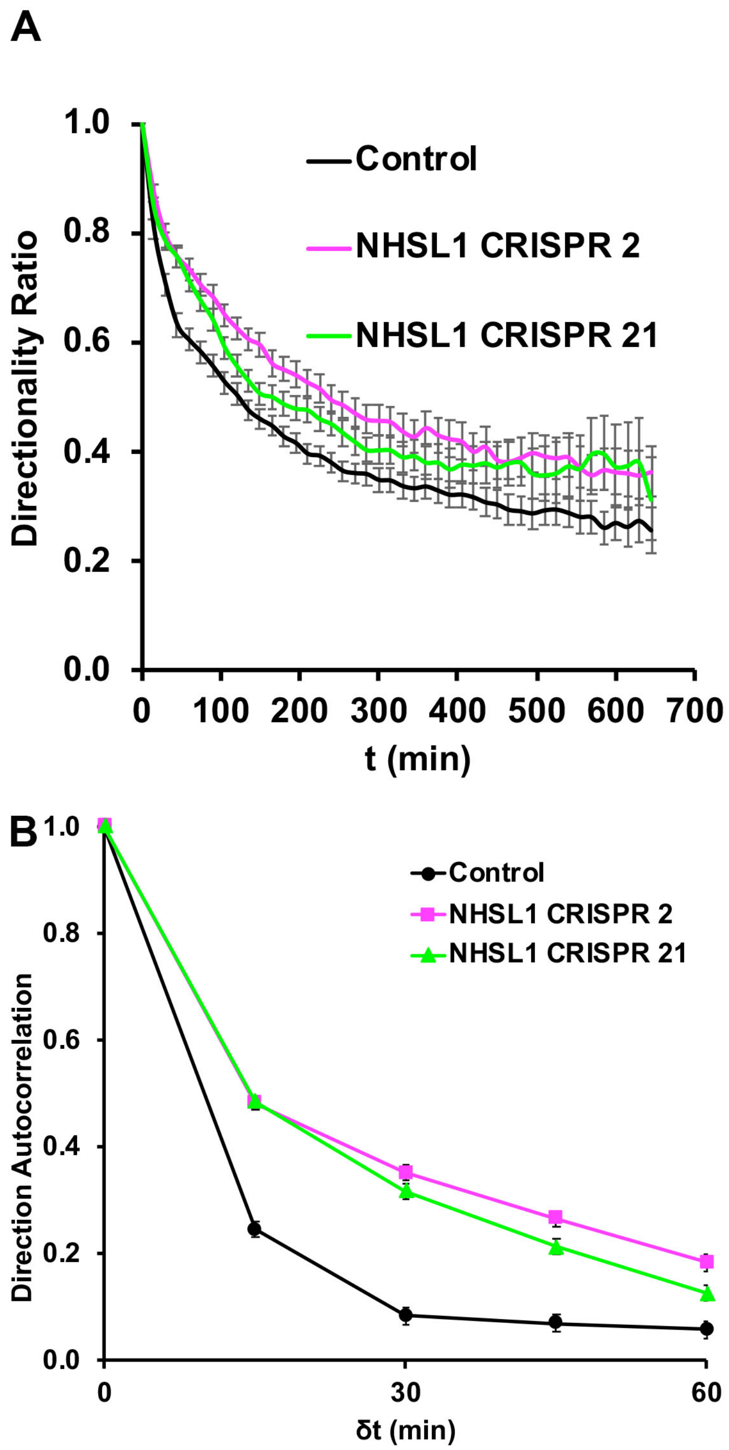
A
1.0
0.8
0.6
0.4
0.2
0.0
Directionality Ratio
Control
NHSL1 CRISPR 2
NHSL1 CRISPR 21
0 100 200 300 400 500 600 700
t (min)
B
1.0
0.8
0.6
0.4
0.2
0.0
Direction Autocorrelation
Control
NHSL1 CRISPR 2
NHSL1 CRISPR 21
0 30 60
δt (min)

### Figure S4

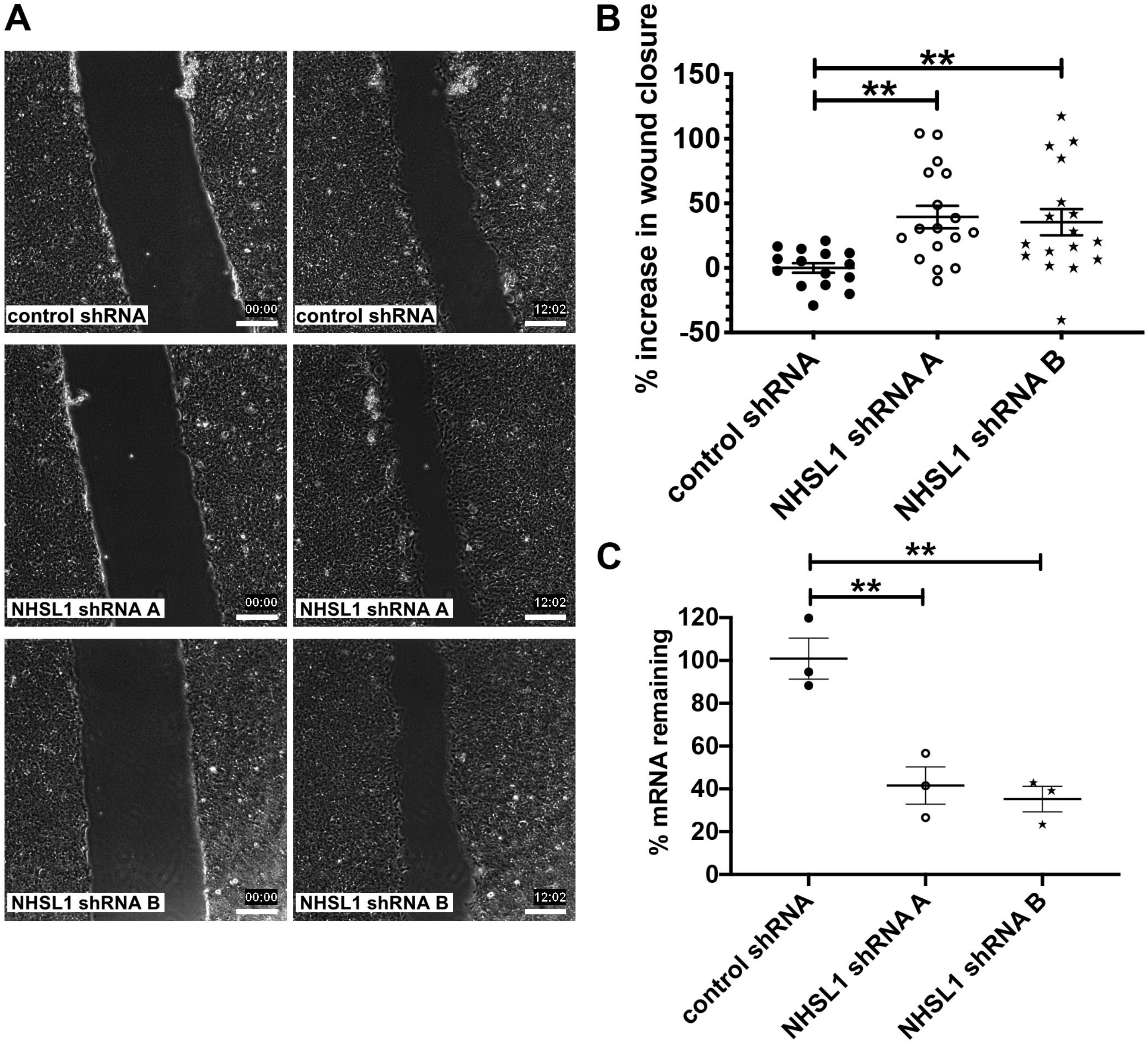
A
control shRNA
00:00
control shRNA
12:02
NHSL1 shRNA A
00:00
NHSL1 shRNA A
12:02
NHSL1 shRNA B
00:00
NHSL1 shRNA B
12:02
B
% increase in wound closure
150
100
50
0
-50
**
**
control shRNA
NHSL1 shRNA A
NHSL1 shRNA B
C
% mRNA remaining
120
100
80
60
40
20
0
**
**
control shRNA
NHSL1 shRNA A
NHSL1 shRNA B

### Figure S5

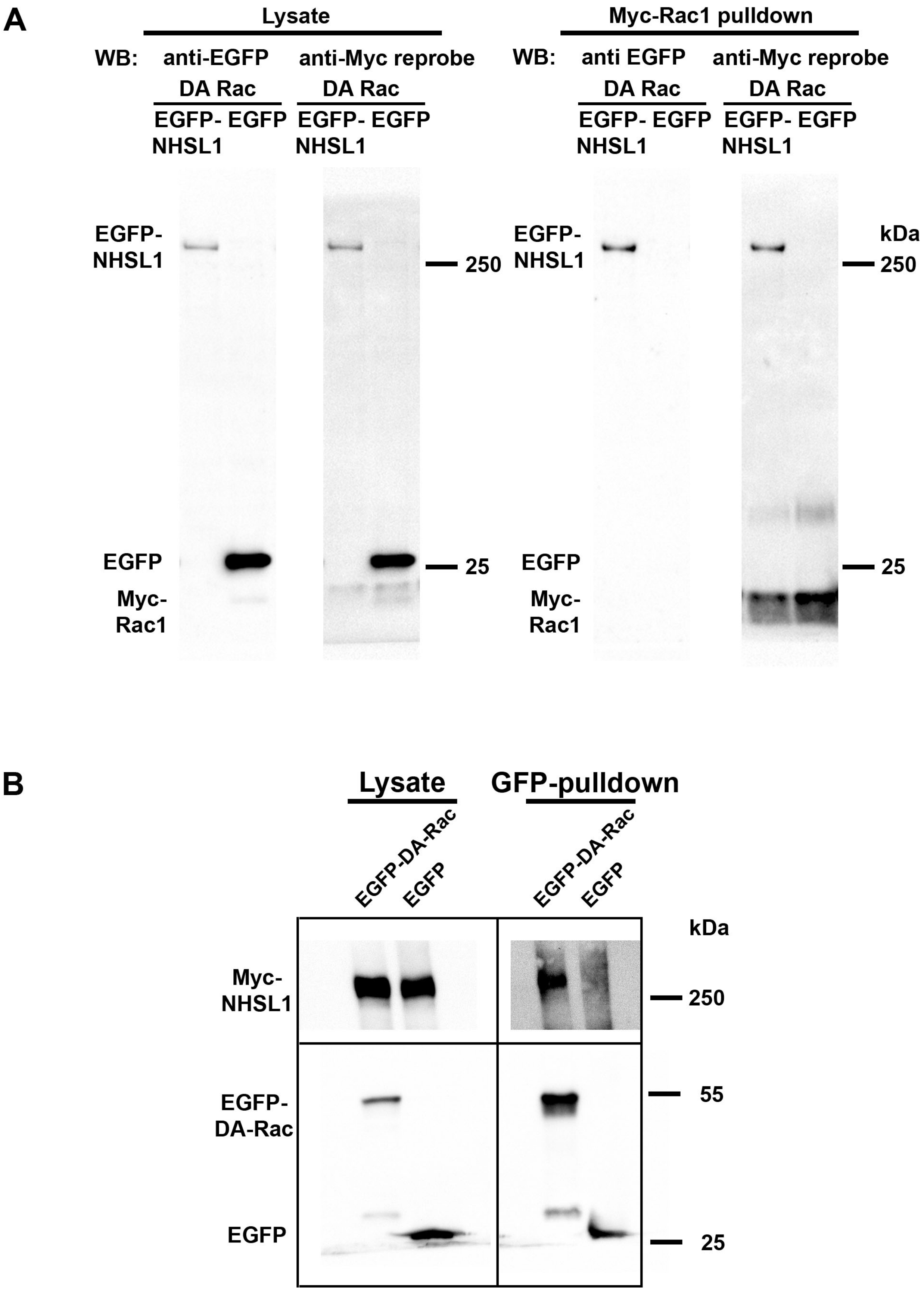
A
Lysate
Myc-Rac1 pulldown
WB:
anti-EGFP
anti-Myc reprobe
anti EGFP
DA Rac
EGFP-
NHSL1
EGFP
EGFP-
NHSL1
EGFP
NHSL1
Myc-
Rac1
kDa
250
25
B
Lysate
GFP-pulldown
EGFP-DA-Rac
Myc-
NHSL1
EGFP-
DA-Rac
55

### Figure S6

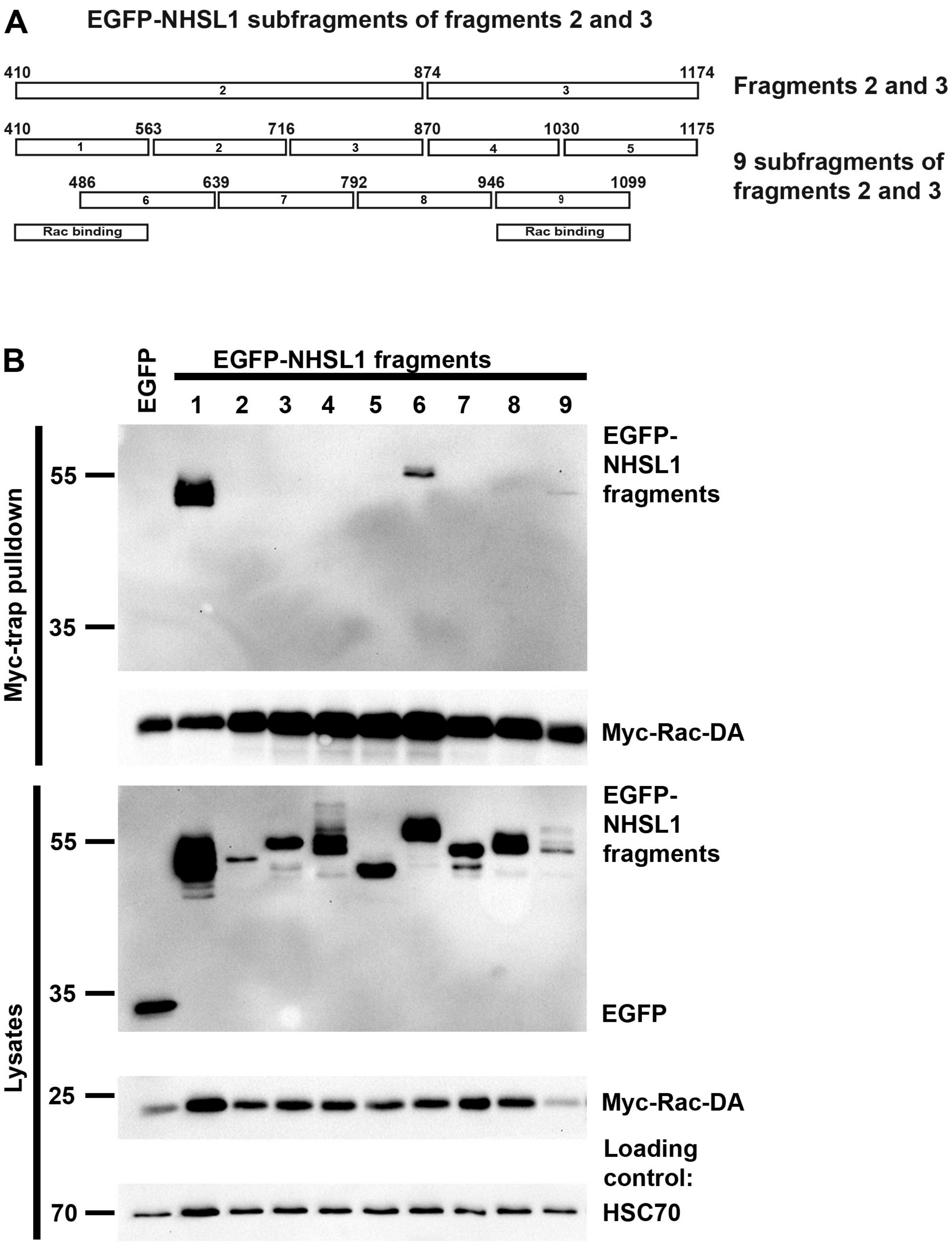
A
EGFP-NHSL1 subfragments of fragments 2 and 3
410
874
1174
2
3
Fragments 2 and 3
410
563
716
870
1030
1175
1
2
3
4
5
486
639
792
946
1099
6
7
8
9
9 subfragments of fragments 2 and 3
Rac binding
Rac binding
B
EGFP
EGFP-NHSL1 fragments
1
2
3
4
5
6
7
8
9
Myc-trap pulldown
55
35
EGFP-NHSL1 fragments
Myc-Rac-DA
Lysates
55
35
EGFP-NHSL1 fragments
EGFP
25
Myc-Rac-DA
Loading control:
70
HSC70

### Figure S7

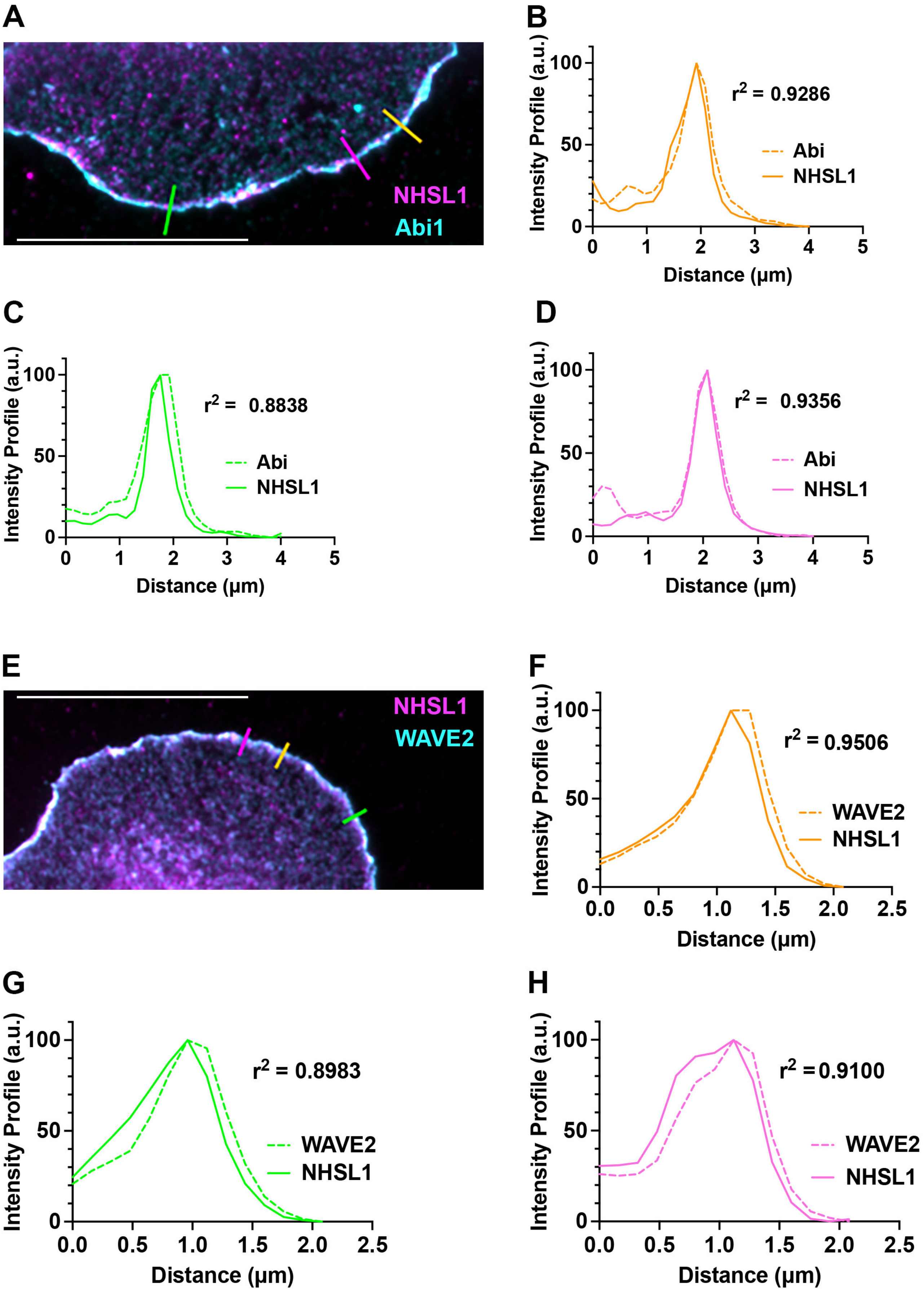
A
NHSL1
Abi1
B
Intensity Profile (a.u.)
Distance (μm)
r2 = 0.9286
Abi
NHSL1
C
r2 = 0.8838
Abi
NHSL1
D
r2 = 0.9356
Abi
NHSL1
E
NHSL1
WAVE2
F
r2 = 0.9506
WAVE2
NHSL1
G
r2 = 0.8983
WAVE2
NHSL1
H
r2 = 0.9100
WAVE2
NHSL1

### Figure S8

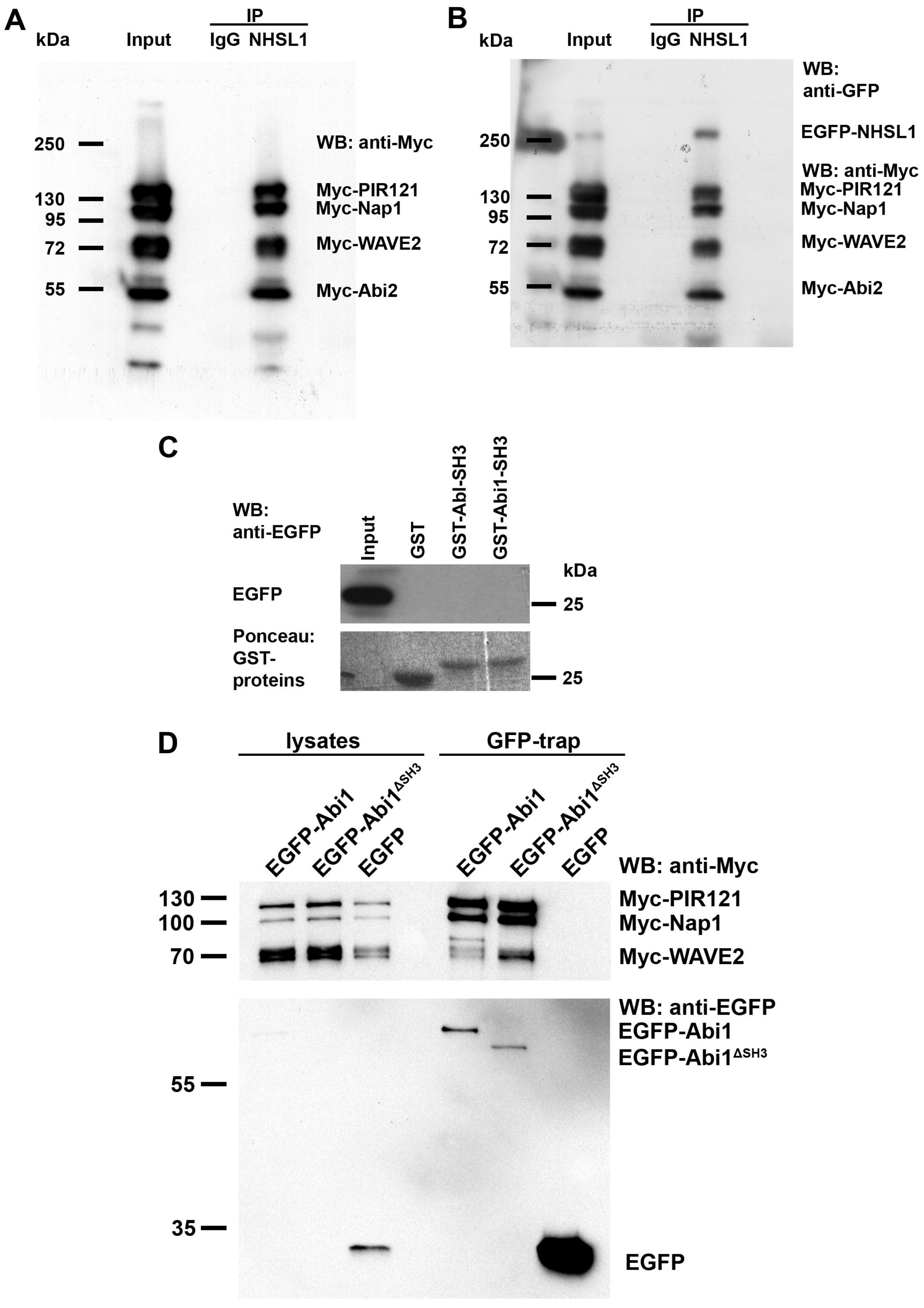
A
kDa
Input
IP
IgG
NHSL1
250
130
95
72
55
WB: anti-Myc
Myc-PIR121
Myc-Nap1
Myc-WAVE2
Myc-Abi2
B
kDa
Input
IP
IgG
NHSL1
WB: anti-GFP
EGFP-NHSL1
250
WB: anti-Myc
130
95
72
55
Myc-PIR121
Myc-Nap1
Myc-WAVE2
Myc-Abi2
C
WB: anti-EGFP
Input
GST
GST-Abl-SH3
GST-Abi1-SH3
kDa
EGFP
25
Ponceau: GST-proteins
25
D
lysates
GFP-trap
EGFP-Abi1
EGFP-Abi1ΔSH3
EGFP
EGFP-Abi1
EGFP-Abi1ΔSH3
EGFP
WB: anti-Myc
130
100
70
Myc-PIR121
Myc-Nap1
Myc-WAVE2
WB: anti-EGFP
EGFP-Abi1
EGFP-Abi1ΔSH3
55
35
EGFP

### Figure S9

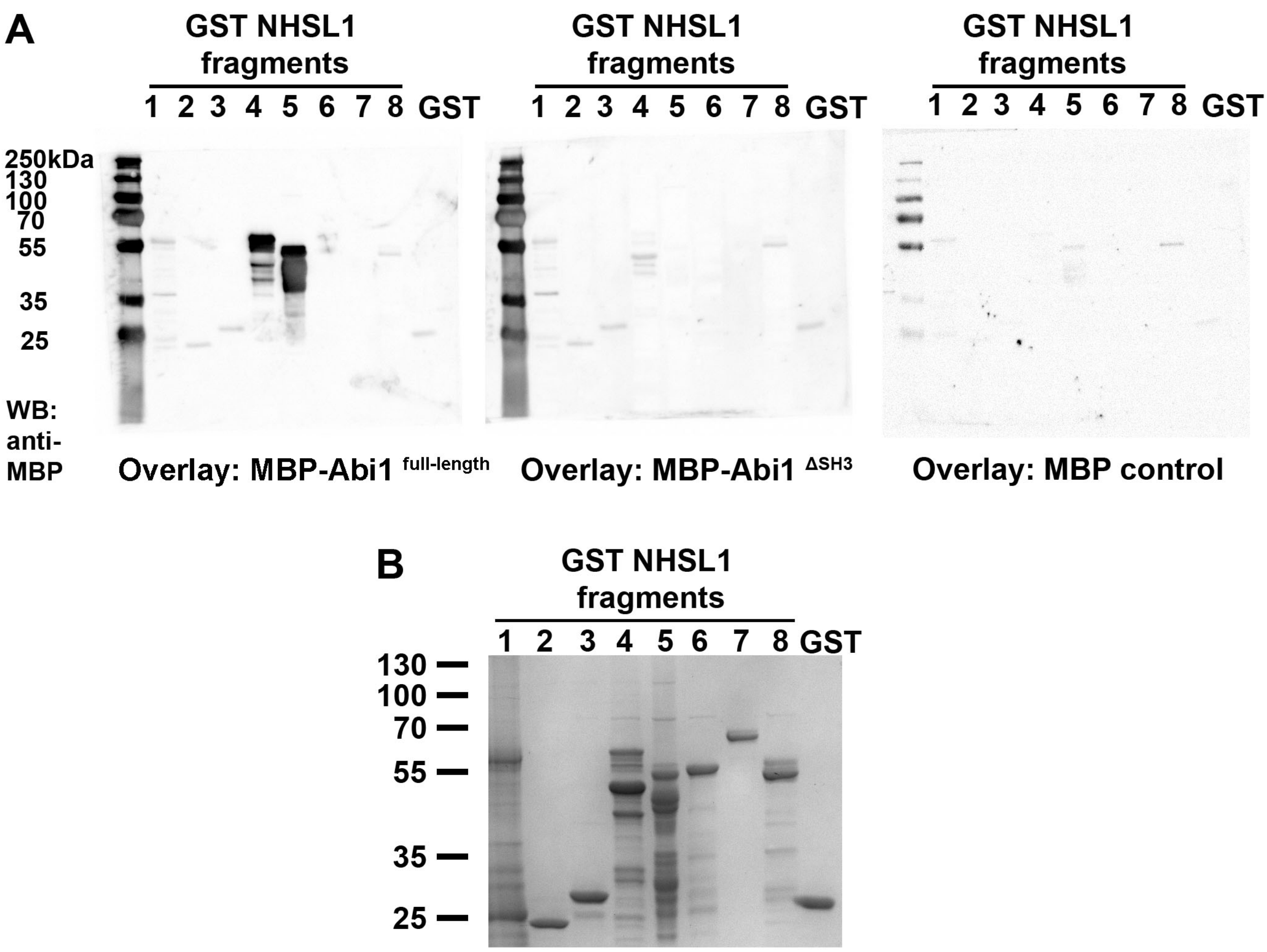
A
GST NHSL1 fragments
1 2 3 4 5 6 7 8 GST
250kDa
130
100
70
55
35
25
WB: anti-MBP
Overlay: MBP-Abi1 full-length
Overlay: MBP-Abi1 ΔSH3
Overlay: MBP control
B
GST NHSL1 fragments
1 2 3 4 5 6 7 8 GST
130
100
70
55
35
25

### Figure S10

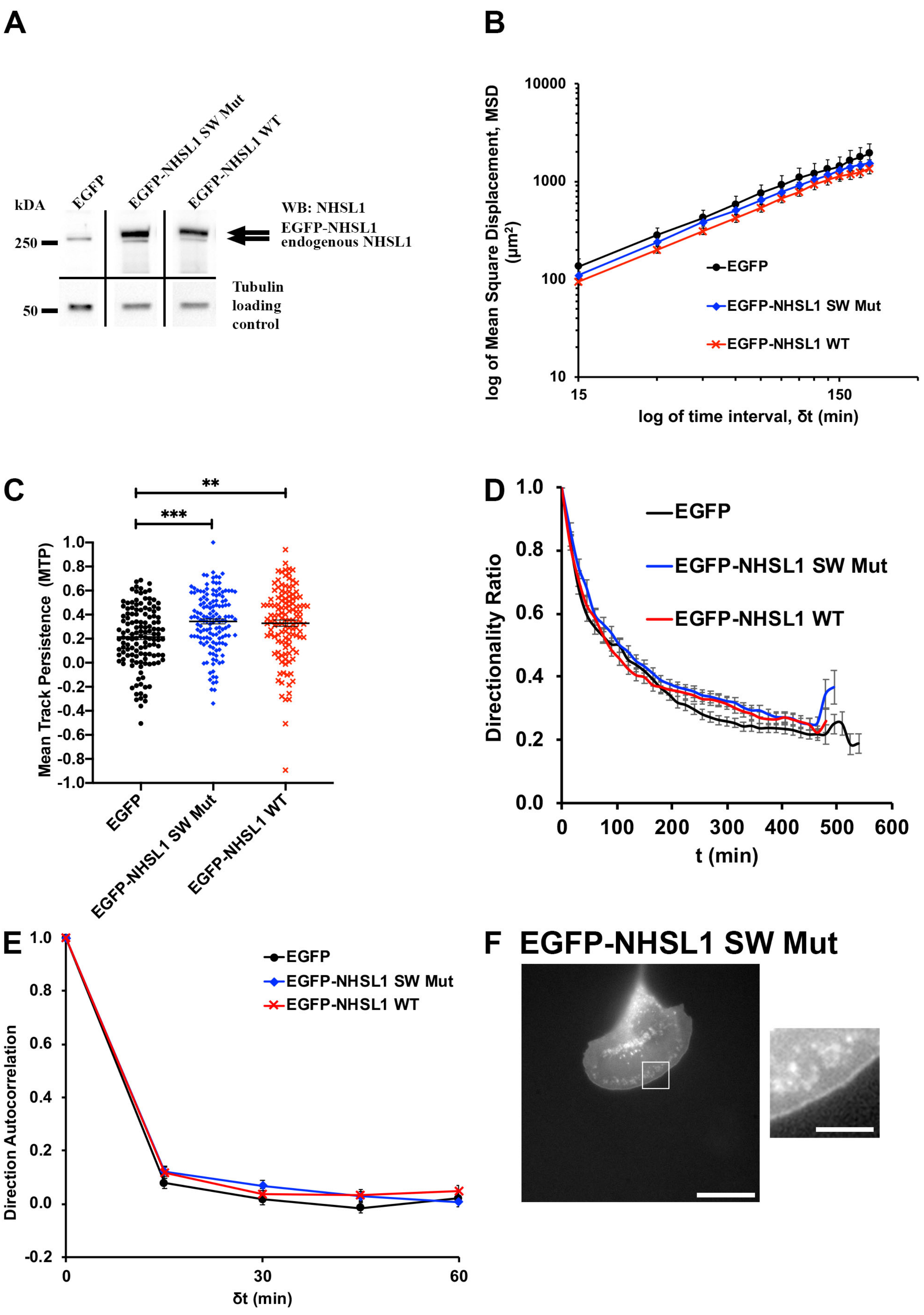
A
kDA
EGFP
EGFP-NHSL1 SW Mut
EGFP-NHSL1 WT
WB: NHSL1
EGFP-NHSL1
endogenous NHSL1
250
50
Tubulin
loading
control
B
log of Mean Square Displacement, MSD (μm²)
10000
1000
100
10
15
150
log of time interval, δt (min)
EGFP
EGFP-NHSL1 SW Mut
EGFP-NHSL1 WT
C
**
***
Mean Track Persistence (MTP)
1.0
0.8
0.6
0.4
0.2
0.0
-0.2
-0.4
-0.6
-0.8
-1.0
EGFP
EGFP-NHSL1 SW Mut
EGFP-NHSL1 WT
D
Directionality Ratio
EGFP
EGFP-NHSL1 SW Mut
EGFP-NHSL1 WT
t (min)
0
100
200
300
400
500
600
E
Direction Autocorrelation
EGFP
EGFP-NHSL1 SW Mut
EGFP-NHSL1 WT
δt (min)
0
30
60
F
EGFP-NHSL1 SW Mut

### Figure S11

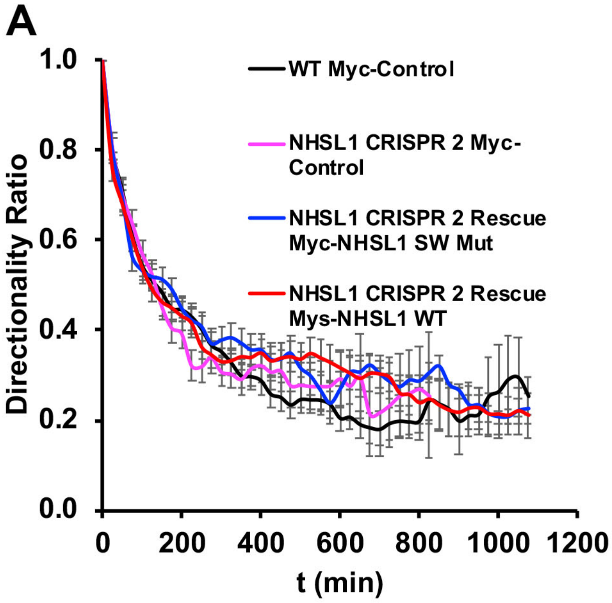
A
1.0
0.8
0.6
0.4
0.2
0.0
Directionality Ratio
0
200
400
600
800
1000
1200
t (min)
WT Myc-Control
NHSL1 CRISPR 2 Myc-Control
NHSL1 CRISPR 2 Rescue Myc-NHSL1 SW Mut
NHSL1 CRISPR 2 Rescue Mys-NHSL1 WT

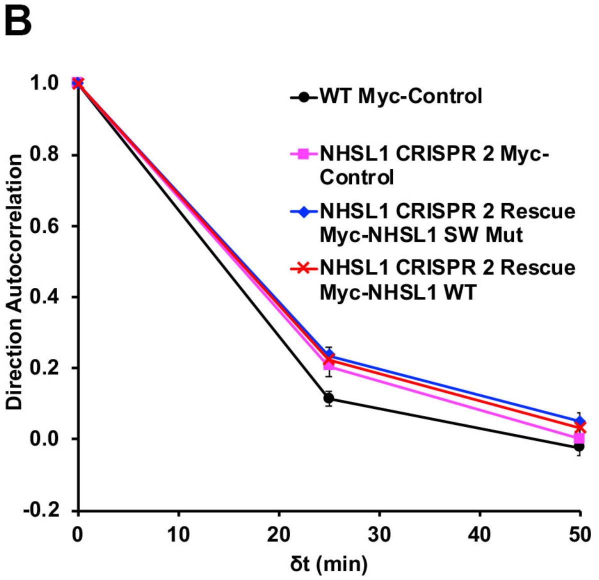
B
1.0
0.8
0.6
0.4
0.2
0.0
-0.2
Direction Autocorrelation
0
10
20
30
40
50
δt (min)
WT Myc-Control
NHSL1 CRISPR 2 Myc-Control
NHSL1 CRISPR 2 Rescue Myc-NHSL1 SW Mut
NHSL1 CRISPR 2 Rescue Myc-NHSL1 WT

### Figure S12

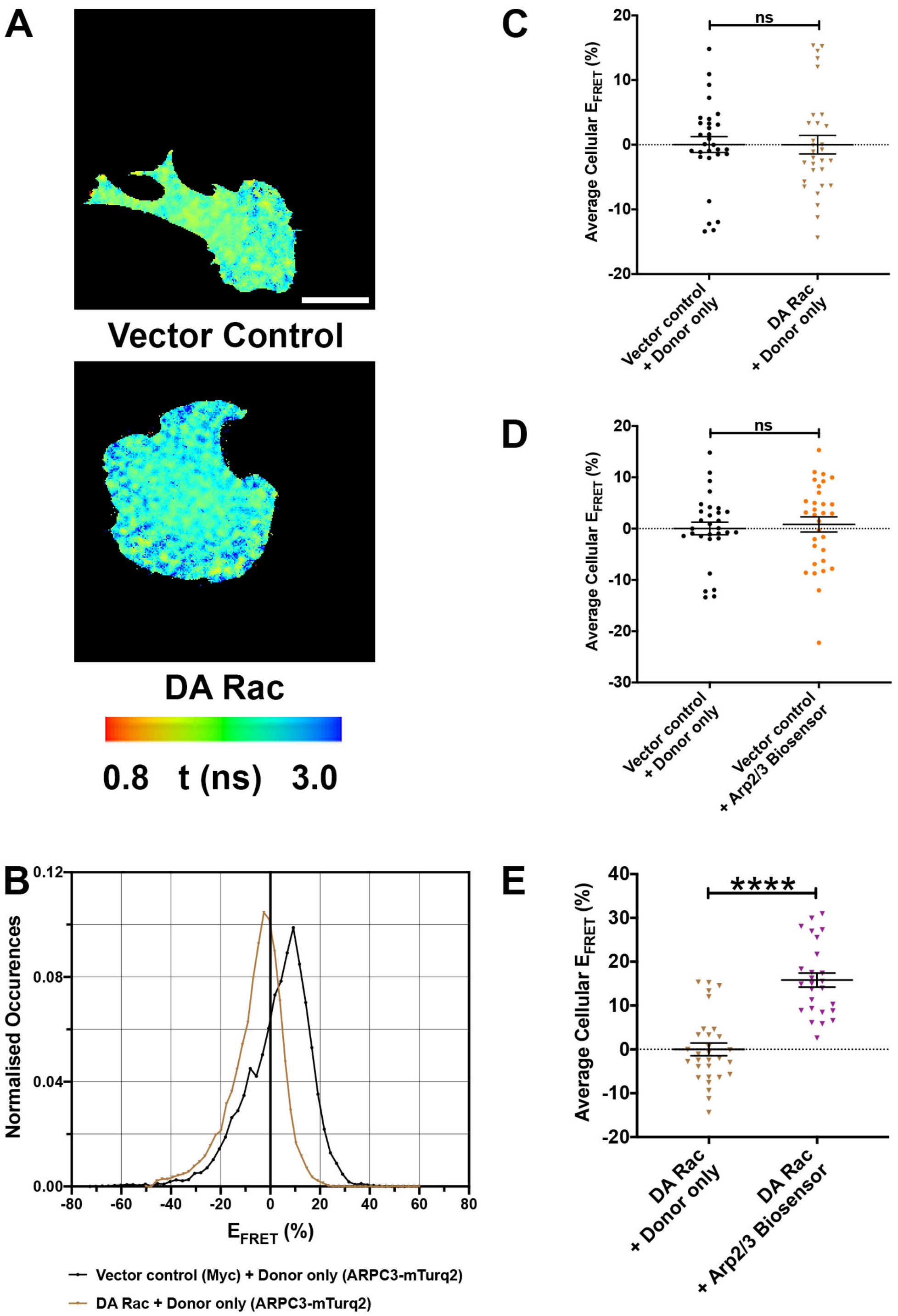

A
Vector Control
DA Rac
0.8 t (ns) 3.0
B
Normalised Occurences
0.12
0.08
0.04
0.00
-80 -60 -40 -20 0 20 40 60 80
$E_{FRET}$ (%)
Vector control (Myc) + Donor only (ARPC3-mTurq2)
DA Rac + Donor only (ARPC3-mTurq2)
C
ns
Average Cellular $E_{FRET}$ (%)
20
10
0
-10
-20
Vector control + Donor only
DA Rac + Donor only
D
ns
Average Cellular $E_{FRET}$ (%)
20
10
0
-10
-20
-30
Vector control + Donor only
Vector control + Arp2/3 Biosensor
E
****
Average Cellular $E_{FRET}$ (%)
40
30
20
10
0
-10
-20
DA Rac + Donor only
DA Rac + Arp2/3 Biosensor


### Figure S13

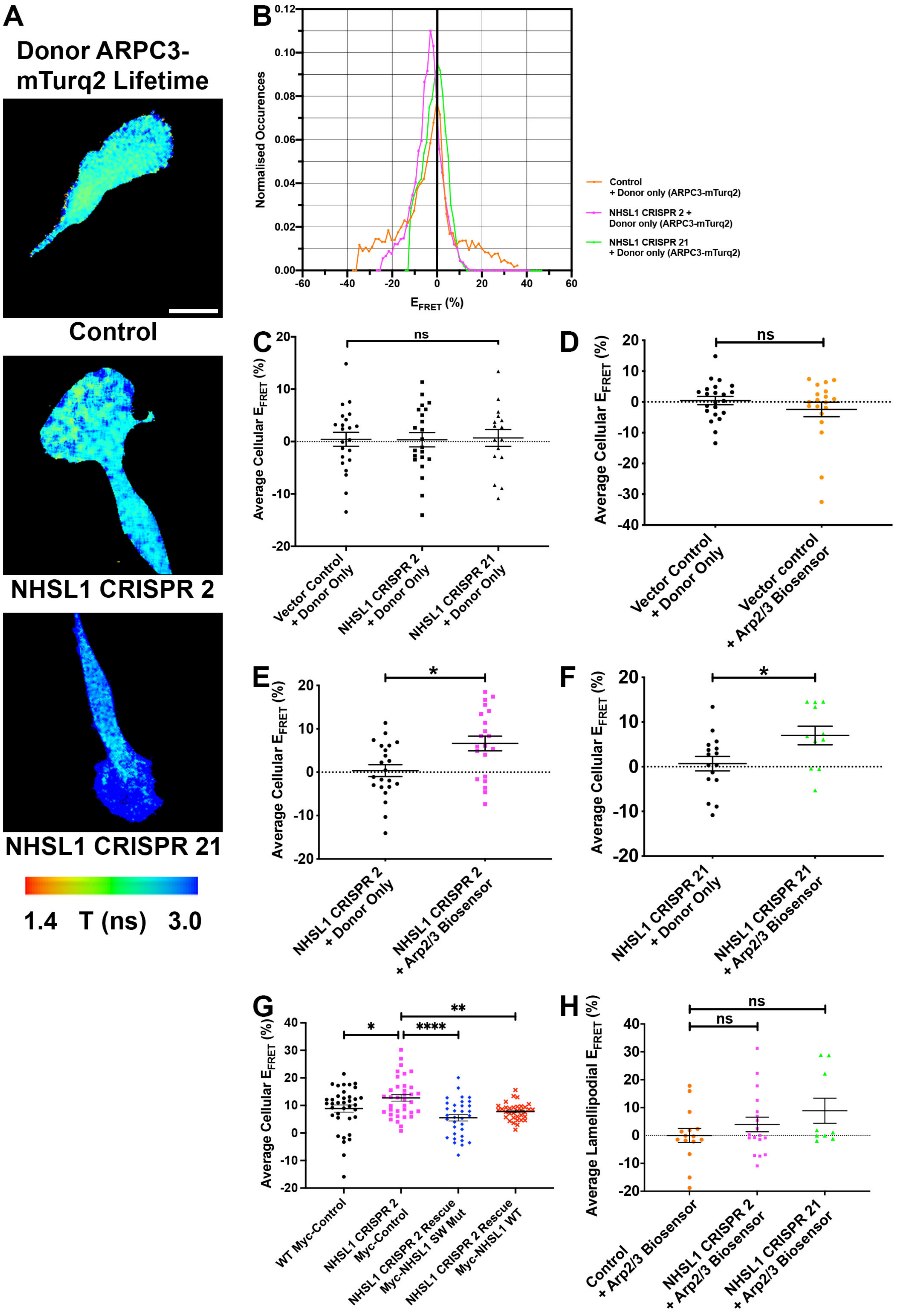
A
Donor ARPC3-mTurq2 Lifetime
Control
NHSL1 CRISPR 2
NHSL1 CRISPR 21
1.4 T (ns) 3.0
B
Normalised Occurences
EFRET (%)
Control + Donor only (ARPC3-mTurq2)
NHSL1 CRISPR 2 + Donor only (ARPC3-mTurq2)
NHSL1 CRISPR 21 + Donor only (ARPC3-mTurq2)
C
Average Cellular EFRET (%)
ns
Vector Control + Donor Only
NHSL1 CRISPR 2 + Donor Only
NHSL1 CRISPR 21 + Donor Only
D
Average Cellular EFRET (%)
ns
Vector Control + Donor Only
Vector control + Arp2/3 Biosensor
E
Average Cellular EFRET (%)
*
NHSL1 CRISPR 2 + Donor Only
NHSL1 CRISPR 2 + Arp2/3 Biosensor
F
Average Cellular EFRET (%)
*
NHSL1 CRISPR 21 + Donor Only
NHSL1 CRISPR 21 + Arp2/3 Biosensor
G
Average Cellular EFRET (%)
*
****
**
WT Myc-Control
NHSL1 CRISPR 2 Myc-Control
NHSL1 CRISPR 2 Rescue Myc-NHSL1 SW Mut
NHSL1 CRISPR 2 Rescue Myc-NHSL1 WT
H
Average Lamellipodial EFRET (%)
ns
ns
Control + Arp2/3 Biosensor
NHSL1 CRISPR 2 + Arp2/3 Biosensor
NHSL1 CRISPR 21 + Arp2/3 Biosensor

Figure S14

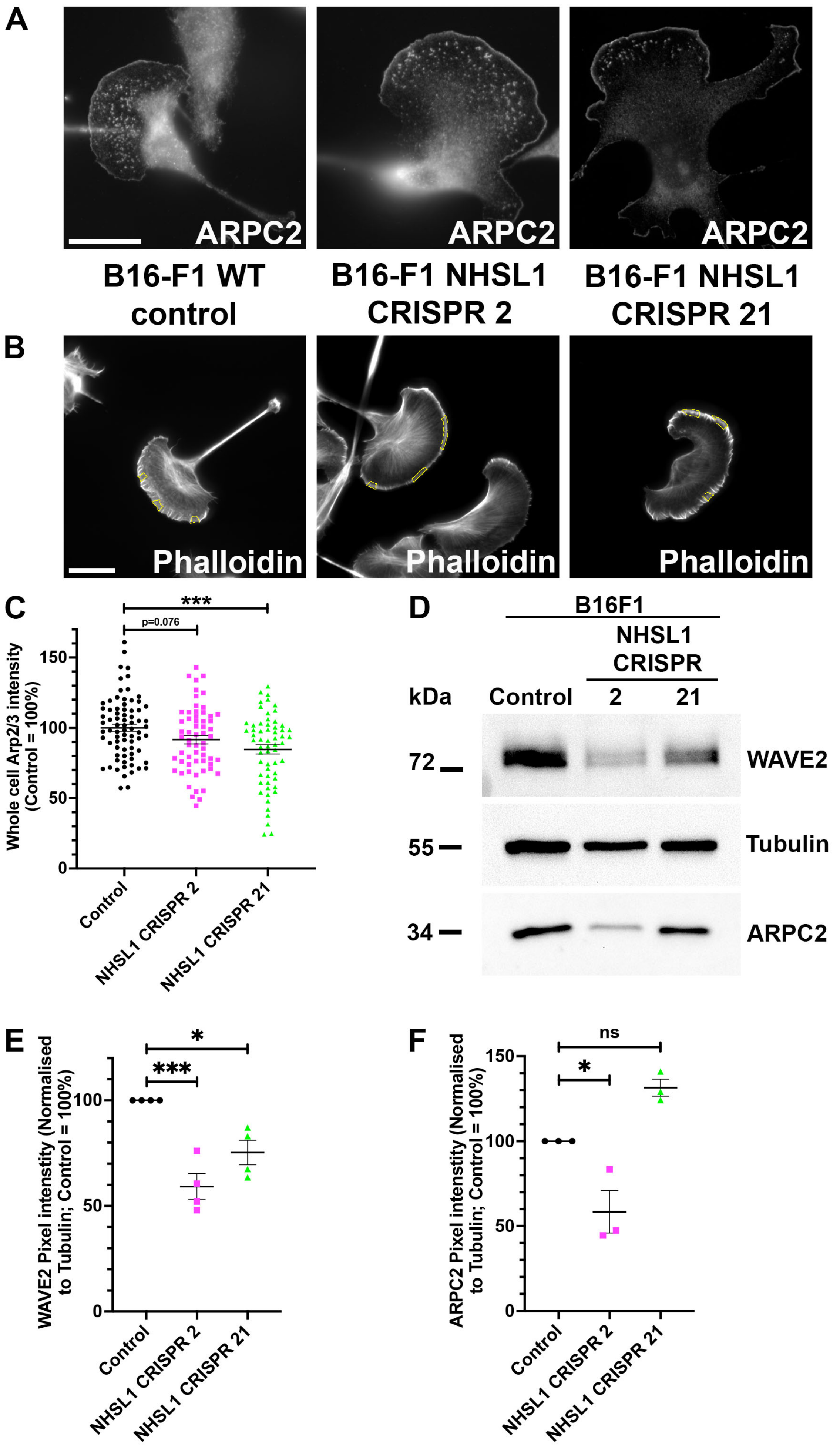

A
ARPC2
ARPC2
ARPC2
B16-F1 WT control
B16-F1 NHSL1 CRISPR 2
B16-F1 NHSL1 CRISPR 21
B
Phalloidin
Phalloidin
Phalloidin
C
***
p=0.076
Whole cell Arp2/3 intensity (Control = 100%)
150
100
50
0
Control
NHSL1 CRISPR 2
NHSL1 CRISPR 21
D
B16F1
NHSL1 CRISPR
kDa
Control
2
21
72
55
34
WAVE2
Tubulin
ARPC2
E
*
***
WAVE2 Pixel intensity (Normalised to Tubulin; Control = 100%)
100
50
0
Control
NHSL1 CRISPR 2
NHSL1 CRISPR 21
F
ns
*
ARPC2 Pixel intensity (Normalised to Tubulin; Control = 100%)
150
100
50
0
Control
NHSL1 CRISPR 2
NHSL1 CRISPR 21


### Figure S15

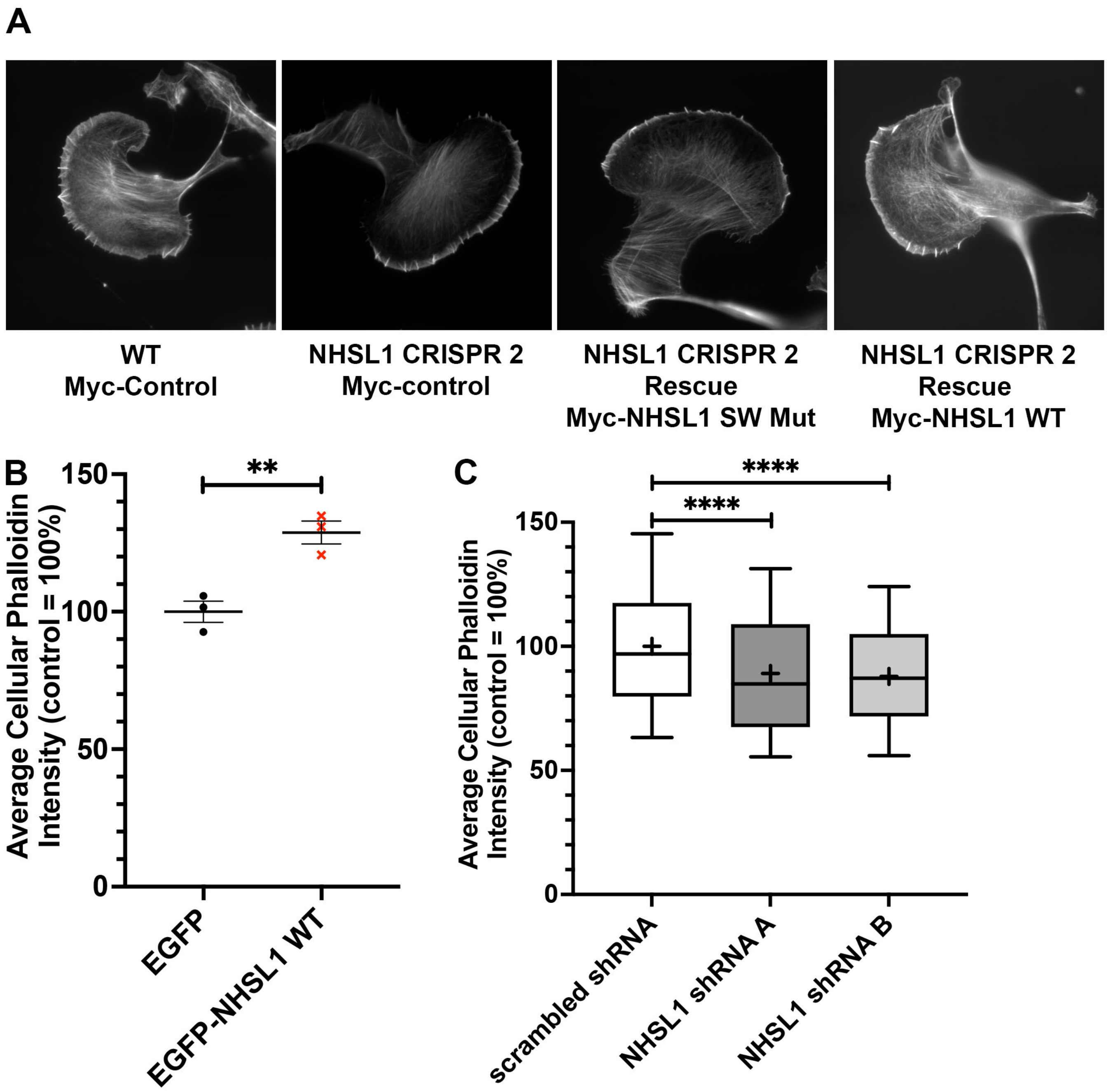
A
WT
Myc-Control
NHSL1 CRISPR 2
Myc-control
NHSL1 CRISPR 2
Rescue
Myc-NHSL1 SW Mut
NHSL1 CRISPR 2
Rescue
Myc-NHSL1 WT
B
Average Cellular Phalloidin
Intensity (control = 100%)
150
100
50
0
**
EGFP
EGFP-NHSL1 WT
C
Average Cellular Phalloidin
Intensity (control = 100%)
150
100
50
0
****
****
scrambled shRNA
NHSL1 shRNA A
NHSL1 shRNA B

### Figure S16

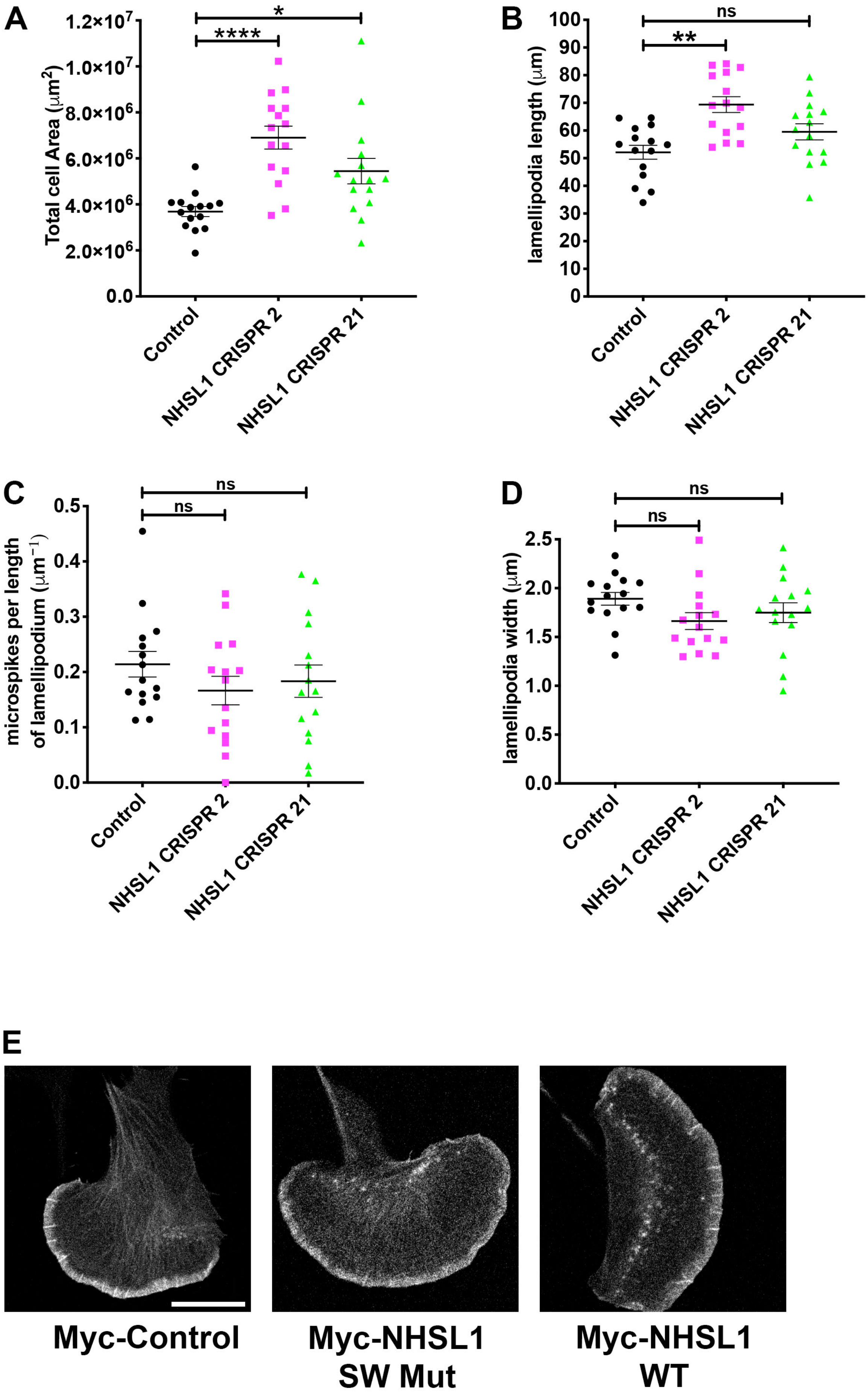
A
Total cell Area (μm²)
1.2×10⁷
1.0×10⁷
8.0×10⁶
6.0×10⁶
4.0×10⁶
2.0×10⁶
0.0
*
****
Control
NHSL1 CRISPR 2
NHSL1 CRISPR 21
B
lamellipodia length (μm)
100
90
80
70
60
50
40
30
20
10
0
ns
**
Control
NHSL1 CRISPR 2
NHSL1 CRISPR 21
C
microspikes per length of lamellipodium (μm⁻¹)
0.5
0.4
0.3
0.2
0.1
0.0
ns
ns
Control
NHSL1 CRISPR 2
NHSL1 CRISPR 21
D
lamellipodia width (μm)
2.5
2.0
1.5
1.0
0.5
0.0
ns
ns
Control
NHSL1 CRISPR 2
NHSL1 CRISPR 21
E
Myc-Control
Myc-NHSL1 SW Mut
Myc-NHSL1 WT

### Figure S17

A

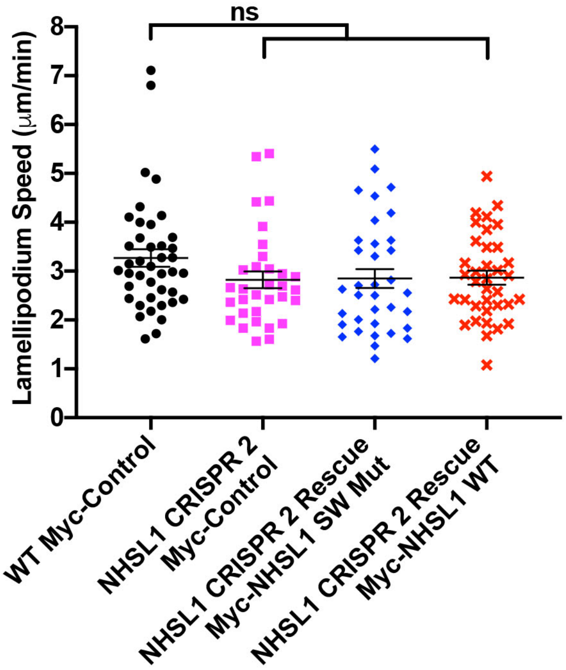


B

| % AA identity | hsNHS-Uniprot-Q6T4R5 | hsNHSL1-Uniprot-Q5SYE7 | hsNHSL2-Uniprot-Q5HYW2 |
|---|---|---|---|
| hsNHS-Uniprot-Q6T4R5 | 100.0 | 30.3 | 31.7 |
| hsNHSL1-Uniprot-Q5SYE7 | 30.3 | 100.0 | 25.1 |
| hsNHSL2-Uniprot-Q5HYW2 | 31.7 | 25.1 | 100.0 |

C **NHSL1 Abi binding site 2**

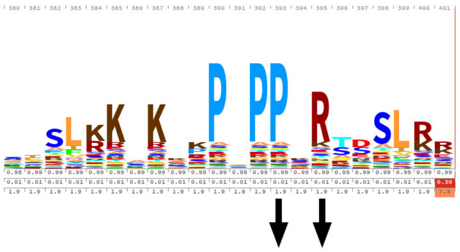

Mutated: A A

D **NHSL1 Abi binding site 3**

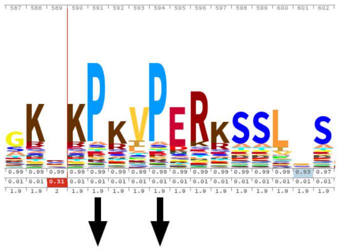

Mutated: A A

### Figure S18
